# Reimagining Gene-Environment Interaction Analysis for Human Complex Traits

**DOI:** 10.1101/2022.12.11.519973

**Authors:** Jiacheng Miao, Gefei Song, Yixuan Wu, Jiaxin Hu, Yuchang Wu, Shubhashrita Basu, James S. Andrews, Katherine Schaumberg, Jason M. Fletcher, Lauren L. Schmitz, Qiongshi Lu

## Abstract

In this study, we introduce PIGEON—a novel statistical framework for quantifying and estimating polygenic gene-environment interaction (GxE) using a variance component analytical approach. Based on PIGEON, we outline the main objectives in GxE studies, demonstrate the flaws in existing GxE approaches, and introduce an innovative estimation procedure which only requires summary statistics as input. We demonstrate the statistical superiority of PIGEON through extensive theoretical and empirical analyses and showcase its performance in multiple analytic settings, including a quasi-experimental GxE study of health outcomes, gene-by-sex interaction for 530 traits, and gene-by-treatment interaction in a randomized clinical trial. Our results show that PIGEON provides an innovative solution to many long-standing challenges in GxE inference and may fundamentally reshape analytical strategies in future GxE studies.

## Introduction

The environment is often ignored or treated as a nuisance in human complex trait genetics research. However, in epidemiology, social sciences, medicine, and many other related disciplines, there is a great interest in quantifying the effect heterogeneity of an exposure (e.g., a treatment, a policy change, a natural experiment), and, more specifically, its interaction with genetics, as a means of identifying subgroups that may maximally shift in response to environmental instruments^1-5^. This is broadly referred to as gene-environment interaction (GxE)^6^. While this concept seems intuitive, GxE has not been consistently defined in the literature, especially for complex traits due to their polygenic nature^7^. Perhaps more importantly, many existing complex trait GxE methods lack a solid statistical foundation. It is often unclear how to compare different GxE approaches or even whether they can be compared, since they may be estimating entirely different parameters. We will illustrate this issue throughout the current paper. These issues have held back a broader consensus of findings in the GxE field, for which we propose a solution in this study. Here, we aim to achieve two main goals. First, we introduce a unified statistical framework to model polygenic GxE effects for complex traits, which allows us to define the parameters of interest and compare existing GxE approaches. Second, we introduce an innovative approach to estimating GxE interactions using genome-wide summary data.

The evolution of GxE methodology mirrors method development in genome-wide association studies (GWAS). Early G×E studies were primarily based on a candidate gene approach^8^ which suffered from low replicability^9^. High-throughput genotype data made it possible to perform GxE scans for millions of single nucleotide polymorphisms (SNPs)^10^, a design often referred to as genome-wide interaction study (GWIS). Although GWIS improves the replicability and robustness of interaction findings, it introduces an extreme burden of multiple testing which severely limits its statistical power^11^. Therefore, a two-step approach is sometimes employed to first filter SNPs (e.g., based on GWAS associations) and then only test GxE using selected SNPs^12-16^.

However, it is now well established that most human complex traits are highly polygenic^7^. As a result, modern GWAS analyses have generally focused less on individual SNPs, but instead employ tools that embrace polygenicity, including genome-wide heritability estimation^17,18^ and enrichment analysis^19,20^, genetic correlation analysis which quantifies shared genetics across multiple traits^21,22^, and polygenic scores (PGS) which estimate genetic predisposition by aggregating effects of many SNPs^22-25^. GxE studies are going through a similar transition, focusing more on how the polygenic basis of a trait varies across environments^26-29^. For example, some studies estimate the phenotypic variance explained by many SNPxE interaction terms using similar methods from heritability estimation^28-31^. Other studies perform stratified GWAS in different environments and then test differential heritability and/or imperfect genetic correlation between the environments^26,27,32^. Further, PGSxE studies have gained popularity in the GxE literature^3,33-35^. It is a two-step approach that first summarizes each individual’s genetic predisposition into a PGS, and then tests the interaction between PGS and the environment^1,36-38^. Although all these approaches are referred to as GxE in the literature, the relationship between these approaches is poorly understood. For example, it is unclear whether PGSxE and differential heritability analysis aim to estimate the same parameter (as we explain below, they do not). Existing methods are also plagued by technical challenges including statistical biases, computational burden, and constraints in the data. We will provide detailed discussions of these issues in the following sections.

In this paper, we introduce a statistical framework named **p**olygen**i**c **g**ene-**e**nvironment interacti**on** (PIGEON) for quantifying and estimating polygenic GxE. Using this framework, we demonstrate the relation and differences between existing GxE methods. We also equip PIGEON with an estimation method only requiring GWIS and GWAS summary statistics as inputs. Our method provides unbiased estimates, is robust to sample overlap and heteroskedasticity, and allows for hypothesis-free scans for PGSxE across many PGS. We demonstrate PIGEON’s superior performance over existing methods through extensive theoretical analysis, simulation studies, and real data applications. In this study, we pursue three main applications, all of which leverage genome-wide data and exogenous environmental exposures, to showcase the broad applications of PIGEON. We validated our approach by replicating a quasi-experimental PGS-by-education interaction (PGSxEdu) study for health-related outcomes in the UK Biobank (UKB)^1^. We then used PIGEON to build a catalog of polygenic gene-by-sex interactions (GxSex) for 530 traits in UKB. We further applied PIGEON to investigate the effect heterogeneity of smoking cessation treatment in a randomized clinical trial.

## Results

### Two main objectives in polygenic GxE inference

The PIGEON model is illustrated in **Figure 1**. It is built on a linear mixed model that captures both the additive effects and GxE effects for many SNPs.

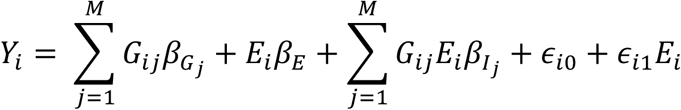

Here, *Y*_*i*_ is the standardized phenotype with a mean of 0 and variance of 1 for the i-th individual, *G*_*ij*_ is the j-th standardized SNP, *E*_*i*_ is the standardized environment, *ϵ*_*i*0_ is the noise term, and *ϵ*_*i*1_*E* quantifies the heteroskedasticity due to residual-environment interaction (i.e., varying residual variance across environments)^30,39^. Polygenic additive effects and interaction effects (i.e., 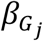 and 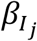) are modeled as random variables. Details on this model and its more generalized forms are presented in **Methods** and **Supplementary Note 1**.

**Figure 1.**
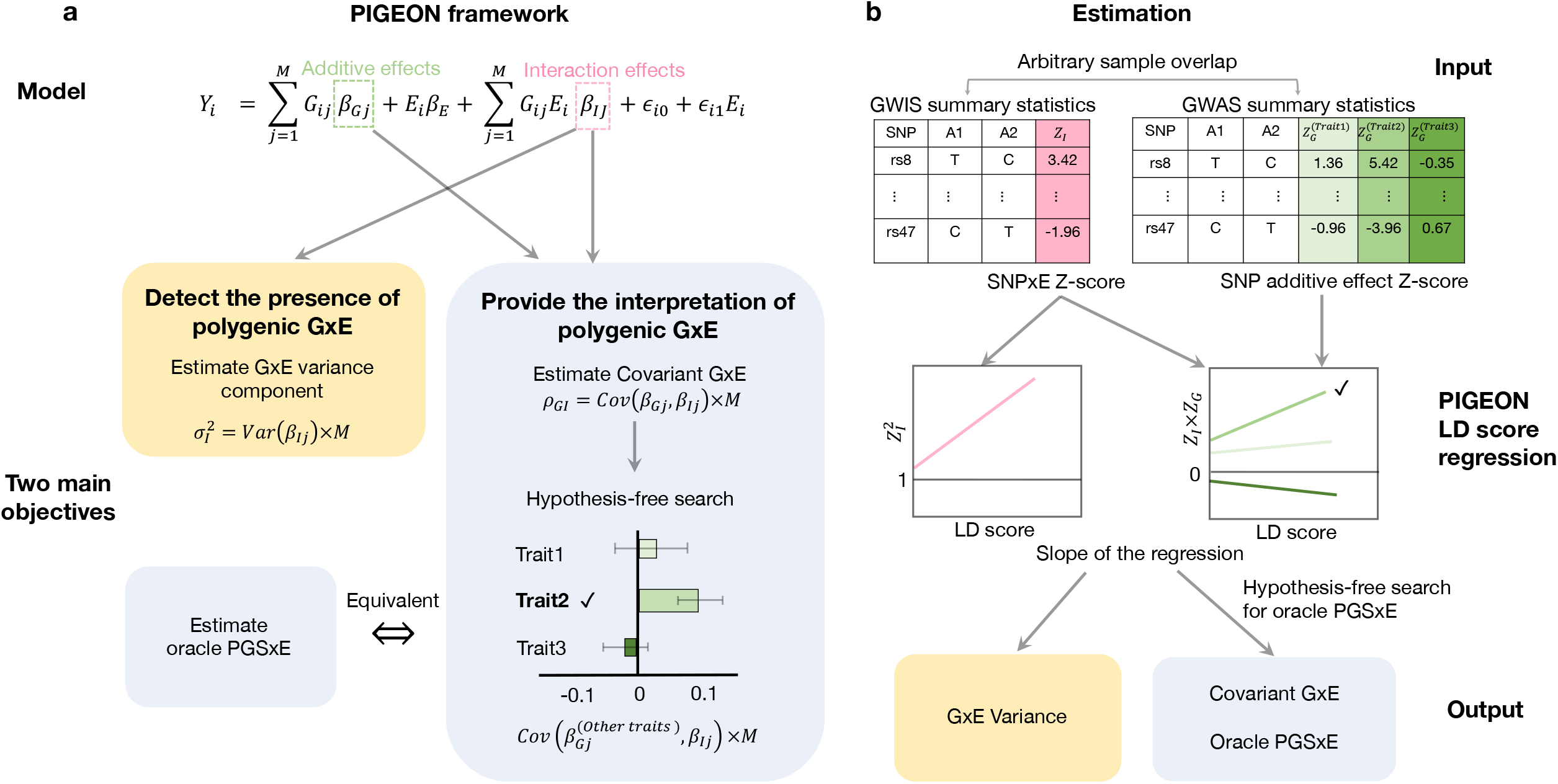
PIGEON workflow. **(a)** Two main objectives in polygenic GxE inference based on the PIGEON model **(b)** Estimating polygenic GxE using GWIS and GWAS summary statistics.

PIGEON defines two main objectives in polygenic GxE inference using a variance component analysis framework. First, the overall GxE contribution is quantified by the variance of interaction effects (i.e.,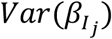) and the proportion of the phenotypic variance attributed to it. This is conceptually similar to SNP heritability^23^ but focuses on interaction effects rather than additive effects. Hypothesis testing on this quantity provides evidence for the existence of GxE. Its magnitude quantifies the degree of GxE for the trait of interest.

However, a non-zero GxE variance component alone does not provide much mechanistic insight. Therefore, we propose another key objective in polygenic GxE analysis– estimating covariant GxE, defined as the covariance between SNP additive effects and SNPxE interaction effects (i.e.,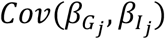). We note that additive effects and interaction effects can be obtained from the same trait or two different traits (**Methods**). Here, covariant GxE provides crucial insights into the whole-genome interaction mechanisms by correlating SNPs’ effects on complex traits with their tendency to interact with the environment. This is analogous to genetic correlation analysis in the GWAS literature where researchers use existing GWAS to help interpret genetic associations obtained in a new GWAS.

Together, these two objectives lay out the foundation for (1) quantifying the evidence for polygenic GxE interaction and (2) interpreting the mechanisms underlying these interactions. In the next section, we demonstrate that existing GxE approaches, i.e., GxE variance estimation, differential heritability analysis between environments, genetic correlation analysis between environments, and PGSxE analysis can all be linked to these two objectives, which allows us to understand the connection and distinction between these approaches.

### Comparing polygenic GxE methods in the PIGEON framework

Next, we consolidate and compare several GxE methods under the PIGEON framework and demonstrate the advantages of variance-covariance component analysis for polygenic GxE. We present the statistical details and technical discussions in **Methods** and **Supplementary Note 2**. For illustration, we assume G-E independence (i.e., the environment has zero heritability), but later we will relax this assumption and investigate how the correlation between genes and environments affect polygenic GxE inference. **Table 1** provides a summary of the comparison between PIGEON and other approaches.

**Table 1.**
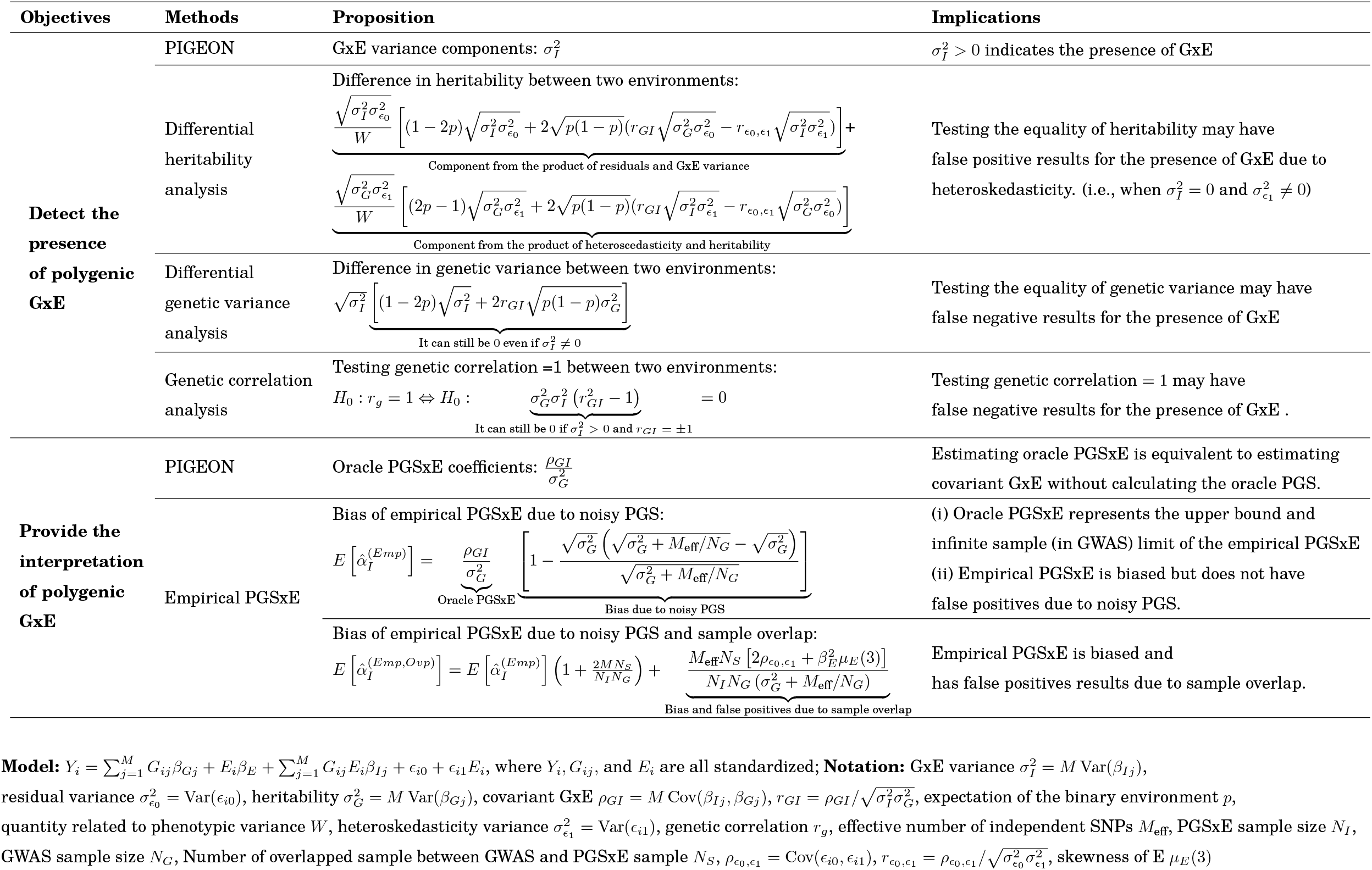
Summary of the theoretical results of PIGEON and other approaches for polygenic GxE inference.

To perform statistical inference on the presence of GxE, we can test the null hypothesis that no SNPs have interactions with the environment, i.e., *H*_0_: *β*_*Ij*_ = 0 for all j. However, the high dimensionality in GWAS data creates a challenge. In PIGEON, the same null hypothesis can be specified as having a zero GxE variance, i.e., 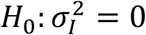, which only requires estimating one parameter^28-31^. Here, 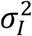 is the total variance of SNPxE effects in the genome, i.e., 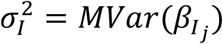. Several commonly-used approaches provide flawed estimates of this quantity. For example, heritability could vary across discrete environments just because of heteroscedasticity (i.e., difference in the non-genetic variance components) in the absence of GxE, leading to false positive results (**Table 1**). Notably, comparing genetic variance instead of heritability^27^ does not solve the issue either – genetic variance may be the same across environments in the presence of GxE, leading to false negative results (**Table 1**). Similarly, genetic correlation analysis between environments has its limitations. A perfect genetic correlation can be achieved when the SNP additive effects are proportional between environments (**Table 1**; also known as “amplification” in the GxE literature^40^), leading to false negative results. Even testing both genetic variance and perfect genetic correlation between environments may fail to identify GxE (**Supplementary Note 2**). Therefore, heritability, genetic variance, and genetic correlation analyses fail to properly estimate polygenic GxE effects. Some of these approaches also suffer from technical issues such as an inability to handle quantitative environmental exposures. To assess the presence or absence of polygenic GxE, researchers should estimate the GxE variance component.

In PGSxE analysis, we are interested in quantifying the interaction between the environment and each individual’s true PGS (i.e., PGS computed from each SNP’s true effect size). We refer to this as oracle PGSxE analysis. We show that estimating oracle PGSxE is equivalent to estimating covariant GxE (**Supplementary Note 2**). This finding has several major implications. It shows that oracle PGSxE coefficient can be estimated without calculating PGS. Instead, we could equivalently estimate the covariance between polygenic additive and interaction effects. It also suggests that although PGSxE analysis and several other approaches we have discussed above (e.g., differential heritability) are all believed to estimate “GxE”, they in fact quantify two different objectives in GxE discussed above. A null PGSxE, which could simply result from uncorrelated SNP additive and SNPxE interaction effects, does not imply the absence of GxE. In addition, we refer to the PGSxE analysis based on scores estimated from GWAS as empirical PGSxE. This analytical strategy is substantially affected by the imprecision in PGS estimation due to limited sample sizes in GWAS^41^. Ignoring the uncertainty in empirical PGS will lead to interaction estimates that are biased towards zero (**Table 1** and **Supplementary Note 3**). Oracle PGSxE represents the upper bound and infinite sample (in GWAS) limit of the empirical PGSxE (**Methods**), analogous to the heritability being the upper bound of PGS predictive R-squared in the GWAS literature^42^. It is also important to note that empirical PGSxE analysis requires no overlap between the GWAS used to construct PGS and the sample for PGSxE analysis. Under sample overlap, PGS will overfit and cause biased interaction estimates and false findings in empirical PGSxE analysis (**Table 1**). Therefore, estimating covariant GxE through variance component analysis is a superior alternative to replace the commonly used PGSxE analysis.

### Estimating GxE using GWIS and GWAS summary statistics

Next, we introduce PIGEON linkage disequilibrium (LD) score regression (PIGEON-LDSC) to estimate GxE variance and covariant GxE using only summary statistics from GWAS and GWIS (**Figure 1b**)^21^. For the GxE variance component estimation, PIGEON-LDSC regresses the squared SNPxE Z-scores from GWIS summary statistics on LD scores, and estimates the GxE variance parameter from the regression slope^31^. To estimate covariant GxE, PIGEON-LDSC uses summary statistics from both GWAS and GWIS, and regresses the product of GWAS and GWIS Z-scores on LD scores. Oracle PGSxE effect size can be subsequently obtained from normalizing covariant GxE by trait heritability (**Methods**).

Importantly, the sample overlap between GWAS and GWIS will only affect the intercept of the regression but not the slope. Therefore, PIGEON-LDSC’s interaction effect estimates are robust to sample overlap. Because of this, a key feature of our estimation framework is the ability to implement hypothesis-free scans for PGSxE across many PGS. That is, given an environmental exposure of interest, we can search through many published GWAS to identify the PGS that modifies the exposure effect – we simply need to estimate the genetic correlation between the GWIS summary data and publicly available GWAS summary statistics for many traits. In addition, PIGEON is computationally efficient, robust to the heteroskedasticity across environments, able to quantify GxE interaction with dichotomized PGS, and can be used for binary outcomes. We provide details on these features in the **Supplementary Note 4-6**. We also present a comparison between PIGEON-LDSC and the empirical PGSxE approach in **Figure 2**.

**Figure 2.**
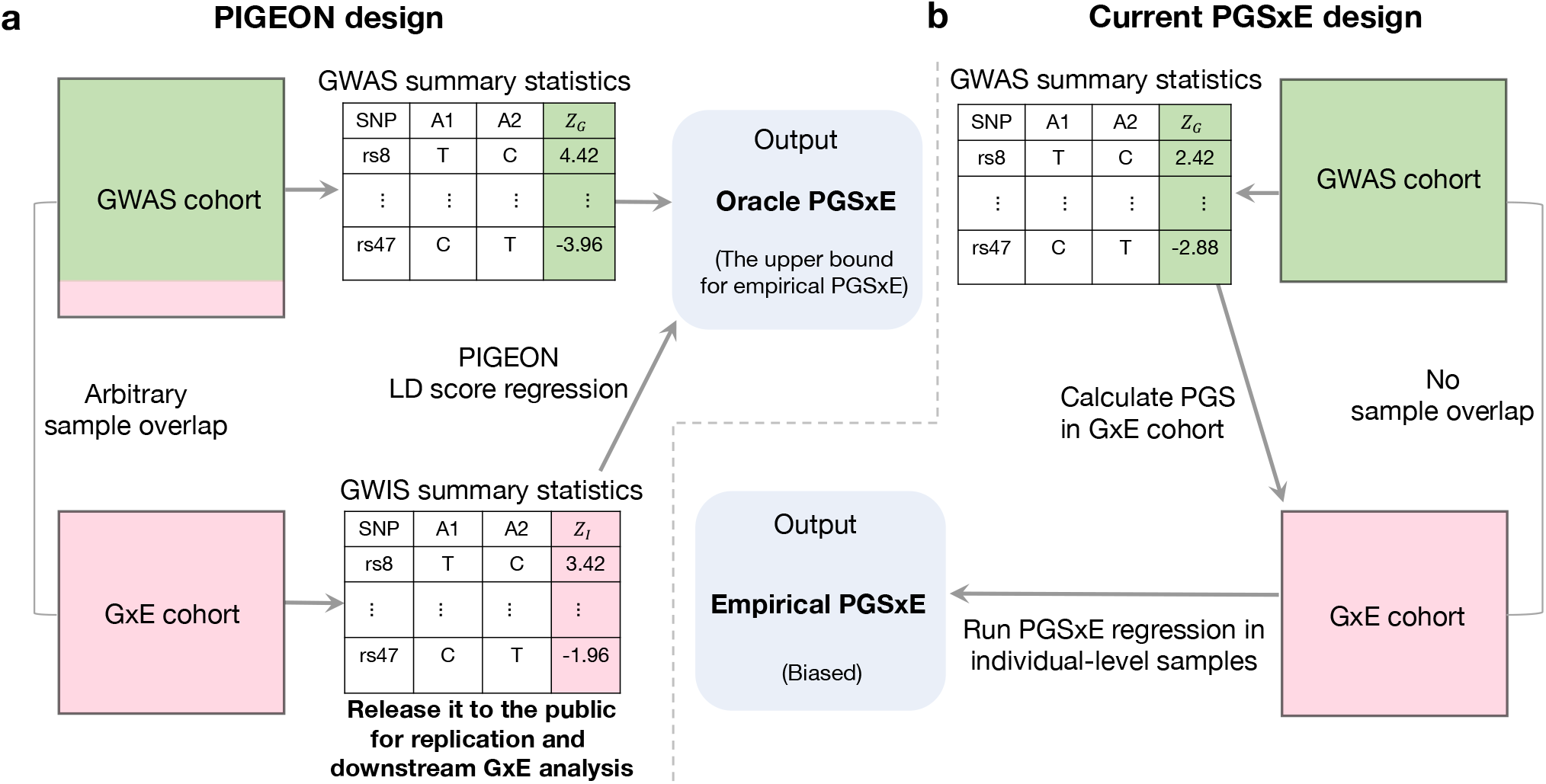
Comparison of PIGEON and PGSxE analysis. **(a)** Estimating oracle PGSxE with PIGEON-LDSC. **(b)** Conventional study design for PGSxE analysis.

### Simulation results

We performed numerous simulations using genotype data from UKB^43^ to demonstrate the unbiasedness, power, and robustness of PIGEON inference results (**Methods**). We included 40,000 independent samples and 734,046 SNPs in the analysis after quality control (QC). Phenotypes were simulated under the polygenic GxE model with a binary environment variable. Each simulation was repeated 100 times.

We equally divided 40,000 samples into two sub-cohorts with 20,000 each. We performed GWIS on the first sub-cohort and applied PIGEON-LDSC to estimate the GxE variance component. We compared PIGEON with differential heritability and genetic correlation analyses. PIGEON showed higher power than both approaches (**Figure 3a** and **Supplementary Figure 1**) and provided unbiased estimates for GxE variance (**Figure 3d** and **Supplementary Figure 1**). We also compared PIGEON with GxEsum, another approach designed to estimate GxE variance component^31^, and demonstrate the bias in its implementation (**Supplementary Figure 2** and **Supplementary Note 7**).

**Figure 3.**
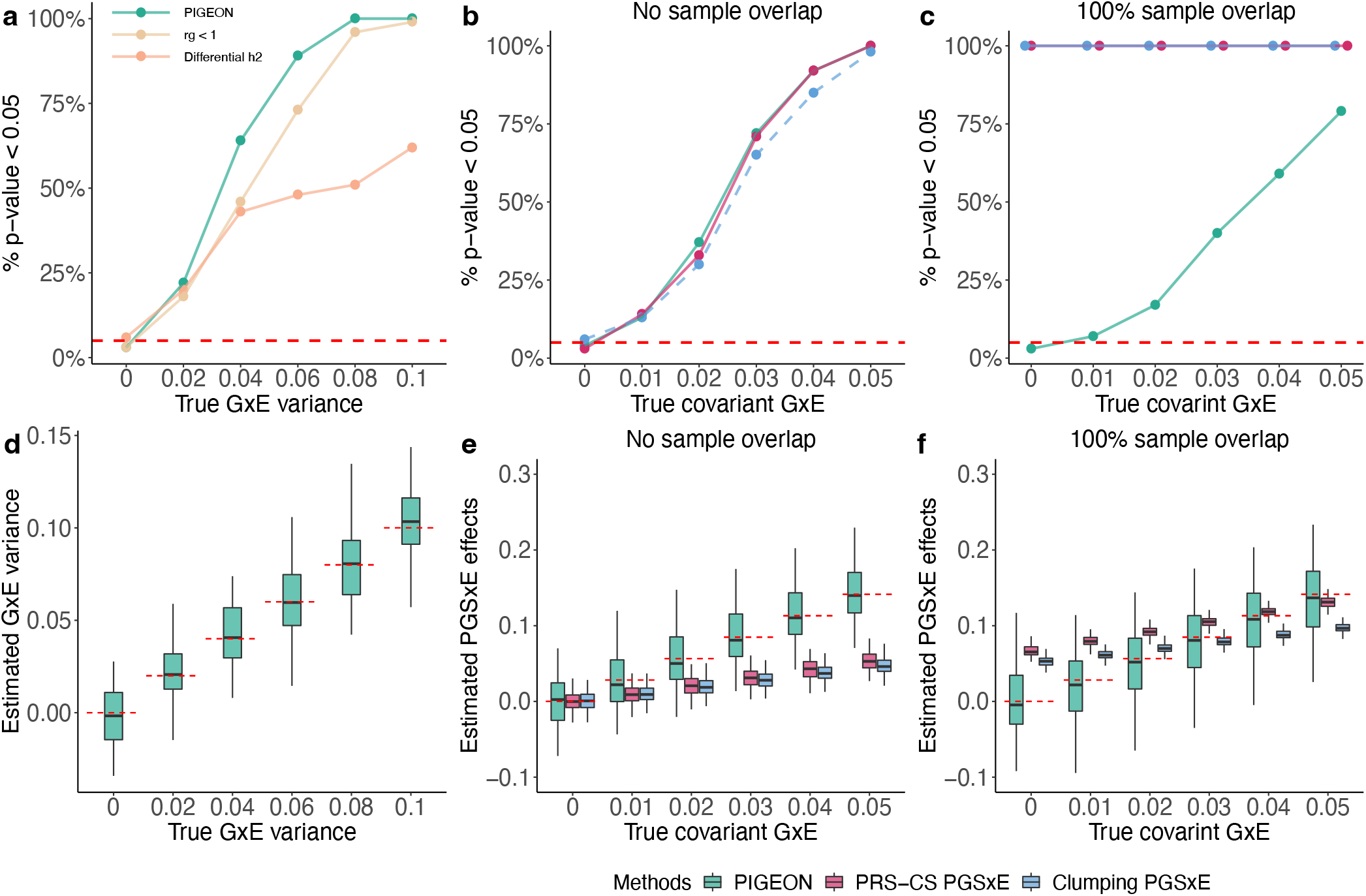
Simulation results. (**a**) Statistical power and type-I error of PIGEON GxE variance component estimation, differential heritability between environments, genetic correlation between environments. (**b**) Statistical power and type-I errors of PIGEON’s oracle PGSxE estimation and empirical PGSxE analysis based on clumping and PRS-CS scores. GWAS and GWIS have no sample overlap. (**c**) Same as **b** except that GWAS and GWIS share 100% of the samples. (**d**) Point estimation for GxE variance components using PIGEON (**e**) Point estimation for PGSxE coefficients when GWAS and GWIS have no sample overlap. (**f**) Same as **e** but with 100% sample overlap.

Next, we evaluated the performance of bivariate PIGEON-LDSC in estimating covariant GxE. We compared PIGEON with the empirical PGSxE approach based on two scores: clumping PRS^44^ and PRS-CS^45^. In the absence of sample overlap, where the GWAS and GWIS were performed in different sub-cohorts, no methods showed inflated type-I error. PIGEON showed similar power compared to PRS-CS PGSxE and slightly higher power than clumping PGS (**Figure 3b**). Importantly, PIGEON provided unbiased estimates for the oracle PGSxE, while both clumping and PRS-CS PGSxE results were severely biased (**Figure 3e**). When GWAS and GWIS were generated in the same sub-cohort with a full sample overlap, PGSxE approaches showed severe type-I error inflation and biased estimates while PIGEON estimates remained unbiased with well-controlled type-I error (**Figure 3c** and **f**). Altering the sample size ratio between GWIS and GWAS reached the same conclusions in simulations (**Supplementary Figure 3**). We also employed an approach to correct for measurement error in PGSxE analysis^46^. Measurement error correction led to both inflated type-I error and lower statistical power, showing inferior performance compared to PIGEON (**Supplementary Note 7** and **Supplementary Figure 4**). These simulation results are consistent with our theoretical analysis and demonstrate the superiority of PIGEON over commonly used GxE approaches.

### PGS-by-education interaction for health outcomes

To further compare PIGEON and the PGSxE approach, we replicated the analysis in Barcellos et al.^1^ to study whether genetics moderate the effect of an education reform on later life health-related outcomes. We focused on their most significant PGSxE findings for the dichotomous summary health index (**Methods**), but also replicated their null results as negative controls. We performed PIGEON and PGSxE to quantify the interaction between the effect of education reform, and genetic predisposition for body mass index (BMI)^47^ and educational attainment (EA)^48^. We used the same BMI and EA GWAS as Barcellos et al., which excluded UKB samples to avoid PGS overfitting.

Our results are summarized in **Table 2**. PIGEON showed similar P-values, but substantially elevated interaction effect estimates by 19.7%-55% compared to PGSxE analysis. This is consistent with our observation in simulations – empirical PGSxE analysis underestimates interaction effects due to measurement error in the constructed PGS measure. Both PIGEON and PGSxE found null results for the continuous health index. We also repeated the PIGEON analysis using two recently-published, larger GWAS for BMI^49^ and EA^50^ which included UKB samples. We obtained highly consistent results in this analysis, demonstrating PIGEON’s robustness to sample overlap. Further, we applied PIGEON to perform a hypothesis-free search for novel PGSxEdu interactions using GWAS summary statistics for 30 complex traits (**Supplementary Tables 1-2**). We found a significant interaction between the education reform and the genetic risk for smoking initiation^51^, suggesting that education is more effective in improving health for people with a higher genetic risk for smoking. We also validated this finding using the empirical PGSxE approach after removing UKB samples from the smoking initiation GWAS^51^ (**Supplementary Table 3**).

**Table 2.**
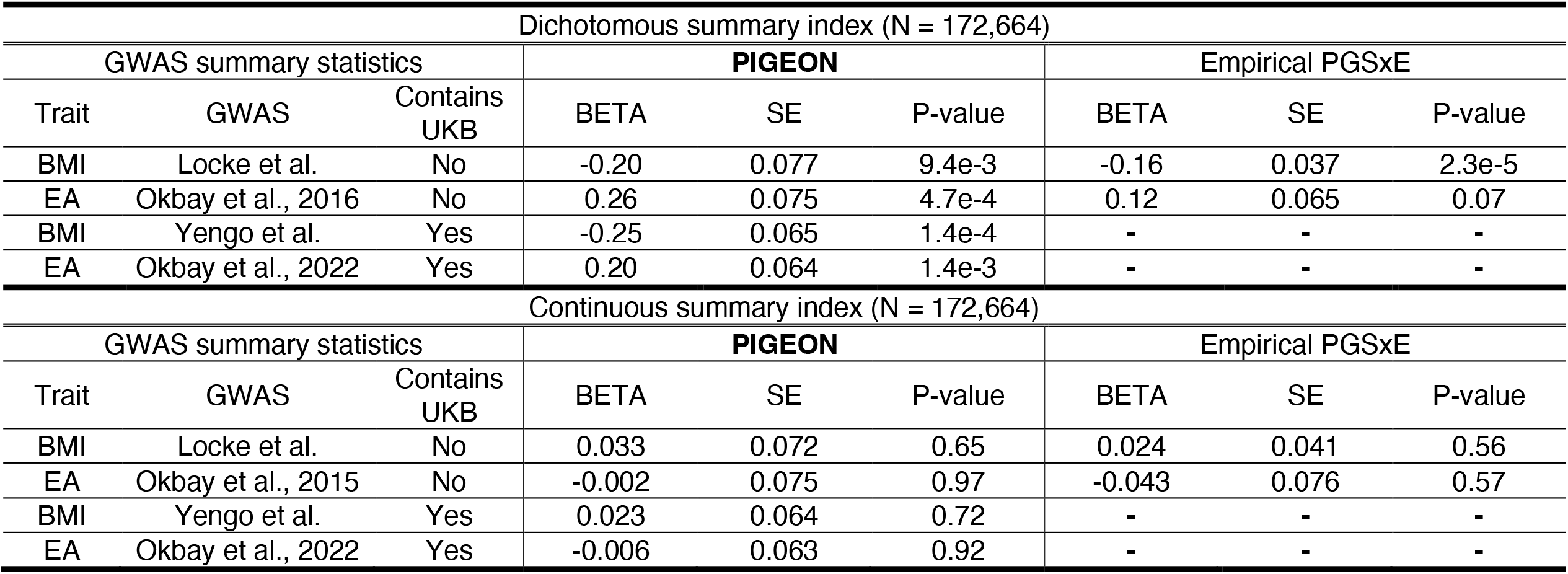
PGS x education effects on summary health index in UKB. The upper and lower table show the results for dichotomous and continuous summary indices, respectively. We compare the results of PIGEON and empirical PGSxE. Dash (-) means that analysis could not be performed due to the sample overlap between GWAS and GxE cohorts. Abbreviations are as follows: BMI, body mass index; EA, educational attainment measured by years of education; SE, standard error.

### Polygenic GxSex interaction for 530 complex traits

Next, we deployed PIGEON to create an atlas of polygenic GxSex interactions for 530 complex traits in UKB. We obtained GWIS summary statistics for 530 traits by transforming the sex-stratified GWAS summary statistics traits released by Bernabeu et al.^27^ (**Supplementary Table 4**), and used them as input for PIGEON-LDSC (**Methods**). We estimated the proportion of phenotypic variance attributed to GxSex and searched for covariant GxSex (or equivalently, oracle PGSxSex interactions) on these traits using 30 GWAS (**Supplementary Table 1**). We also estimated additive effect genetic correlations^21^ between 530 UKB traits and 30 GWAS to help interpret the sign of polygenic GxSex interactions (**Methods**).

PIGEON identified 64 traits with significant GxSex variance components (P < 9.4e-5 using Bonferroni correction). As a comparison, analysis based on differential heritability and genetic correlation between the sexes identified substantially fewer interactions (21 and 45, respectively; **Figure 4a**). Bivariate PIGEON-LDSC further detected 280 significant PGSxSex effects (P < 3.0e-6 using Bonferroni correction) for a total of 87 traits (**Figure 4b**). For example, we found significant BMI PGS-Sex interactions for fat-free mass and fat mass traits but with opposite directions. Here, fat-free mass assessed by bioimpedance analysis is a component of total body mass and is commonly taken as an approximation of skeletal muscle mass^52^. BMI PGS showed larger effects on fat mass traits in females than in males, but its effect on fat-free mass traits is stronger in males than in females (**Figure 4c**).

**Figure 4.**
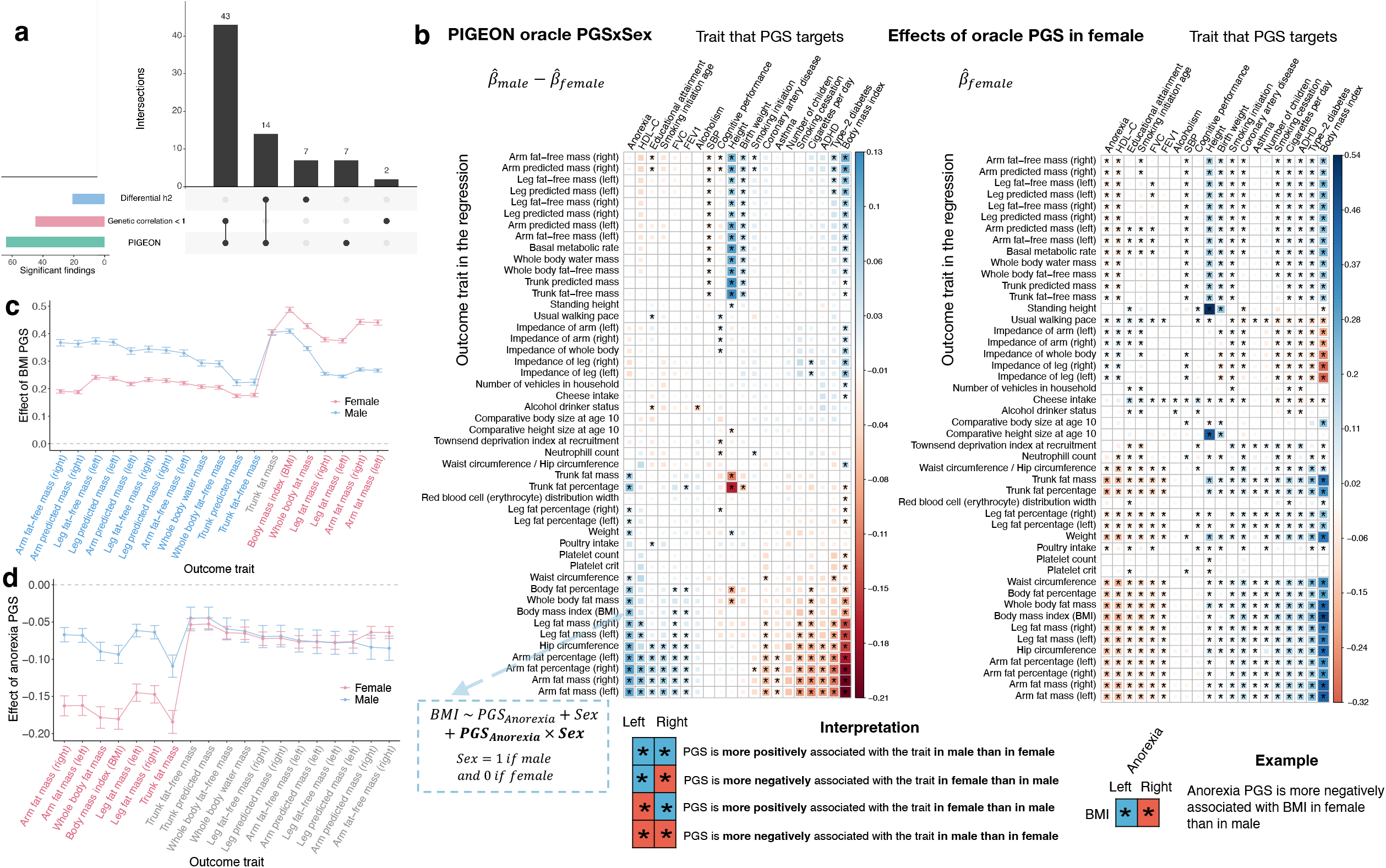
A catalog of polygenic GxSex in UKB. **(a)** The number of significant polygenic GxSex (P< 0.05/530=9.43e-5 with Bonferroni correction) identified by three approaches: PIGEON variance component estimation, differential heritability, and differential genetic correlation. **(b)** Heatmap on the left shows the results for oracle PGSxE with males coded as 1 and females as 0. The right-hand side shows the genetic covariance estimates between external GWAS and female-specific UKB GWAS to assist in interpreting the sign of interactions. Only quantitative traits showing at least one significant interaction (P< 0.05/530/30=3.14e-6 with Bonferroni correction) are shown. The complete results for all traits and all PGS are presented in **Supplementary Table 5. (c)** Effect of BMI PGS on fat mass and lean mass traits. **(d)** Effect of anorexia PGS on fat mass and lean mass traits. Significant interactions with larger PGS effects in females are highlighted in pink. Those with larger effects in males are highlighted in blue. Interactions that did not reach statistical significance are colored in grey.

We also identified a significant anorexia PGSxSex interaction on BMI (P = 5.0e-14; **Figure 4d**), suggesting that the sex difference in BMI genetics is partly explained by the genetics of anorexia – anorexia PGS is substantially more associated with lower BMI in females than in males. We replicated this finding using an independent BMI GWIS cohort from Locke et al.^47^ which does not contain the UKB sample (P = 1.0e-4). We also adjusted for BMI PGSxE in the model to account for the correlation between anorexia and BMI PGS (**Methods**). The anorexia PGS×Sex signal remained significant (P=1.0e-7), indicating an independent contribution of anorexia to the sex differences in BMI genetics. In the context of the literature on anorexia nervosa, this finding aligns with a) strong female bias in disorder presentation (9:1 ratio in females vs. males affected by anorexia^53^, and b) evidence that genetic correlations between fat percentage and anorexia differ according to sex, with more robust genetic associations between (low) body fat percentage and anorexia genetics in females as compared to males^54^. Full result of polygenic GxSex for 530 traits is provided in **Supplementary Figure 5** and **Supplementary Tables 4-5**.

### Heterogenous effect of smoking cessation treatment on lung function

Finally, we investigated PIGEON’s application in genomic precision medicine. We applied PIGEON to the Lung Health Study (LHS) to quantify the heterogenous treatment effect due to individual genetic differences. LHS is a randomized clinical trial designed to test the effectiveness of smoking intervention and bronchodilators in smokers with mild lung function impairment^55^. The main finding of the LHS was that aggressive smoking interventions significantly reduced the age-related decline in expiratory volume in one second (FEV1), but the effect of bronchodilator usage was not statistically significant^56^. Based on this, we considered FEV1 as the outcome and bronchodilator usage and aggressive smoking intervention as two exposures in our analysis (**Methods**). We found a significant interaction between smoking initiation PGS and bronchodilator usage – trial participants with a high genetic risk for smoking initiation benefited from the use of bronchodilator to reduce the decline in FEV1 (P = 6.0e-3). We verified our finding by stratifying individuals using smoking initiation PGS and estimating the bronchodilator effects on FEV1 in each subgroup (**Figure 5**). We found that the use of bronchodilator significantly reduces the declines of FEV1 among individuals in the high smoking initiation PGS group (BETA = 31.5, P = 6.7e-03), while no such effect was observed using all samples (BETA = 13.6, P = 0.1) or in the low PGS group (BETA = -2.84, P = 0.80). In contrast, the effect of smoking intervention on FEV1 was not moderated by any of the genetic scores we tested.

**Figure 5.**
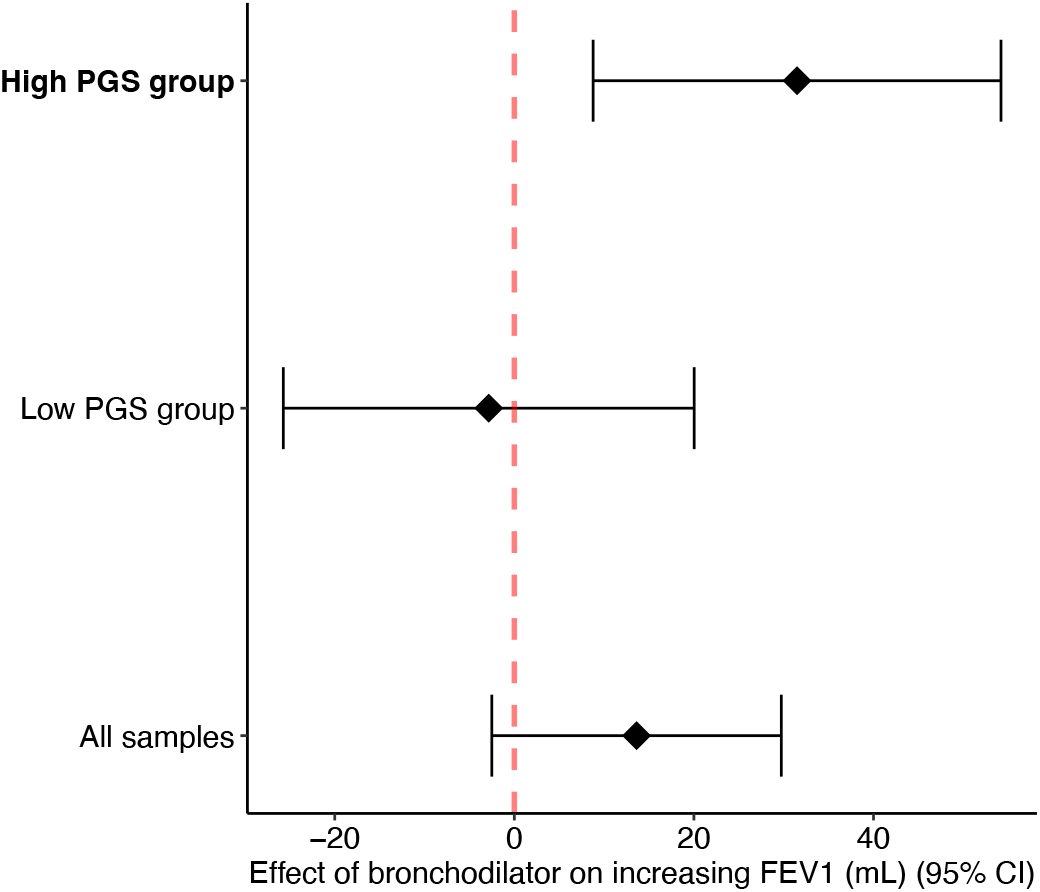
Heterogeneous treatment effect of bronchodilator on FEV1 across PGS groups. Individuals were stratified into low and high PGS groups based on their PGS of smoking initiation with median as the cutoff. The black rhombus indicates the treatment effect estimates in each group. Error bars show the 95% confidence intervals.

### Impact of gene-environment correlation on polygenic GxE inference

Finally, we investigate the impact of gene-environment correlation (rGE) on polygenic GxE inference. Under the polygenic model, we quantify rGE by allowing the environment to be heritable. Additionally, genetic effects on the environment can be correlated with both the additive effects and SNPxE interaction effects on the trait outcome (**Figure 6a**). Using this framework, we derived the bias in GxE variance component estimation introduced by rGE (**Supplementary Note 8**). Notably, given rGE, estimates of GxE variance component are unbiased under the null, suggesting that rGE will not lead to false positive results. In addition, if the additive effects on the environment and SNPxE interaction effect on the trait outcome are uncorrelated, GxE variance-covariance estimation will be unbiased. Note that this is a much weaker condition compared to the typical assumption that the environment is independent from genetics in the GxE literature.

**Figure 6.**
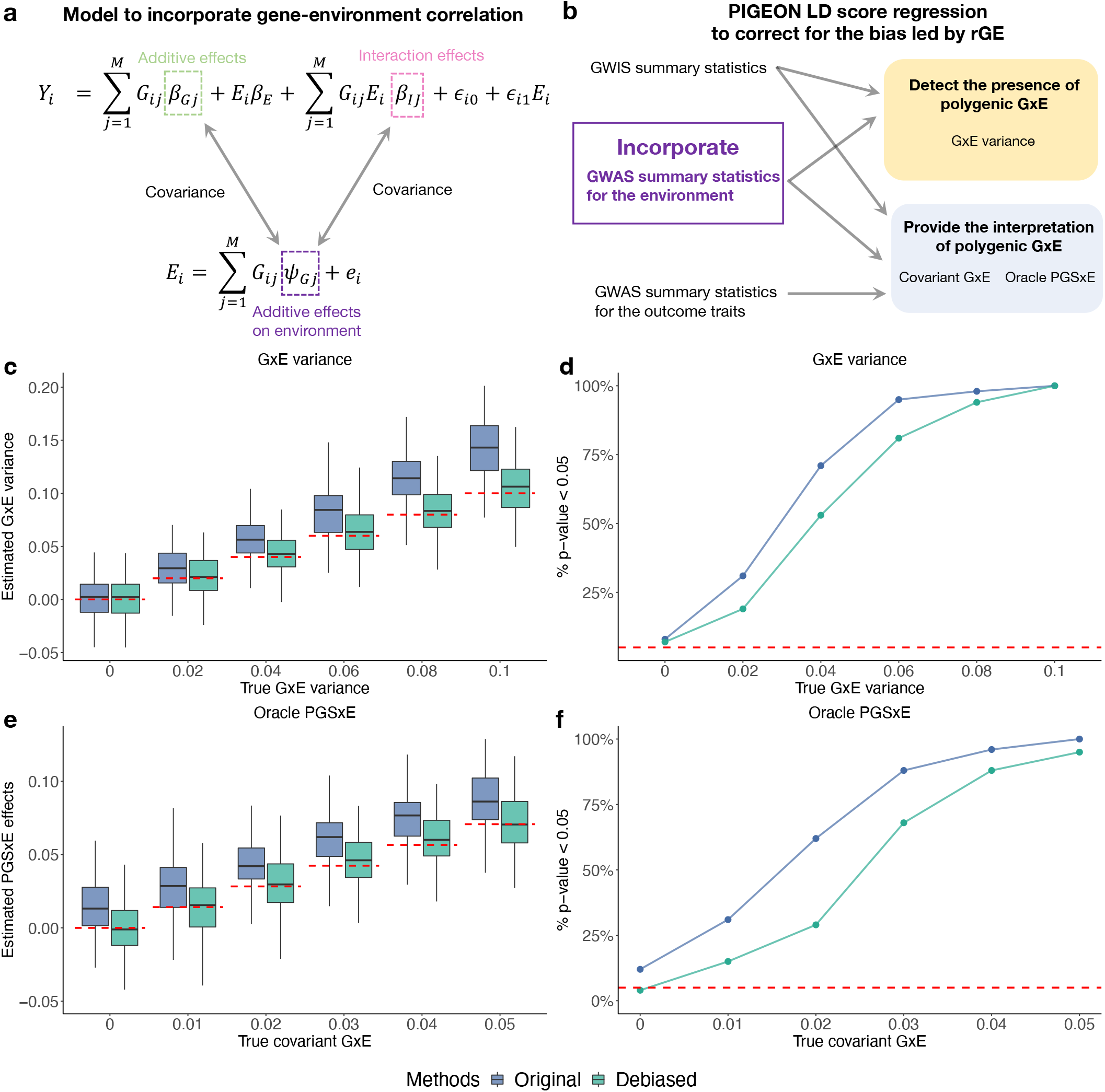
Impact of rGE on polygenic GxE inference. **(a)** To quantify rGE, we allow the environment to have a polygenic genetic basis. The additive genetic effects on the environment can be correlated with both the additive effects and GxE interaction effects on the trait outcome. (**b**) PIGEON-LDSC leverages GWAS summary statistics for the environment to correct for the bias introduced by rGE. (**c** and **d**) Simulation results for GxE variance estimation in the presence of rGE. (**e** and **f**) Simulation results for oracle PGSxE estimation in the presence of rGE. “Original” means using the methods derived under G-E independence.

If this weak condition is violated, we propose an approach to correct the bias in PIGEON parameter estimates. We extended PIGEON LDSC to obtain the debiased estimates by additionally incorporating the GWAS summary statistics for the environment (**Figure 6b, Supplementary Note 8**). Simulation results support the validity of these derivations, showing that rGE leads to biased estimates and potential false positives findings (**Supplementary Figures 6-7**), and that PIGEON provides unbiased estimates and well-controlled false positives for GxE variance, covariant GxE, and oracle PGSxE in the presence of rGE (**Figure 6c-f**).

## Discussion

We presented PIGEON, a unified statistical framework for quantifying and estimating polygenic GxE. Taking a page out of the playbook for polygenic estimation of heritability and genetic correlation, we reimagined GxE analysis from a variance component estimation perspective and demonstrated the limitations of existing GxE methods. We also developed a new estimation approach that uses GWIS and GWAS summary data alone as input. We demonstrated its statistical superiority (i.e., unbiasedness, robustness, and computationally efficient) through extensive theoretical and empirical analyses. For real data applications, we focused on three examples. We replicated and extended the PGSxEdu analysis for health-related outcomes in UKB, identified an atlas of polygenic GxSex results for 530 complex traits, and quantified genetically-moderated treatment effects in a randomized clinical trial. These detailed analyses involving diverse exposures and outcomes showcased the effectiveness of PIGEON and provided a glimpse of the broad issues that could be tackled using this analytical framework.

Our work presents several major advances that will impact future GxE studies. The first contribution is to use linear mixed model and variance-covariance estimation to quantify GxE for complex traits. This allows us to clearly define the target parameters and compare different approaches in polygenic GxE inference. We note that although GxE variance component has been introduced before in the literature^28-31^, the covariance between SNPxE effects and additive genetic effects (i.e., covariant GxE) has not been previously studied. Therefore, it is particularly interesting to discover the equivalence between covariant GxE and oracle PGSxE. This finding allowed us to quantitatively understand the limitations in existing PGSxE approaches and design better statistical estimators. Since the variance component-based analytical framework has been widely used in GWAS applications^57-59^, many advanced GWAS techniques based on this framework may be employed in future interaction studies to further facilitate our understanding of polygenic GxE.

The second major innovation in this study is to introduce the PIGEON-LDSC estimation approach. We have demonstrated several statistical features that make this approach a superior choice for polygenic GxE inference. Among these, perhaps the most important feature is its robustness to sample overlap. In current GxE practice, if the cohorts for GWAS and GxE share samples, PGS cannot be produced and PGSxE analysis is considered impossible. With PIGEON, it is now possible to produce unbiased estimates for covariant GxE and oracle PGSxE under arbitrary sample overlap. Another related but perhaps more subtle feature of PIGEON is that it enables hypothesis-free scans for PGSxE. In the current literature, most PGSxE studies are hypothesis-driven. Given an outcome and an environmental exposure of interest, researchers often hypothesize that a particular PGS moderates the exposure effect and then test PGSxE to see if this is supported by data^1,36-38^. An issue that is not discussed enough in this type of application is how to choose the PGS. With several exceptions^1,60^, most studies use the same outcome in GxE to define PGS^38,61-63^. The covariant GxE perspective in PIGEON sheds important light on this issue. We show that PGSxE analysis is essentially testing genetic correlation between the SNPxE effects in GWIS and some GWAS additive effects. Under this perspective, there is no reason to constrain GWAS and GWIS to have the same outcome. Instead, a better strategy is to perform GWIS and then test its genetic correlation with many published GWAS to gain insights into the mechanism underlying GxE. This is also where sample overlap robustness is shown to be a key feature of the framework. With PIGEON, we can assess PGSxE without concerns about whether the GWAS and GWIS were performed on the same samples or completely different ones. We studied many traits using this type of approach in this paper. For example, we investigated the GxSex GWIS results on BMI and found a significant correlation with anorexia GWAS. We believe this will motivate future GxE studies.

Third, we examined the long-standing issue regarding the impact of rGE on GxE inference. In many areas of GxE applications, it is of great interest to ensure the exogeneity of the exposure. When GxE analysis is performed on observational data, some studies go to great lengths to leverage instrumental variables^1,2^ or other approaches, while other studies ignore the potential correlation between genes and environment^60,63^. The PIGEON framework allowed us to quantitatively assess the impact of rGE. We showed that a much weaker condition than G-E independence, i.e., a zero correlation between SNP additive effects on the environment and SNPxE effects on the outcome, is sufficient for obtaining unbiased estimates and well-controlled false positive rates in polygenic GxE inference. This demonstrates that rGE does not always lead to bias in GxE analysis^64^. Even when this condition is violated, we proposed a strategy within the PIGEON framework to correct for biases introduced by rGE.

Finally, we once again reimagine how future GxE studies may unfold. The incredible success of complex trait genetics that the field has achieved in the past 15 years is largely credited to GWAS meta-analysis conducted by big genetics consortia, sharing of summary association statistics, and statistical analysis only requiring summary data as input^65-67^. The GxE field, however, largely remains in the early GWAS era. GxE analysis is almost always performed in a small cohort with individual-level genetic, exposure, and outcome data. PIGEON is a clear demonstration that modern statistical genetics that embrace the “omnigenicity” of human traits and rely on summary data alone can also apply to GxE research. Based on this, we make a bold prediction – the future success of complex trait GxE research resides in sharing and meta-analyzing GWIS summary statistics^68^, and future GxE method development should focus on techniques that only rely on summary-level data.

Our study is not without limitations. First, we followed a model widely used in the GWAS literature and assumed equal contribution of each standardized SNP for both additive and GxE variance components. A future direction is to incorporate more flexible assumptions on the SNPxE effect size distribution, e.g., allele frequency-dependent models such as LDAK^69^, into the PIGEON framework. Second, we employed a LDSC-type inference procedure to estimate the variance-covariance components in polygenic GxE analysis. Some recent methods have demonstrated improved efficiency in heritability and genetic correlation estimation compared to LDSC^59,70,71^. Whether similar techniques can be incorporated in PIGEON is an interesting open problem warranting future investigation. Third, although we explored diverse types of exposure data in this study, we did not investigate whether PIGEON can be used to study gene-gene interactions^72^. Quantifying how the effects of certain focal genes and variants can be modified by the polygenic genetic background is a topic that is conceptually similar to polygenic GxE^73^. Generalizing PIGEON to GxG applications will be an interesting future direction. Fourth, any effort to produce a universally applicable PGS implicitly assumes genetic effects to be identical across the environments which fundamentally contradicts the main question in GxE^46^. In fact, some PGS have notoriously low “portability” between different environments^74^ and it remains an open question how this issue will affect PGSxE inference results. We are not aware of any current solution to this problem under the empirical PGSxE design, but we derived the necessary conditions for the lack of PGS portability to lead to biases in GxE inference and proposed new analysis strategies to de-bias the estimates (**Supplementary Note 9**). An important future direction is to validate these results in empirical studies.

Taken together, PIGEON is a general and powerful framework that may reshape how we perform complex trait GxE studies in the future. We showcased its performance and demonstrated its superiority over existing methods through numerous examples targeting diverse types of GxE problems. We believe PIGEON offers an innovative solution to many GxE challenges. If the field could follow the success of GWAS and make GWIS summary statistics accessible, it will provide tremendous opportunities to study the relation between genes and environments and provide insights into polygenic GxE for many human complex traits.

## Methods

### PIGEON model

PIGEON is built on a linear mixed model assuming the trait outcome to be influenced by polygenic SNP additive effects, environment effect, polygenic SNPxE effects, residual, and residual-environment interaction (RxE):

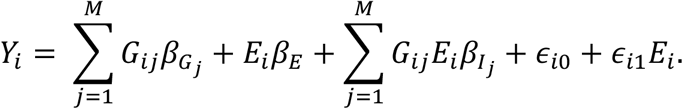

Here, *Y*_*i*_ is the standardized phenotype with a mean of 0 and variance of 1 for the i-th individual, *G*_*ij*_ is the j-th standardized SNP, *E*_*i*_ is the standardized environment, *ϵ*_*i*0_ is the noise term, and *ϵ*_*i*1_*E* quantifies the heteroskedasticity due to RxE^30,39^. Here, we assume the environment to have zero heritability, but implications of rGE are discussed in **Supplementary Note 8**. We treat *β*_*E*_ as fixed and 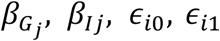 as random. We model all these random variables as independent except for 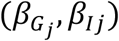 and (*ϵ*_*i*0_, *ϵ*_*i*1_). We assume that 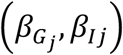 and (*ϵ*_*i*0_, *ϵ*_*i*1_) have mean zero and covariance matrix below:

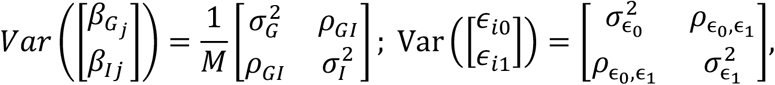

where 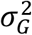 and 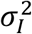 are the variance explained by SNP additive and SNPxE effects, *ρ*_*GI*_ denotes the covariant GxE, 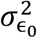 and 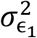 are the variance of residual and RxE, and 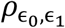 denotes their covariance. Technical discussion of PIGEON model can be found in **Supplementary Note 1**.

### Consolidation and comparison of commonly-applied GxE methods

Next, we consolidate and compare several GxE methods under the PIGEON model. The detailed derivation and technical discussion can be found in **Supplementary Note 2**. To compare that can only be applied to the discrete environment (i.e., differential heritability, differential genetic variance, and testing for genetic correlation < 1), we first consider a special case of our PIGEON model where the raw environment is binary. In this case, we can rewrite the PIGEON model into a pair of environment-stratified models:

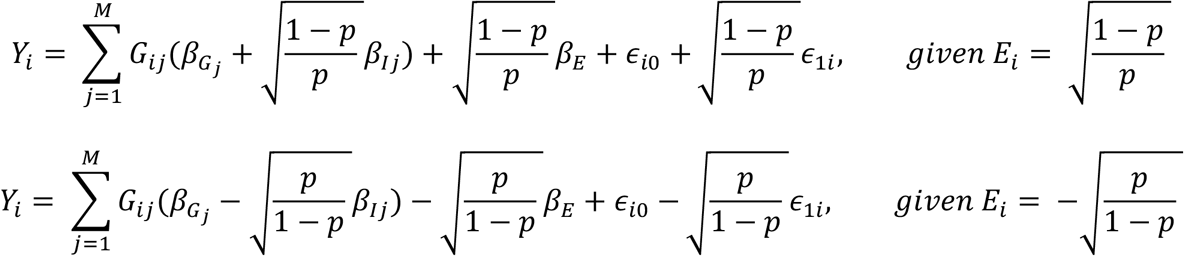

where 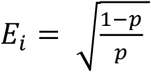 and 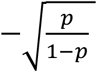 represents the standardized Bernoulli random variable with probability *p* of being 1 and 1 − *p* being 0.

#### Heritability difference

We can show that the heritability difference between populations with 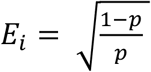 and 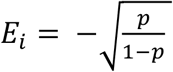 is

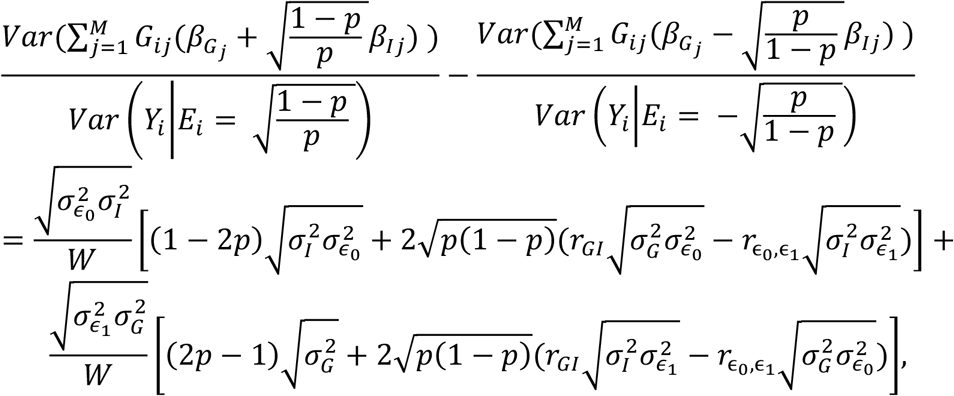

where 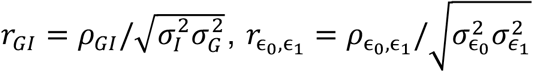 and 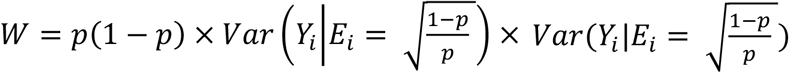 is a scaling factor related to the phenotypic variance. Therefore, the heritability could still differ between two environments due to heteroskedasticity 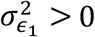even without any 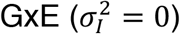.

#### Genetic variance difference

The genetic variance difference between populations with 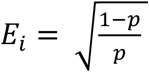 and 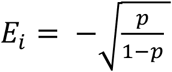 can be denoted as

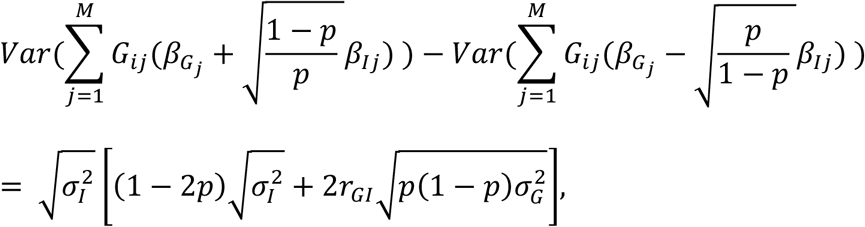

where 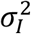 is the GxE variance components. This shows that the genetic variance could be the same across different environments even if there exists 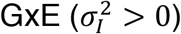.

#### Genetic correlation = 1

It can be shown that genetic correlation = 1 between populations with 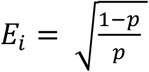 and 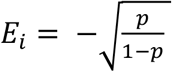 if and only if

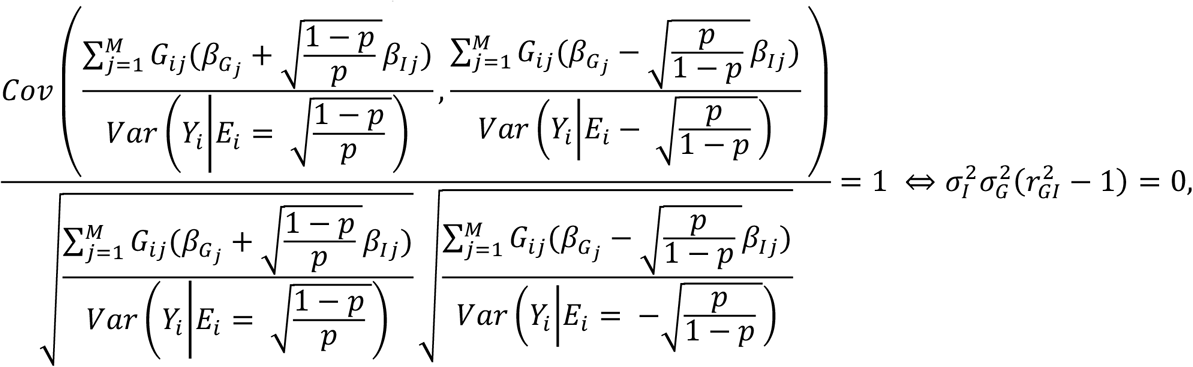

where 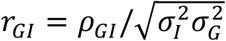. Therefore, even if there are 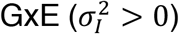, the genetic correlation could be 1 between two environments if the SNP additive and SNPxE effects are perfectly correlated (i.e., *r*_*GI*_ = ±1).

#### Oracle PGSxE

The oracle PGSxE regression can be denoted as

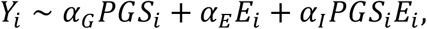

where *Y*_*i*_ is the standardized phenotype, 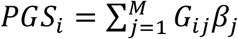 is the oracle PGS based on each SNPs’ true effect *β*_*j*_, *E*_*i*_ denotes the standardized environment with no heritability, and *α*_*I*_ is the interaction coefficient to be estimated. Under the PIGEON model, we showed that normalizing covariant GxE by additive heritability yields oracle PGSxE effects

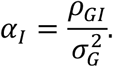

In our analysis and in the implemented software, we estimated the oracle PGSxE coefficient

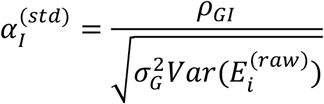

based on the standardized phenotype and PGS (i.e., *Y*_*i*_ and 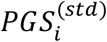), and raw scale environment 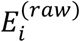 for the regression 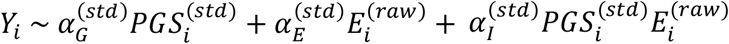 to ensure the interpretation for the coefficient. We note that the equivalence of hypothesis testing for oracle PGSxE and covariant GxE is not affected by the scale and location transformation of phenotype, PGS, and environment (**Supplementary Note 2**). We further present similar results for oracle PGSxE by using PGS for traits other than the outcome trait in the regression in **Supplementary Note 5**.

#### Empirical PGSxE without sample overlap

Next, we consider the current design for empirical PGSxE analysis. We denote the empirical PGSxE regression as 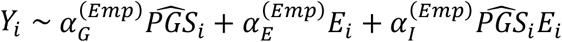 where 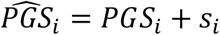 is a noisy version of the oracle PGS, and *s*_*i*_ represents the estimation error in empirical PGS with zero mean and *Cov*(*PGS*_*i*_, *s*_*i*_) = 0. We first consider the case where the GWAS used to generate the PGS have no sample overlap with the empirical PGSxE cohort, represented by *Cov*(*s*_*i*_, *Y*_*i*_) = 0. Under the assumption described above and *Var*(*s*_*i*_) = *M*_*eff*_/*N*_*G*_ described in Daetwyler et al.^75^, we have the expectation of the least squares estimator for the empirical PGSxE regression coefficient:

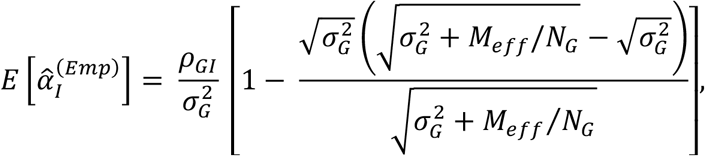

where *M*_*eff*_ is the effective number of independent SNPs^76^, *ρ*_*GI*_ is the covariant GxE, and *N*_*G*_ is the GWAS sample size. This shows that the empirical PGSxE estimator is biased towards zero compared to the oracle PGSxE coefficient 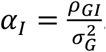. The oracle PGSxE is the upper bound 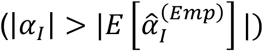 and the infinity sample (GWAS sample size) limit of the empirical PGSxE 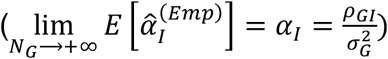.

#### Empirical PGSxE with sample overlap

Next, we consider the case where the GWAS used to generate PGS have *N*_*S*_ shared samples with the empirical PGSxE cohort with *N*_*I*_ individuals, which can be quantified by *Cov*(*s*_*i*_, *Y*_*i*_) ≠ 0. Then, we have the expectation of the least squares estimator for the empirical PGSxE coefficient

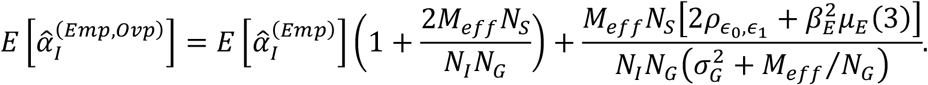

Therefore, the sample overlap will lead to biases and false positive results for empirical PGSxE analysis due to the existence of the second term 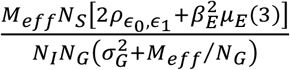.

### PIGEON LD score regression

To estimate the GxE variance component, we only need Z-scores for SNPxE effects in GWIS summary statistics. The expected value of the squared Z-score for the j-th SNPxE interaction effect *z*_*Ij*_ is

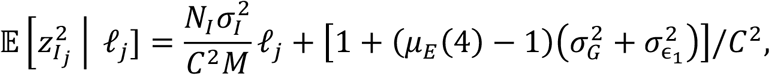

where *N*_*I*_ denotes the GWIS sample size, 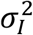 is the GxE variance component, *M* is the number of SNPs, 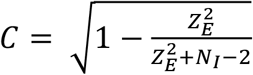 is a correction factor to account for the environmental effect on Z-score approximation in GWIS (**Supplementary Note 4** and **10**), 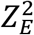 is the Z-score of environmental effect, *ℓ*_*j*_ is the LD score, and *μ*_*E*_(4) is the kurtosis of the environment.

To estimate covariant GxE, we only require the GWIS and GWAS summary statistics with arbitrary sample overlap. The expected value of the product of additive effect Z-scores and SNPxE effect Z-scores is

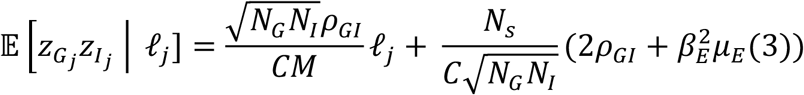

where *N*_*I*_ and *N*_*G*_ represents the GWIS and GWAS sample size, *ρ*_*GI*_ is the covariant GxE, *M* is the number of SNPs, 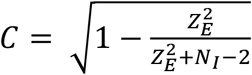 is a correction factor described above, 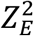 is the Z-score of environment effects, *ℓ*_*j*_ is the LD score, *N*_*s*_ is the number of overlapped samples between GWIS and GWAS analysis, *μ*_*E*_(3) is the skewness of the environment. The oracle PGS can be obtained by normalizing the covariant GxE by heritability. We use block jackknife to calculate the standard error of the estimates and regression weights to account for heteroskedasticity of residual in PIGEON-LDSC^77^. The detailed derivation of PIGEON-LDSC can be found in the **Supplementary Note 4**.

### Simulation settings

We conducted a series of simulations using imputed genotype data from UKB. We restricted the analysis to autosomal SNPs with imputation quality score > 0.9, minor allele frequency (MAF) ≥ 0.05, missing call rate ≤ 0.01, and Hardy-Weinberg equilibrium test p-value ≥ 1.0e-6. We further extracted SNPs in the HapMap3 SNP list and 1000 Genomes Project Phase III LD reference data for European ancestry^78^. 734,046 SNPs remained after QC. We randomly selected 40,000 independent samples with European ancestry and equally divided them into two sub-cohorts with 20,000 each. The simulations are repeated 100 times.

We used the first sub-cohort for GxE variance component simulations. We evaluated the point estimates of PIGEON and compared the statistical power of PIGEON with differential heritability and genetic correlation < 1 analyses. We first generated the binary environment from a Bernoulli distribution with a probability of 0.5 being 1 and then standardized it to have a mean of zero and variance of 1. We then simulated the phenotype using standardized genotype *G*_*ij*_ and standardized environment *E*_*i*_ by PIGEON model where the SNP effect size and residual were simulated from a multivariate normal distribution

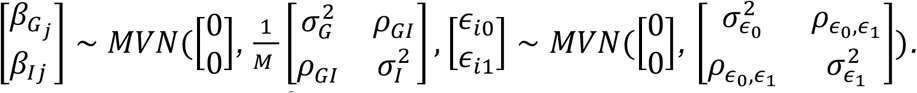

Here, we set the GxE variance 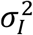 value to be 0, 0.02, 0.04, 0.06, 0.08, and 0.1, heritability 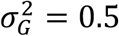, covariant 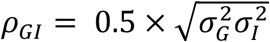, residual variance 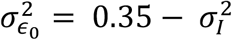, heteroskedasticity variance 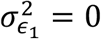, and the environmental effect 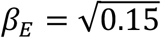 such that the variance of the phenotype is 1. We note that non-zero heteroskedasticity variance may lead to type-I error inflation in differential heritability analysis. Therefore, to ensure the fair comparison of statistical power between differential heritability and PIGEON, we did not consider a non-zero heteroskedasticity variance 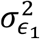 in this simulation. Instead, we investigated its impact in our secondary simulations (**Supplementary Note 11**). We used PLINK^79^ to obtain the summary statistics from genome-wide SNPxE and environment-stratified GWAS analyses. Then, these summary statistics were used as input for all approaches. Both differential heritability and genetic correlation < 1 analyses were implemented using LDSC^21,77^. The LD scores were estimated using the whole 40,000 individuals. We further reduced the GWIS sample size to 5,000 and replicate the analysis to mimic the unbalanced sample size between GWIS and GWAS in real applications. Next, we compared PIGEON with the empirical PGSxE approach based on two PGS: clumping PRS and PRS-CS under zero and 100% sample overlap. We used the PIGEON model with SNP effect size and residual simulated from a multivariate normal distribution with heritability 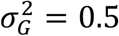, GxE variance 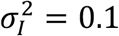, covariant GxE *ρ*_*GI*_ with values 0, 0.01, 0.02, 0.03, 0.04, and 0.05, residual variance 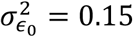, heteroskedasticity variance 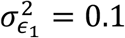, covariant residual 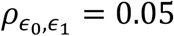, and the environmental effect 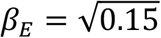 using the same simulated environment described above. Summary statistics for GWAS and GWIS were generated in different and the same sub-cohort for zero and 100% sample overlap settings, respectively. We used GWAS summary statistics as input for PRSice-2^44^ with its default setting (--clump-kb=250, --clump-r2=0.1, and --clump-p=1) and PRS-CS-auto^45^ to generate the clumping and PRS-CS scores, respectively. We aimed to estimate the oracle PGSxE effects based on the standardized phenotype, standardized PGS, and raw-scale environment. We used summary statistics for GWAS and GWIS as input for PIGEON. When there is no sample overlap, we restrict the intercept in PIGEON LDSC to 0 to reduce the standard error for the estimates. For empirical PGSxE, we fit a regression for 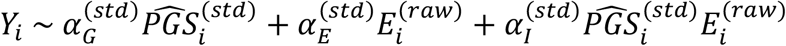, where 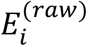 is the raw binary environment, *Y*_*i*_ and 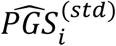 are standardized to have a mean of 0 and a variance of 1. For each replicate, we recorded the estimates for 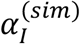 and corresponding p-values to test 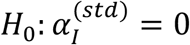. The details for secondary simulations and additional analyses with the presence of rGE can be found in **Supplementary Note 11**.

### PGSxEdu interaction for health outcomes

We replicated the analysis in Barcellos et al.^1^ to study whether genetics moderate the effect of education reform on health-related outcomes. Following Barcellos et al., we considered both continuous and dichotomous summary indices for health as outcomes. Details of constructing the summary indices were described before^1^. Briefly, the continuous summary index is a weighted average of body size, blood pressure, and lung function traits, and it is coded such that a higher number indicates worse health. The dichotomous threshold summary index is an indicator for whether the continuous summary index is above a threshold. We followed Barcellos et al. to restrict the samples to individuals with European ancestry and further removed the related individuals. 172,664 individuals with phenotype data remained in the analysis after QC. We used the imputed genotype data provided by UKB throughout the analysis.

We used same BMI and EA GWAS as input for PRS-CS^45^ to generate PGS and same model to perform PGSxE regression as Barcellos et al.^1^. We fine-tuned the PGS model using a validation set of 10,000 individuals randomly selected from remaining UKB samples. We used the month of birth rather than the date of birth to cluster the standard errors due to the limited data access^1^. For the PIGEON analysis, we generated GWIS summary statistics using the same PGSxE model except for changing PGS into SNPs. We then applied PIGEON-LDSC to estimate the oracle PGSxE using GWIS and the two GWAS described above. LD scores were calculated in the UKB EUR samples^80^. We also estimated the oracle PGSxE using two recently-published, larger GWAS for BMI^49^ and EA^50^ which included UKB samples. Further, we used PIGEON to perform a hypothesis-free scan for PGSxEdu interactions using GWAS summary statistics for 30 complex traits (**Supplementary Table 1**).

### Polygenic GxSex interaction for 530 complex traits

We estimated GxSex variance component and performed a hypothesis-free scan for oracle PGSxSex interactions using GWAS summary statistics for 30 complex traits (**Supplementary Table 1**). We transformed the sex-stratified GWAS summary statistics released by Bernabeu et al.^27^ into SNPxSex summary statistics by

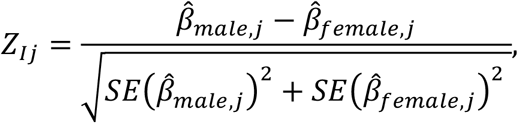

where *Z*_*Ij*_ is the SNPxSex interaction Z-score for the j-th SNP, and 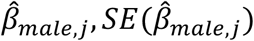 and 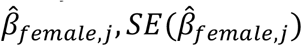 are estimated SNP effects and their standard error in sex-stratified GWAS summary statistics. We also estimated additive effect genetic correlations between 530 UKB traits and 30 GWAS to help interpret the sign of polygenic GxSex interactions using LDSC. We determined the significant findings by a Bonferroni-corrected P-value cutoff 0.05/530 = 9.43e-5 for GxSex variance component and 0.05/530/30 = 3.14e-6 for oracle PGSxSex. To rule out the possibility that the anorexia oracle PGSxSex signals is driven by the genetic overlap between BMI and anorexia, we introduced the conditional oracle PGSxE analysis in PIGEON, where we fit a multiple regression model conditioning on the BMI PGSxE effect (**Supplementary Note 6**).

### LHS data analysis

A detailed description of LHS has been provided elsewhere^55^. We applied pre-imputation QC by keeping autosomal biallelic SNPs with MAF > 0.01 and Hardy-Weinberg equilibrium test p-value ≥ 1.0e-6. We phased and imputed the genotype data using the Haplotype Reference Consortium reference panel version r1.1 2016 available on the Michigan Imputation server^81^. We also performed post-imputation QC by removing the duplicated and strand-ambiguous SNPs as well as SNPs with MAF < 0.01 and imputation quality < 0.9. 12,030,369 SNPs remained after QC.

LHS participants were assigned into three non-overlapping groups: SIA (smoking intervention and the inhaled bronchodilator ipratropium bromide), SIP (smoking intervention and an inhaled placebo), and UC (usual care who received no intervention). We considered the average change of the post-treatment FEV1 from the baseline as the outcome. We performed two separate analyses, one using bronchodilators and the other using smoking interventions as the treatment. We only included individuals with FEV1 measurements at both baseline and all five annual follow-up visits in the analyses^82^. When considering the bronchodilator (ipratropium bromide) as the treatment, we excluded individuals in UC group, and coded the individuals in SIA group as 1 and SIP group as 0 (N=2,089). When considering the smoking interventions as the treatment, we excluded individuals in SIA group, and coded the individuals in SIP group as 1 and UC group as 0 (N=2,122). We performed the genome-wide SNPxE analysis using PLINK with sex, age, age^2^, age×sex, age^2^×sex, 20 genetic principal components computed using flashPCA2^83^, and interaction between the treatment and these variables as covariates^84^. Then, we estimated oracle PGSxE using the GWIS and 30 GWAS summary statistics using PIGEON. Since LHS has no sample overlap with any of these GWAS, we constrained the intercept to be 0 to improve estimation efficiency. We used PRS-CS-auto to calculate the smoking initiation PGS. We estimated the treatment effects using all samples or stratified samples in high/low PGS groups defined by the median of the smoking initiation PGS.

### URLs

UK Biobank (http://www.ukbiobank.ac.uk/);

Lung Health Study (https://clinicaltrials.gov/ct2/show/NCT00000568);

PRS-CS (https://github.com/getian107/PRScs);

PRSice-2 (https://github.com/choishingwan/PRSice);

PLINK (https://www.cog-genomics.org/plink/2.0/);

LDSC (https://github.com/bulik/ldsc)

### Data and code availability

PIGEON software package is publicly available at https://github.com/qlu-lab/PIGEON The SNPxE summary statistics used in the paper are available at http://qlu-lab.org/data.html

## Acknowledgments

The authors gratefully acknowledge research support from National Institutes of Health (NIH) grant U01 HG012039, the National Institute on Aging (NIA) (R00 AG056599), and support from the University of Wisconsin-Madison Office of the Chancellor and the Vice Chancellor for Research and Graduate Education with funding from the Wisconsin Alumni Research Foundation (WARF). We also acknowledge use of the facilities of the Center for Demography of Health and Aging at the University of Wisconsin-Madison, funded by NIA Center Grant P30 AG017266. We thank members of the Social Genomics Working Group at University of Wisconsin for helpful comments. This research has been conducted using the UK Biobank Resource under Application 42148.

## Author contribution

J.M. and Q.L. conceived and designed the study.

J.M. developed the statistical framework, performed the simulations and data analysis, and implemented the software.

S.G. assisted in GxSex analysis in UK Biobank.

Y.W. assisted in implementing the software

J.H. assisted in developing the statistical framework.

Y.W. assisted in UKB data preparation.

S.B. assisted in Lung Health Study data preparation.

J.S.A., K.S., L.S., and J.F. advised on result interpretation.

Q.L. advised on statistical and genetic issues.

J.M. and Q.L. wrote the manuscript.

All authors contributed to manuscript editing and approved the manuscript.

## Notes

### Competing Interest Statement

The authors have declared no competing interest.

