## Supplementary Figures for "Reimagining Gene-Environment Interaction Analysis for Human Complex Traits"

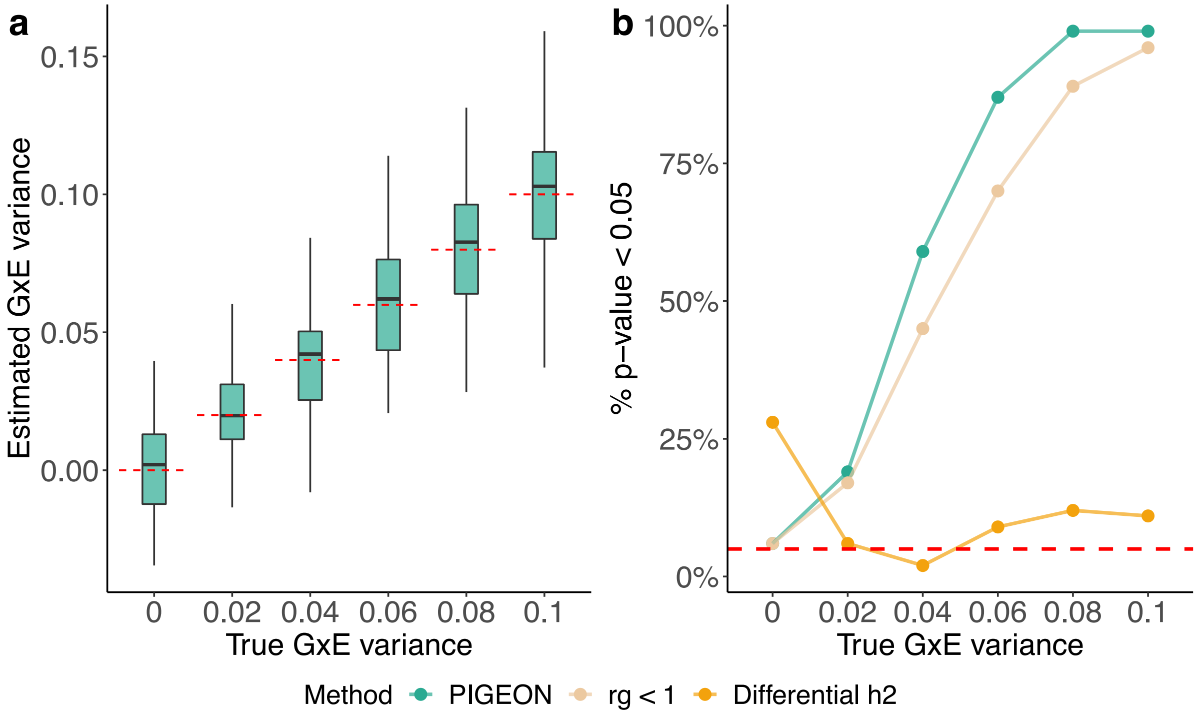


**Supplementary Figure 1. Comparison of GxE approaches for estimating GxE variance component in the presence of heteroskedasticity (i.e., RxE).**

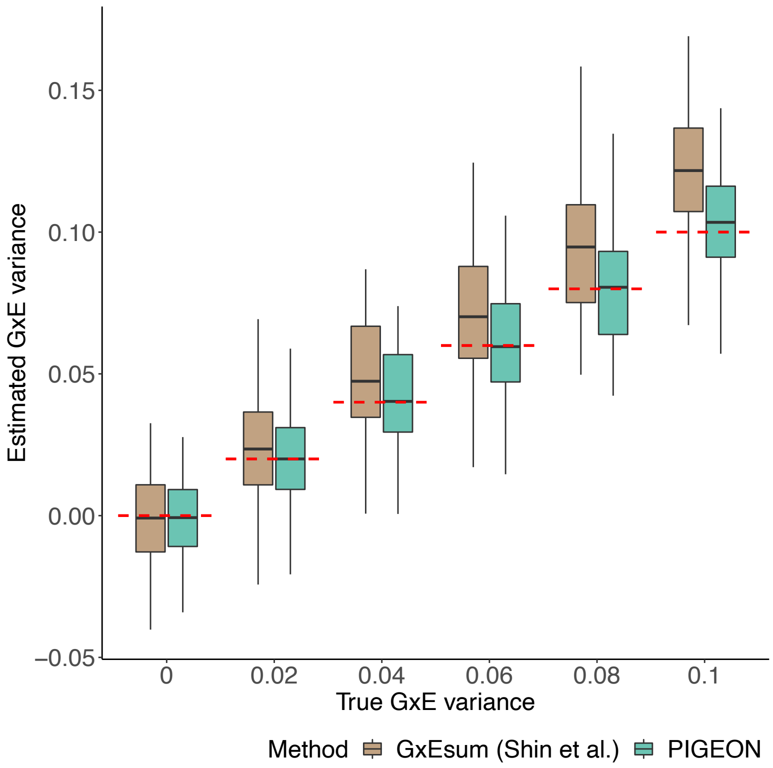


**Supplementary Figure 2. Comparison of PIGEON and GxEsum for estimating GxE variance component without rGE.** Both methods used SNPxE interaction Z-score from GWIS as input.

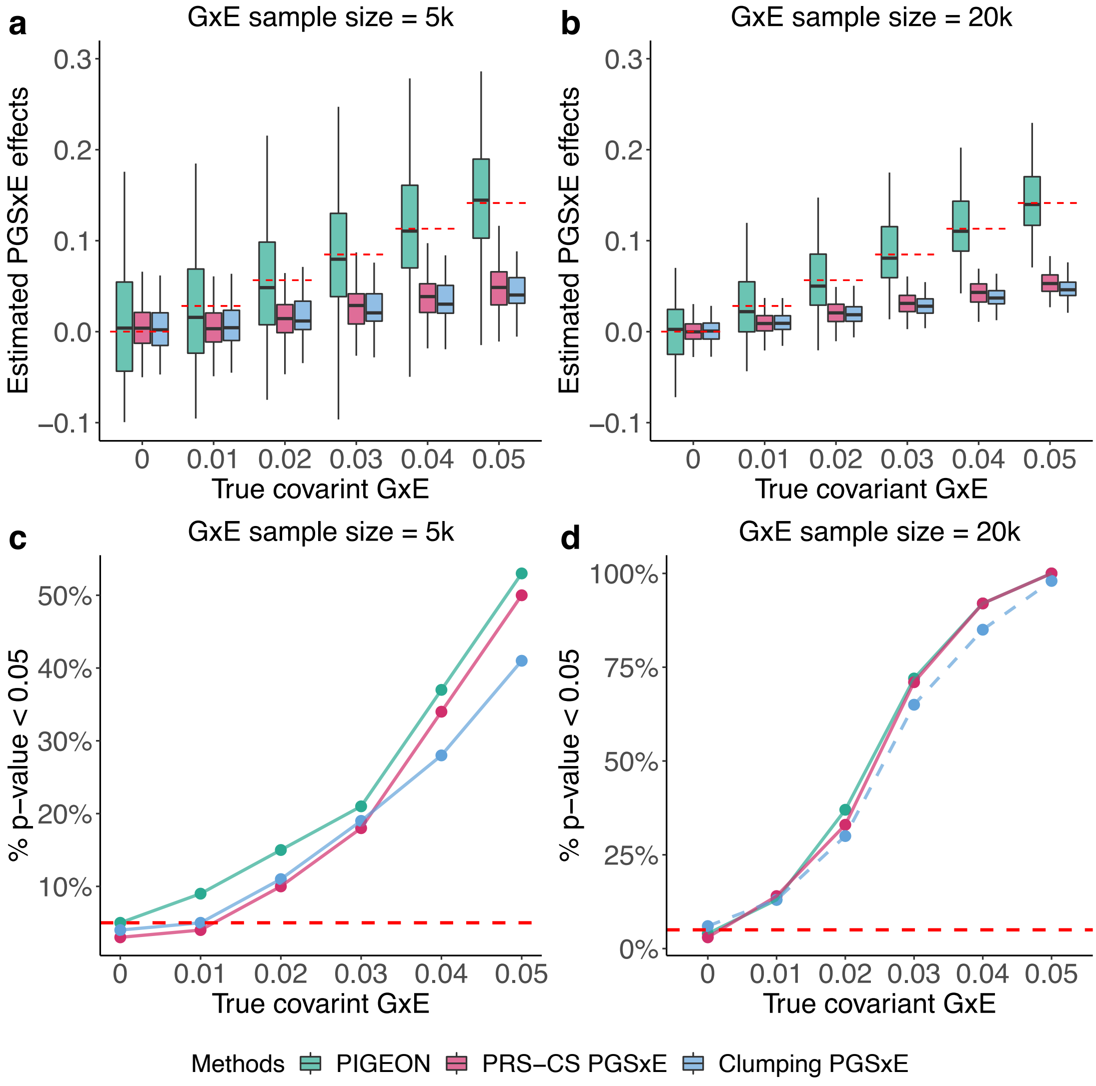


**Supplementary Figure 3. Comparison of PIGEON and empirical PGSxE under various sample size ratios between GWIS and GWAS. (a** and **c)** Simulations results with 5,000 samples in the GxE cohort and 20,000 samples in the GWAS cohort. **(b** and **d)** Simulations results with 20,000 samples in the GxE cohort and 20,000 samples in the GWAS cohort.

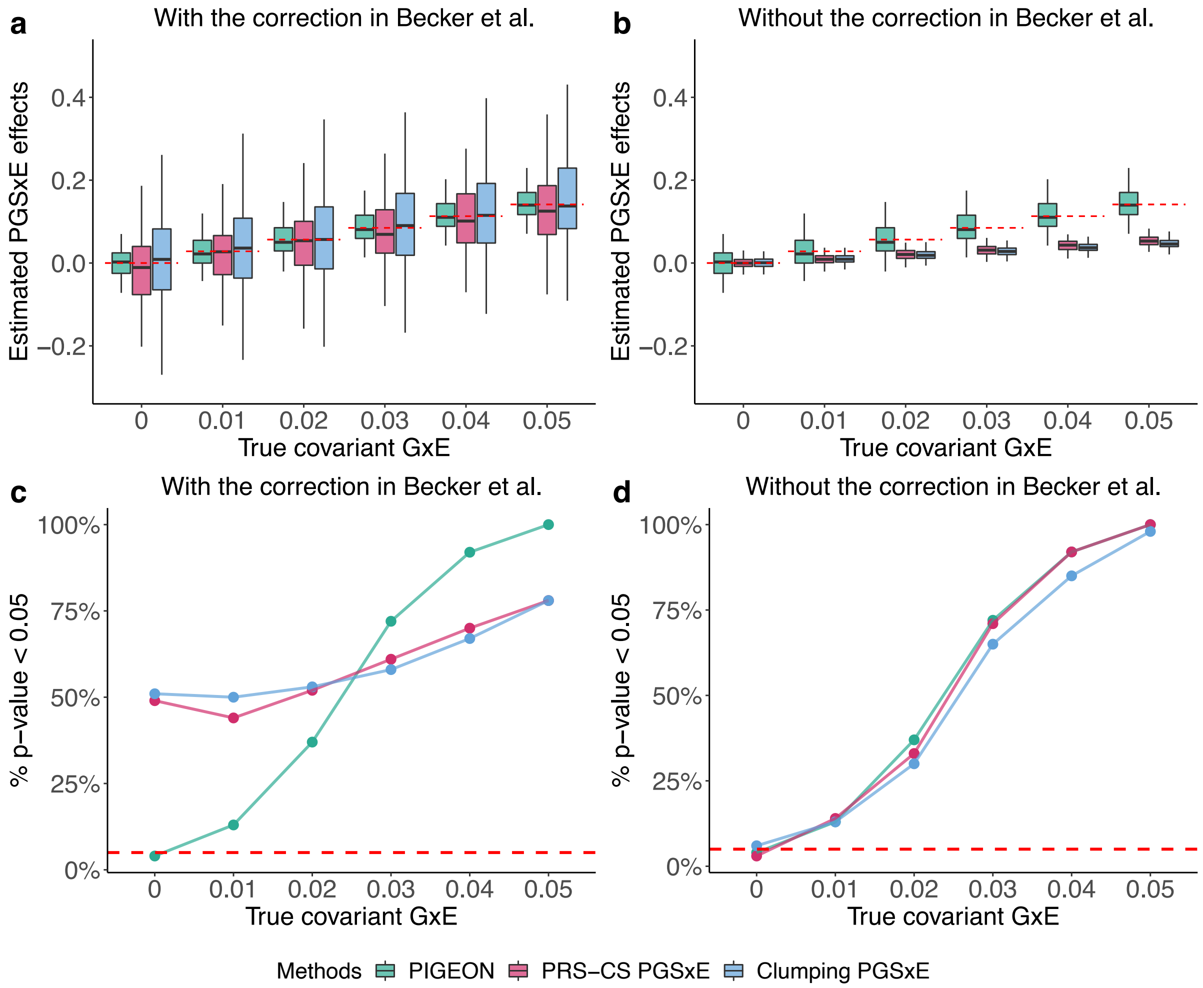


**Supplementary Figure 4. Comparison of PIGEON and measurement error-corrected PGSxE estimator in Becker et al.**

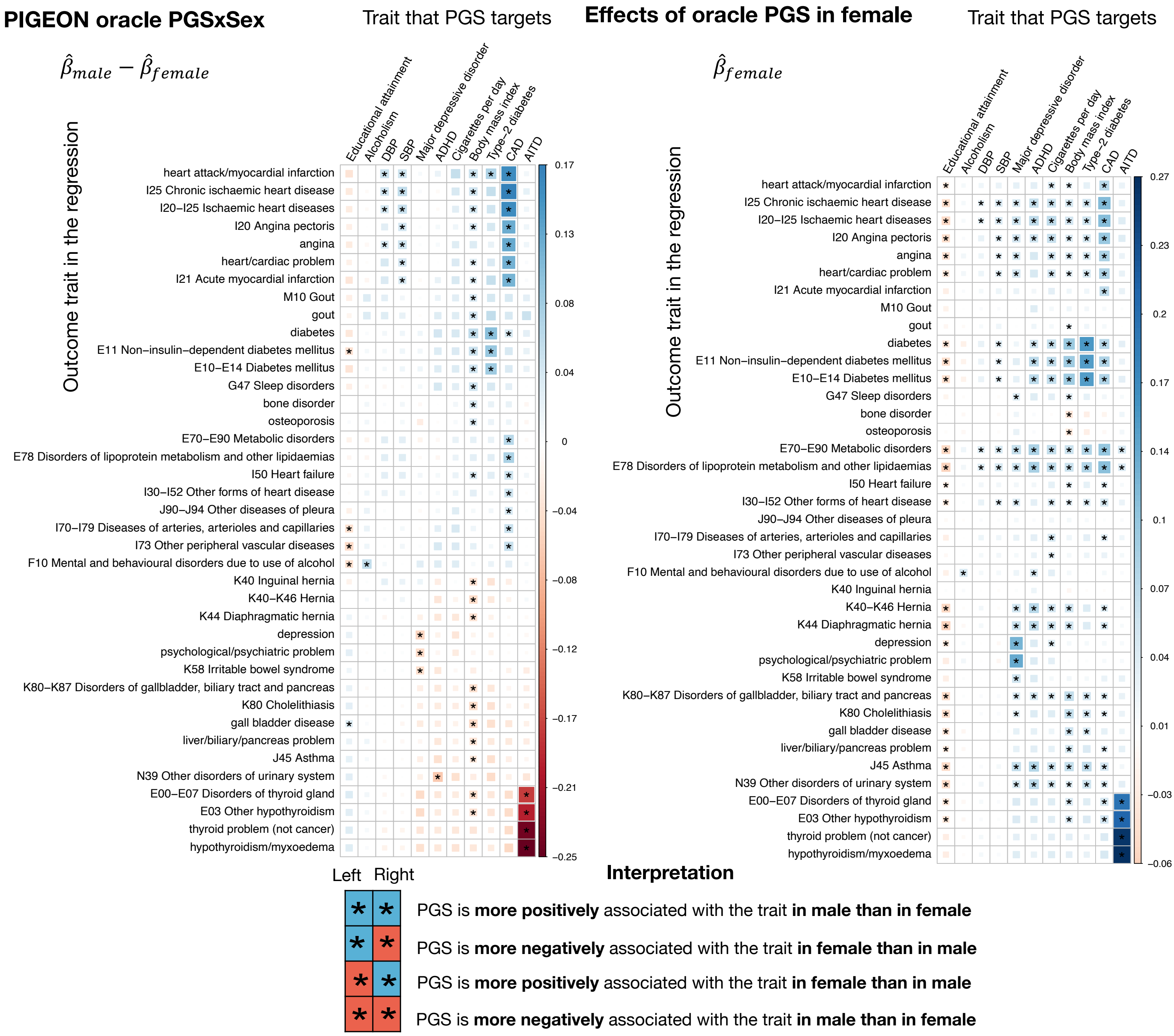


**Supplementary Figure 5. Oracle GxSex interaction for binary traits in UKB.** Only traits and PGS with at least one significant interaction are shown. The left-hand side shows the results for oracle PGSxE with males coded as 1 and females as 0. The right-hand side shows the results for additive genetic covariance results in females to interpret the sign of interaction. The complete results for all binary traits and all PGS can be found in **Supplementary Table 5**.

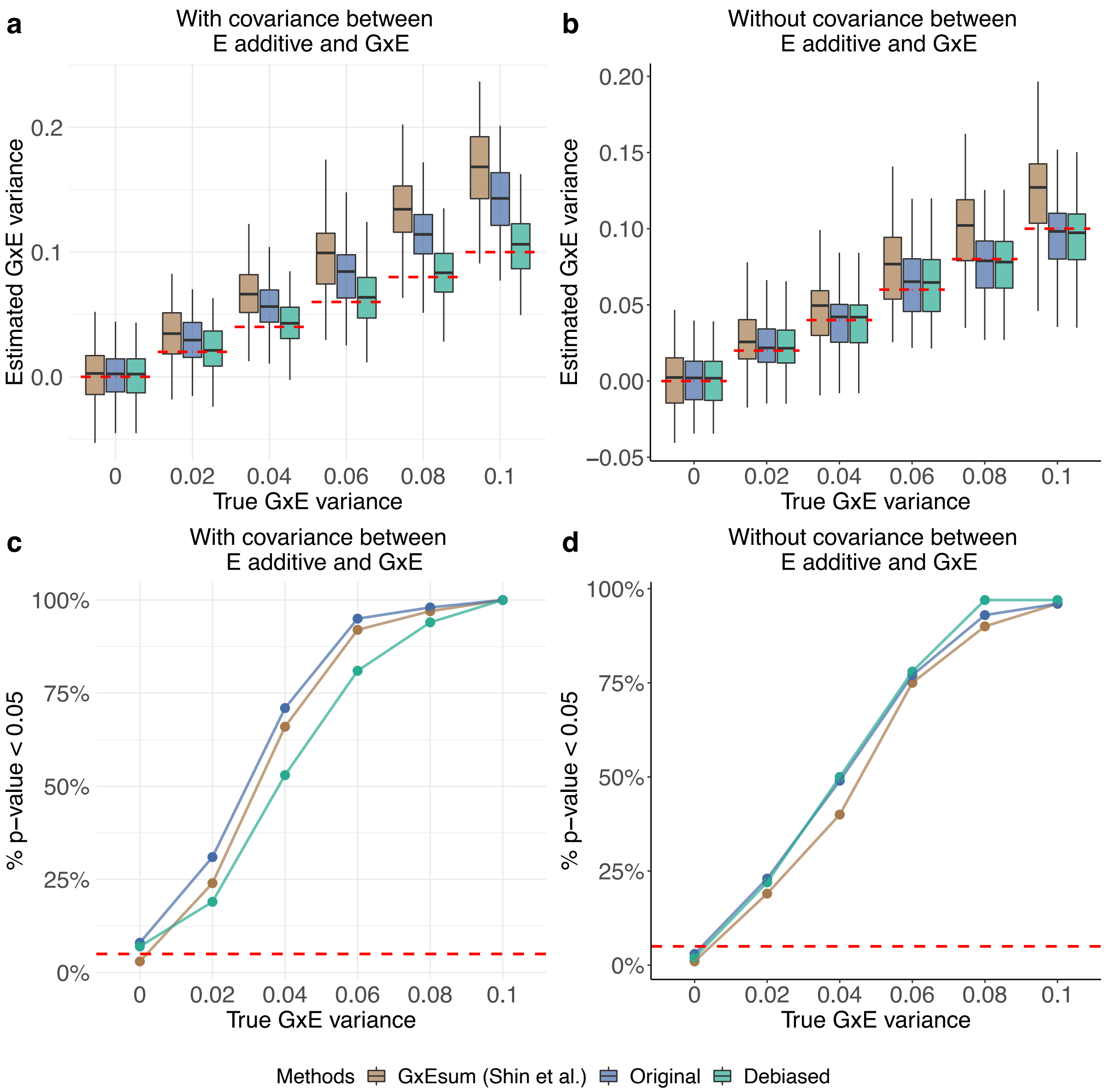


**Supplementary Figure 6. Simulation results of PIGEON for estimating GxE variance in the presence of rGE. (a** and **c)** Simulation results when the additive genetic effects on the environment are uncorrelated with the GxE effects on the outcome trait. **(b** and **d)** Simulations results when the additive genetic effects on the environment and GxE effects on the outcome trait are correlated. "GxEsum (Shin et al.)" means applying the GxEsum proposed in Shin et al. "Original" means using the methods derived under G-E independence.

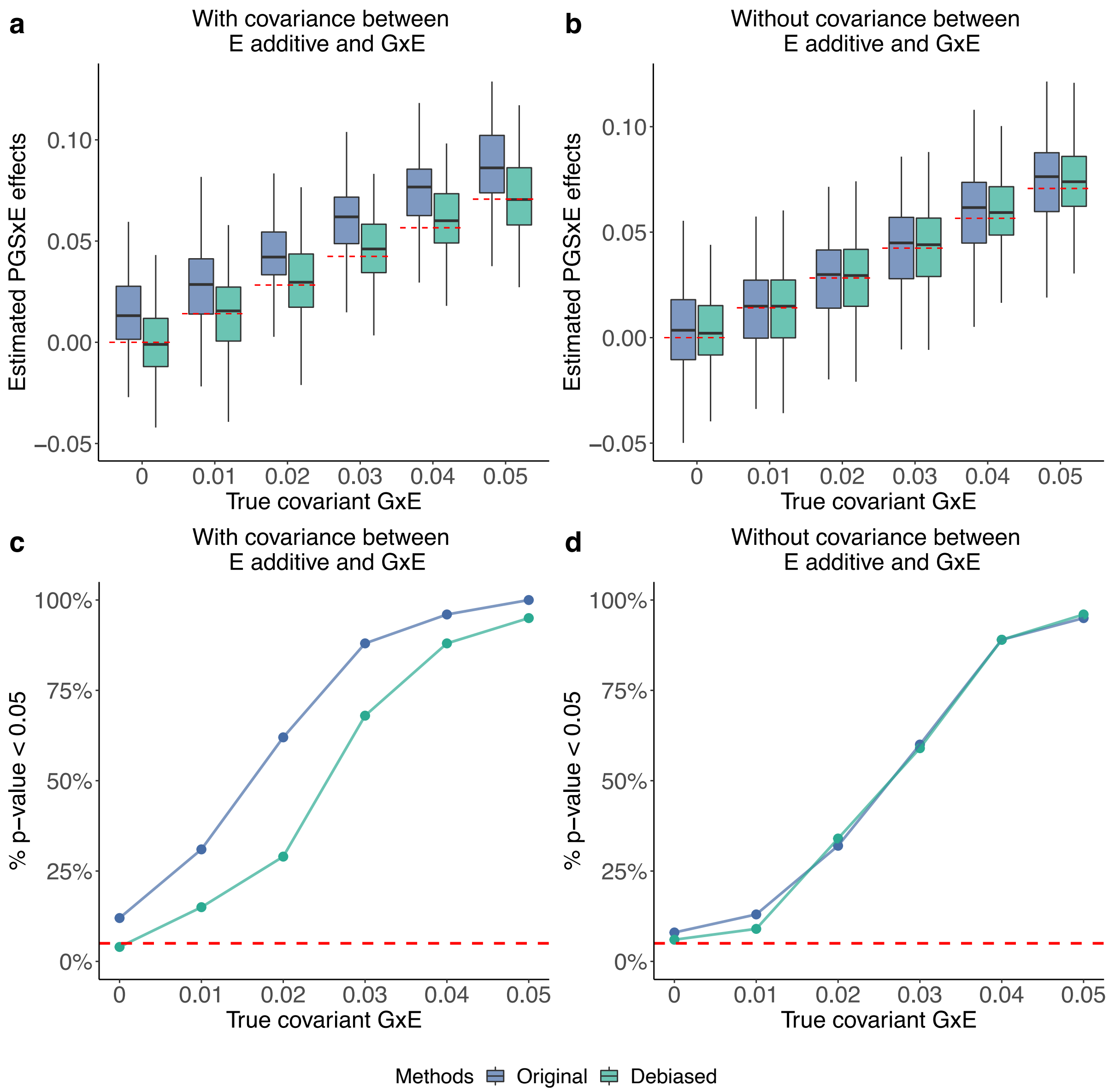


**Supplementary Figure 7. Simulation results of PIGEON for estimating oracle PGSxE in the presence of rGE. (a** and **c)** Simulation results when the additive genetic effects on the environment are uncorrelated with GxE interaction effects on the outcome trait. **(b** and **d)** Simulations results when the additive genetic effects on the environment and GxE interaction effects on the outcome trait are correlated. "Original" means using the methods derived under G-E independence.
