## Supplementary Note for "Reimagining Gene-Environment Interaction Analysis for Human Complex Traits"

### 1 Model

#### 1.1 PIGEON framework

PIGEON is built on a linear mixed model that captures both the polygenic additive effects and polygenic GxE effects for many SNPs:

$$Y_i = \sum_{j=1}^{M} G_{ij}\beta_{G_j} + E_i\beta_E + \sum_{j=1}^{M} G_{ij}E_i\beta_{I_j} + \epsilon_{i0} + \epsilon_{i1}E_i.$$

We rewrite into matrix form:

$$\underbrace{\boldsymbol{Y}}_{\text{Trait}} = \underbrace{\boldsymbol{G}\boldsymbol{\beta}_G}_{\text{Additive effects}} + \underbrace{\boldsymbol{E}\boldsymbol{\beta}_{\boldsymbol{E}}}_{\text{E's effects}} + \underbrace{\boldsymbol{E}\boldsymbol{G}\boldsymbol{\beta}_I}_{\text{GxE}} + \underbrace{\boldsymbol{\epsilon}_0}_{\text{Residual}} + \underbrace{\boldsymbol{E}\boldsymbol{\epsilon}_1}_{\text{RxE}} \tag{1}$$

where $\boldsymbol{Y}$ is a $N$-dimensional standardized trait vector, $\boldsymbol{G}$ are $N \times M$ standardized genotype matrix, $M$ is the number of SNPs, $\boldsymbol{E} = \text{diag}(\boldsymbol{e})$ is a $N \times N$ diagonal matrix, $\boldsymbol{e}$ is a $N$-dimensional vector for standardized environment, $\boldsymbol{\epsilon}_0$ is the $N$-dimensional residual effects vector, $\boldsymbol{E}\boldsymbol{\epsilon}_1$ is the $N$-dimensional vector quantifies the heteroskedasticity due to residual-environment interaction, $\boldsymbol{\beta}_G$ and $\boldsymbol{\beta}_I$ denote the genetic additive effects and GxE effects, $\boldsymbol{\beta}_E = [\beta_E, \cdots, \beta_E]^T$ is the $N$-dimensional vector representing environmental effects. We treat $\boldsymbol{\beta}_E$ as fixed and $\boldsymbol{\beta}_G, \boldsymbol{\beta}_I, \boldsymbol{\epsilon}_0, \boldsymbol{\epsilon}_1$ as random. We model all of these random variables as independent except for $(\boldsymbol{\beta}_G, \boldsymbol{\beta}_I)$ and $(\boldsymbol{\epsilon}_0, \boldsymbol{\epsilon}_1)$.

Suppose that $(\boldsymbol{\beta}_G, \boldsymbol{\beta}_I)$ have mean zero and covariance matrix below:

$$\text{Var}\left(\begin{bmatrix} \boldsymbol{\beta}_G \\ \boldsymbol{\beta}_I \end{bmatrix}\right) = \frac{1}{M}\begin{bmatrix} \sigma_G^2 I_M & \rho_{GI} I_M \\ \rho_{GI} I_M & \sigma_I^2 I_M \end{bmatrix}$$

where $\sigma_G^2$ and $\sigma_I^2$ quantify the variance explained by genetic additive effects and GxE effects. We further denote the $r_{GI} = \frac{\rho_{GI}}{\sqrt{\sigma_G^2 \sigma_I^2}}$. We also assume $(\epsilon_{0i}, \epsilon_{1i})$ have mean zero and covariance matrix below:

$$\text{Var}\left(\begin{bmatrix} \epsilon_{0i} \\ \epsilon_{1i} \end{bmatrix}\right) = \begin{bmatrix} \sigma_{\epsilon_0}^2 & \rho_{\epsilon_0,\epsilon_1} \\ \rho_{\epsilon_0,\epsilon_1} & \sigma_{\epsilon_1}^2 \end{bmatrix}$$

where $\sigma_{\epsilon_0}^2$ and $\sigma_{\epsilon_1}^2$ quantify the variance of residual and RxE. We further denote the $r_{\epsilon_0,\epsilon_1} = \frac{\rho_{\epsilon_0,\epsilon_1}}{\sqrt{\sigma_{\epsilon_0}^2 \sigma_{\epsilon_1}^2}}$.

We note that our model can be rewritten to include a single error term whose variance differs across environment:

$$Y_i = \sum_{j=1}^{M} G_{ij}\beta_{G_j} + E_i\beta_E + \sum_{j=1}^{M} G_{ij}E_i\beta_{I_j} + \epsilon_i,$$

where the marginal variance of the single error term is $\text{Var}(\epsilon_i) = \sigma_{\epsilon_0}^2 + \sigma_{\epsilon_1}^2$, and the conditional variance of the single error given the environment $E_i$ is $\text{Var}(\epsilon_i | E_i) = \sigma_{\epsilon_0}^2 + \sigma_{\epsilon_1}^2 E_i^2 + 2\rho_{\epsilon_0,\epsilon_1} E_i$.

#### 1.2 Notations when GWIS and GWAS are performed in different cohorts

Next, we describe the notations when there are two separate cohorts, one for GWIS (SNPxE) analysis with sample size $N_I$ and the other for GWAS with sample size $N_G$. We denote the PIGEON model in GWIS cohort with subscript $(I)$ and GWAS cohort with superscript $(G)$ as:

$$\begin{aligned} \boldsymbol{Y}^{(I)} &= \boldsymbol{G}^{(I)}\boldsymbol{\beta}_G + \boldsymbol{E}^{(I)}\boldsymbol{\beta}_{\boldsymbol{E}} + \boldsymbol{E}^{(I)}\boldsymbol{G}^{(I)}\boldsymbol{\beta}_I + \boldsymbol{\epsilon}_0^{(I)} + \boldsymbol{E}^{(I)}\boldsymbol{\epsilon}_1^{(I)} \\ \boldsymbol{Y}^{(G)} &= \boldsymbol{G}^{(G)}\boldsymbol{\beta}_G + \boldsymbol{E}^{(G)}\boldsymbol{\beta}_{\boldsymbol{E}} + \boldsymbol{E}^{(G)}\boldsymbol{G}^{(G)}\boldsymbol{\beta}_I + \boldsymbol{\epsilon}_0^{(G)} + \boldsymbol{E}^{(G)}\boldsymbol{\epsilon}_1^{(G)}, \end{aligned} \tag{2}$$

where we assume there are $N_S$ individuals who are included in both cohorts.

### 2 Comparing polygenic GxE methods in the PIGEON framework

In sections 2.1 and 2.2, we first consider a special case where the environment is binary to unify the differential heritability (genetic variance) analysis and the test that assesses whether genetic correlation is below 1 across the environments.

Suppose the raw-scale environment is a Bernoulli random variable with the probability of $p$ for being 1 and $1-p$ for being 0. After standardization to have a mean of 0 and a variance of 1, we have

$$\mathbb{P}(E_i = \sqrt{\frac{1-p}{p}}) = p \text{ and } \mathbb{P}(E_i = -\sqrt{\frac{p}{1-p}}) = 1-p$$

Then, the PIGEON model (1) in each environment can be written as

$$\begin{aligned} Y_i|(E_i = \sqrt{\frac{1-p}{p}}) &= \sum_{j=1}^{M} G_{ij}(\beta_{Gj} + \sqrt{\frac{1-p}{p}}\beta_{Ij}) + \sqrt{\frac{1-p}{p}}\beta_E + \epsilon_{0i} + \sqrt{\frac{1-p}{p}}\epsilon_{1i} \\ Y_i|(E_i = -\sqrt{\frac{p}{1-p}}) &= \sum_{j=1}^{M} G_{ij}(\beta_{Gj} - \sqrt{\frac{p}{1-p}}\beta_{Ij}) - \sqrt{\frac{p}{1-p}}\beta_E + \epsilon_{0i} - \sqrt{\frac{p}{1-p}}\epsilon_{1i} \end{aligned} \tag{3}$$

Based on this PIGEON model with a standardized binary environment, we have the following propositions about what differential heritability (genetic variance) and genetic correlation < 1 analysis target.

#### 2.1 Differential heritability

**Proposition 1.** *Under the PIGEON model with standardized binary environment (3), the heritability difference between two populations with $E_i = \sqrt{\frac{1-p}{p}}$ and $E_i = -\sqrt{\frac{p}{1-p}}$ is equal to*

$$\frac{\sqrt{\sigma_I^2\sigma_{\epsilon_0}^2}}{W}\left[(1-2p)\sqrt{\sigma_I^2\sigma_{\epsilon_0}^2} + 2\sqrt{p(1-p)}(r_{GI}\sqrt{\sigma_G^2\sigma_{\epsilon_0}^2} - r_{\epsilon_0,\epsilon_1}\sqrt{\sigma_I^2\sigma_{\epsilon_1}^2})\right] +$$
$$\frac{\sqrt{\sigma_G^2\sigma_{\epsilon_1}^2}}{W}\left[(2p-1)\sqrt{\sigma_G^2\sigma_{\epsilon_1}^2} + 2\sqrt{p(1-p)}(r_{GI}\sqrt{\sigma_I^2\sigma_{\epsilon_1}^2} - r_{\epsilon_0,\epsilon_1}\sqrt{\sigma_G^2\sigma_{\epsilon_0}^2})\right],$$

*where $W = p(1-p) \times \mathrm{Var}\left(Y_i \mid E_i = \sqrt{\frac{1-p}{p}}\right) \times \mathrm{Var}\left(Y_i \mid E_i = \sqrt{\frac{1-p}{p}}\right)$ is a scaling factor related to the phenotypic variance and mean of the environment.*

*Proof.* See Proof of Proposition 1 in Appendix. □

This shows that the heritability difference can be decomposed into two components. One comes from the product of the residual variance $\sigma^2_{\epsilon_0}$ and GxE variance components $\sigma^2_I$, and the other comes from the product of heteroskedasticity variance $\sigma^2_{\epsilon_1}$ and SNP-based heritability $\sigma^2_G$. Therefore, the heritability could still differ between two environments due to heteroskedasticity $\left(\sigma^2_{\epsilon_1} > 0\right)$ even without any GxE $\left(\sigma^2_I = 0\right)$.

#### 2.2 Differential genetic variance

**Proposition 2.** *Under the PIGEON model with standardized binary environment* (3)*, the genetic variance difference between two populations with* $E_i = \sqrt{\frac{1-p}{p}}$ *and* $E_i = -\sqrt{\frac{p}{1-p}}$ *is equal to*

$$\sqrt{\sigma^2_I}\left[(1-2p)\sqrt{\sigma^2_I} + 2r_{GI}\sqrt{p(1-p)\sigma^2_G}\right]$$

*Proof.* See Proof of Proposition 2 in Appendix. □

This is almost identical to the first components in heritability difference except for the residual variance $\sigma^2_{\epsilon_0}$ term. It indicates that the genetic variance difference ignores the impact of heteroskedasticity. However, if the GxE variance components $\sigma^2_I > 0$ (i.e, there exists GxE effects), the genetic variance can still be the same in two environment *i.e.,* $\left[(1-2p)\sqrt{\sigma^2_I} + \; 2r_{GI}\sqrt{p(1-p)\sigma^2_G}\right] = 0$. Therefore, differential genetic variance analysis could lead to false negative results.

#### 2.3 Genetic correlation < 1

**Proposition 3.** *Under the PIGEON model with standardized binary environment* (3)*, the genetic correlation = 1 between two populations with* $E_i = \sqrt{\frac{1-p}{p}}$ *and* $E_i = -\sqrt{\frac{p}{1-p}}$ *if and only if*

$$\sigma^2_I\sigma^2_G(r^2_{GI} - 1) = 0$$

*Proof.* See Proof of Proposition 3 in Appendix. □

Therefore, even if $\sigma^2_I > 0$, the genetic correlation between two environments can still be 1 if the SNP additive and SNPxE effects are perfectly correlated (i.e, $r_{GI} = \pm 1$ ). This indicates that test genetic correlation $= 1$ may have false negative results for testing the existence of any GxE.

#### 2.4 Assessing both equal genetic variance and perfect genetic correlation

**Proposition 4.** *A sufficient condition for* $\sigma^2_I > 0$ *to hold under both equal genetic variance and perfect genetic correlation, i.e.,*

$$\sigma^2_I\sigma^2_G(r^2_{GI} - 1) = 0 \text{ and } \sqrt{\sigma^2_I}\left[(1-2p)\sqrt{\sigma^2_I} + 2r_{GI}\sqrt{p(1-p)\sigma^2_G}\right] = 0,$$

*is*

$$r_{GI} = \pm 1, \sigma_G^2 > 0, \textit{ and } \sigma_I^2 = \frac{4p(1-p)\sigma_G^2}{(1-2p)^2}$$

*Proof.* This can be proven by plugging in $r_{GI} = \pm 1$ into both equations above. □

Therefore, even comparing both genetic variance and perfect genetic correlation between environments may fail to identify GxE.

#### 2.5 Oracle PGSxE

Next, we compare the PGSxE analysis in the PIGEON framework. Here, we don't require the raw environment to be binary. For illustration, we assume G-E independence (*i.e.*, the environment has zero heritability), but later we will relax this assumption and investigate the effect of gene-environment correlation (rGE) on polygenic GxE inference.

**Condition 1** (G-E independence)**.** *The environment has zero heritability.*

We then come to our main theorem showing that normalizing covariant GxE yields oracle PGSxE.

**Theorem 1** (Normalizing covariant GxE yields oracle PGSxE)**.** *Consider the oracle PGSxE regression*

$$Y_i \sim \alpha_G PGS_i + \alpha_E E_i + \alpha_I PGS_i E_i,$$

*where $PGS_i = \sum_j^M G_{ij}\beta_j$. Under the PIGEON model* (1) *and G-E independence assumption in condition 1, the oracle PGSxE interaction coefficient*

$$\alpha_I = \frac{\rho_{GI}}{\sigma_G^2},$$

*where $\rho_{GI}$ is the covariant GxE and $\sigma_G^2$ is the trait heritability.*

*Proof.* See Proof of Theorem 1 in Appendix. □

We further showed that the equivalence of hypothesis testing for between oracle PGSxE and covariant is invariant to the scale and location transformation of phenotype, PGS, and environment .

**Lemma 1** (Hypothesis testing for GxE is invariant to scale and location transformation)**.** *Given the arbitrary location and scale transformation of the trait, oracle PGS and environment, the interaction coefficient $\alpha_I^{(arb)}$ of oracle PGSxE regression*

$$Y_i^{(arb)} \sim \mu^{(arb)} + \alpha_G^{(arb)} PGS_i^{(arb)} + \alpha_E^{(arb)} E_i^{(arb)} + \alpha_I^{(arb)} PGS_i^{(arb)} E_i^{(arb)}$$

*can be transformed from the interaction regression coefficient described in theorem 1 by:*

$$\alpha_I^{(arb)} = \frac{\sqrt{\mathrm{Var}(Y_i^{(arb)})\sigma_G^2}}{\sqrt{\mathrm{Var}(PGS^{(arb)})\,\mathrm{Var}(E^{(arb)})}}\alpha_I$$

*Proof.* See Proof of Lemma 1 in Appendix. □

Therefore, we derived an estimate of the scale of interest implemented in our software and analysis:

**Corollary 1.** *The interaction coefficient in oracle PGSxE regression with standardized trait, standardized oracle PGS and raw scale environment for regression*

$$Y_i^{(std)} \sim \mu^{(std)} + \alpha_G^{(std)} PGS_i^{(std)} + \alpha_E^{(std)} E_i^{(raw)} + \alpha_I^{(std)} PGS_i^{(std)} E_i^{(raw)}$$

*is*

$$\alpha_I^{(std)} = \frac{\rho_{GI}}{\sqrt{\mathrm{Var}(E^{(raw)})\sigma_G^2}},$$

*where $\rho_{GI}$ is the covariant GxE, $\mathrm{Var}(E^{(raw)})$ is the variance of the raw-scale environment, ,and $\sigma_G^2$ is the additive heritability.*

*Proof.* This is easy to obtain by applying Lemma 1 in Theorem 1 □

### 3 The magnitude of bias for empirical PGSxE

In this section, we derive the bias due to the uncertainty in PGS and sample overlap between GWAS and PGSxE samples. We first define the empirical PGS calculated using estimated SNP weights, in contrast to the oracle PGS based on true SNP weights.

**Definition 1** (Empirical PGS)**.** *The i-th individual's empirical PGS calculated using discovery GWAS with sample size $N_G$ is defined as*

$$\widehat{PGS}_i^{(I)} = PGS_i^{(I)} + s_i,$$

*where $s_i$ represents the mean-zero estimation error in empirical PGS, $\mathrm{Cov}(PGS_i^{(I)}, s_i) = 0$, and $lim_{N_G \to \infty} s_i = 0$*

To better interpret the magnitude of the uncertainty for PGS, we followed Daetwyler et al. (2008) to have the following condition:

**Condition 2** (Magnitude of the uncertainty for PGS)**.** $\mathrm{Var}(s_i) = \frac{M}{N_G}$. *Here, $M$ is the effective number of independent SNPs and $N_G$ is the sample size of the discovery GWAS used to calculate the PGS.*

#### 3.1 No sample overlap

To investigate the bias solely due to the uncertainty in PGS, we first consider the case where there is no sample overlap between the GWAS and PGSxE samples.

**Condition 3** (Empirical PGS with no sample overlap)**.** *The estimation error* $s_i$ *in empirical PGS for i-th individual is uncorrelated with the trait* $Y_i^{(I)}$ *in the PGSxE sample*

$$\mathrm{Cov}(Y_i^{(I)}, s_i) = 0$$

**Theorem 2** (Bias in empirical PGSxE due to noisy PGS)**.** *Under the PIGEON model* (1)*, G-E independence assumption in condition 1, magnitude of the uncertainty for PGS in condition 2, and empirical PGS with no sample overlap in condition 3, the empirical PGSxE interaction coefficient for regression*

$$Y_i^{(I)} \sim \alpha_G^{(Emp)} \widehat{PGS}_i^{(I)} + \alpha_E^{(Emp)} E_i^{(I)} + \alpha_I^{(Emp)} \widehat{PGS}_i^{(I)} E_i^{(I)}$$

*is*

$$\alpha_I^{(Emp)} = \alpha_I + \frac{\sqrt{\sigma_G^2}(\sqrt{\sigma_G^2} - \sqrt{\sigma_G^2 + \frac{M}{N_G}})}{\sqrt{(\sigma_G^2 + \frac{M}{N_G})}} \alpha_I,$$

*Proof.* See Proof of Theorem 2 in Appendix. □

If the SNP additive heritability is non-zero (i.e. $\sigma_G^2 > 0$, which is true for most complex traits), we have $\sqrt{\sigma_G^2}(\sqrt{\sigma_G^2} - \sqrt{\sigma_G^2 + \frac{M}{N_G}}) < 0$. This indicates that the oracle PGSxE coefficient $\alpha_I$ represents an upper bound and infinity sample limit (GWAS sample size $N_G \to +\infty$) of the empirical PGSxE coefficient $\alpha_I^{(Emp)}$.

Moreover, denote the least squares estimator for empirical PGSxE interaction coefficient as $\hat{\alpha}_I^{(Emp)}$, by the property of least squares estimator, we have $\mathbb{E}[\hat{\alpha}_I^{(Emp)}] = \alpha_I^{(Emp)}$. Thus, $\hat{\alpha}_I^{(Emp)}$ is a biased estimate for oracle PGSxE coefficient $\alpha_I$ unless the discovery GWAS sample size is infinity or the SNP additive heritability is zero.

#### 3.2 With sample overlap

We then investigate the role of sample overlap in empirical PGSxE analysis.

**Condition 4** (Empirical PGS with sample overlap)**.** *The estimation error* $s_i$ *in empirical PGS for i-th individual is correlated with the trait* $Y_i^{(I)}$

$$\mathrm{Cov}(Y_i^{(I)}, s_i) \neq 0$$

**Theorem 3** (Bias in empirical PGSxE due to noisy PGS and sample overlap)**.** *Under the PIGEON model* (1)*, G-E independence assumption in condition 1, magnitude of the uncertainty for PGS in ondition 2, and empirical PGS with sample overlap in condition 4, the empirical PGSxE interaction coefficient for regression*

$$Y_i^{(I)} \sim \alpha_G^{(Emp,Ovp)} \widehat{PGS}_i^{(I)} + \alpha_E^{(Emp,Ovp)} E_i^{(I)} + \alpha_I^{(Emp,Ovp)} \widehat{PGS}_i^{(I)} E_i^{(I)},$$

*is*

$$\alpha_I^{(Emp,Ovp)} = \alpha_I \left[1 + \frac{\sqrt{\sigma_G^2}(\sqrt{\sigma_G^2} - \sqrt{\sigma_G^2 + \frac{M}{N_G}})}{\sqrt{(\sigma_G^2 + \frac{M}{N_G})}}\right](1 + \frac{2MN_S}{N_I N_G}) + \frac{MN_S(2\rho_{\epsilon_0,\epsilon_1} + \beta_E^2 \mu_E(3))}{N_I N_G(\frac{M}{N_G} + \sigma_G^2)}$$

*Proof.* See Proof of Theorem 3 in Appendix. □

Similarly, we expect the least squares estimator $\mathbb{E}[\hat{\alpha}_I^{(Emp,Ovp)}] = \alpha_I^{(Emp)}$. Thus, $\hat{\alpha}_I^{(Emp,Ovp)}$ is a biased estimates for oracle PGSxE coefficient $\alpha_I$. Also, we may get false positives due to sample overlap since we have the additional term in $\alpha_I^{(Emp,Ovp)}$ that is not a linear function of $\alpha_I$.

### 4 Estimation using summary-level data

We first define two concepts for the estimation using summary-level data.

**Definition 2** (LD score). *The LD score (Bulik-Sullivan et al., 2015b) of variant $j$ is $\ell_j = \sum_k r_{jk}^2$, where $r_{jk} = \mathbb{E}[G_{ij}G_{ik}]$ and the summation is taken over all other SNPs .*

**Definition 3** (Z-score in GWAS and GWIS summary statistics). *Under the PIGEON model (1), consider the GWAS (additive) with sample size $N_G$ and GWIS (SNPxE) with sample size $N_I$, we have the z-score for j-th SNP in GWAS (additive) is*

$$z_{Gj} := \frac{(\boldsymbol{G}_j^{(G)})^T \boldsymbol{Y}^{(G)}}{\sqrt{N_G}}.$$

*The z-score for j-th SNP in GWIS (SNPxE) is*

$$z_{Ij} := \frac{(\boldsymbol{E}^{(I)}\boldsymbol{G}^{(I)})_j^T \boldsymbol{Y}^{(I)}}{C\sqrt{N_I}},$$

*where the superscripts $(G)$ and $(I)$ indicate the fact that GWAS and GWIS are performed in separate cohorts (2). $C = \sqrt{1 - \frac{\mathrm{z}_\mathrm{E}^2}{\mathrm{z}_\mathrm{E}^2 + \mathrm{N}_I - 2}}$ is a correction factor to correct for the influence of the environment main effects on the z-score approximation in the derivation, $\mathrm{z}_\mathrm{E}$ is z-score for environmental effects.*

**Note:** The GWAS z-scores $z_{Gj}$ are obtained from standard marginal analysis. There is no need to add interaction terms when obtaining these additive effect z-scores.

#### 4.1 GxE variance components

**Proposition 5** (PIGEON-LDSC for GxE variance components). *Under the PIGEON model (1) and G-E independence assumption (1), we have*

$$\mathbb{E}\left[z_{Ij}^2 \mid \ell_j\right] = \frac{N_I \sigma_I^2}{C^2 M}\ell_j + [1 + (\beta_E^2 + \sigma_{\epsilon_1}^2 + \sigma_I^2)(\mu_E(4) - 1)]\frac{1}{C^2},$$

*where $C = \sqrt{1 - \frac{\mathrm{z}_\mathrm{E}^2}{\mathrm{z}_\mathrm{E}^2 + \mathrm{N}_G - 2}}$ is a correction factor to correct for the influence of the environment main effects on the Z-score approximation in the derivation.*

*Proof.* See Proof of Proposition 4 in Appendix. □

#### 4.2 Covariant GxE and Oracle PGSxE

**Proposition 6** (Bivariate PIGEON-LDSC for covariant GxE and oracle PGSxE)**.** *Under the PIGEON model* (1) *and G-E independence assumption* (1)*, we have*

$$\mathbb{E}\left[z_{Ij}z_{Gj} \mid \ell_j\right] = \frac{\sqrt{N_I N_G}\rho_{GI}}{CM}\ell_j + \frac{N_S}{C\sqrt{N_I N_G}}(2\rho_{GI} + 2\rho_{\epsilon_0,\epsilon_1} + \beta_E^2\mu_E(3)),$$

where $C = \sqrt{1 - \frac{\mathrm{z}_\mathrm{E}^2}{\mathrm{z}_\mathrm{E}^2 + \mathrm{N}_G - 2}}$ is a correction factor to correct for the influence of the environment main effects on the Z-score approximation in the derivation.

*Proof.* See Proof of Proposition 5 in Appendix. □

**Note:** The GWAS z-scores $z_{Gj}$ are obtained from standard marginal analysis. There is no need to add interaction terms when obtaining these additive effect z-scores.

The oracle PGSxE can be estimated by normalizing the covariant GxE by SNP heritability as described in Theorem 1.

### 5 Extending covariant GxE

Next, we expand our theory of covariant GxE to multiple traits for hypothesis-free search of PGSxE.

#### 5.1 Multi-trait model

Consider two studies for two trait $\boldsymbol{Y}_1$ and $\boldsymbol{Y}_2$ with sample size $N_I$ and $N_G$, respectively, and assume the standardized traits $\boldsymbol{Y}_1$ and $\boldsymbol{Y}_2$ follow the linear models below:

$$\begin{aligned}
\underbrace{\boldsymbol{Y}_1}_{\text{Trait}} &= \underbrace{\boldsymbol{G}\boldsymbol{\beta}_G}_{\text{Additive effects}} + \underbrace{\boldsymbol{E}\boldsymbol{\beta}_{\boldsymbol{E}}}_{\text{Envir's effects}} + \underbrace{\boldsymbol{E}\boldsymbol{G}\boldsymbol{\beta}_I}_{\text{GxE}} + \underbrace{\boldsymbol{\epsilon}_0}_{\text{Residual}} + \underbrace{\boldsymbol{E}\boldsymbol{\epsilon}_1}_{\text{RxE}} \\
\underbrace{\boldsymbol{Y}_2}_{\text{Trait}} &= \underbrace{\boldsymbol{Z}\boldsymbol{\gamma}_G}_{\text{Additive effects}} + \underbrace{\boldsymbol{X}\boldsymbol{\gamma}_{\boldsymbol{E}}}_{\text{Envir's effects}} + \underbrace{\boldsymbol{X}\boldsymbol{Z}\boldsymbol{\gamma}_I}_{\text{GxE}} + \underbrace{\boldsymbol{\delta}_0}_{\text{Residual}} + \underbrace{\boldsymbol{X}\boldsymbol{\delta}_1}_{\text{RxE}}
\end{aligned} \tag{4}$$

Here, $\boldsymbol{G}$ and $\boldsymbol{Z}$ are $N_I \times M$ and $N_G \times M$ standardized genotype matrix, $M$ is the number of SNPs, $\boldsymbol{E} = \text{diag}(\boldsymbol{e})$ and $\boldsymbol{X} = \text{diag}(\boldsymbol{x})$ is the $N_I \times N_I$ and $N_G \times N_G$ matrix, $\boldsymbol{e}$ and $\boldsymbol{x}$ are the the $N_I$ and $N_G$-dimensional vector for the same environment in two studies, $\boldsymbol{\epsilon}_0$ and $\boldsymbol{\delta}_0$ are the $N_I$ and $N_G$-dimensional residual effects vector, $\boldsymbol{\epsilon}_1\boldsymbol{E}$ and $\boldsymbol{X}\boldsymbol{\delta}_1$is the $N_I$ and $N_G$-dimensional vector representing RxE effects vector. $\boldsymbol{\beta}_E = [\beta_E, \cdots, \beta_E]^T$ and $\boldsymbol{\gamma}_E = [\gamma_E, \cdots, \gamma_E]^T$ is the $N_I$ and $N_G$-dimensional vector representing environmental additive effects. $\boldsymbol{\beta}_G, \boldsymbol{\beta}_I$ and $\boldsymbol{\gamma}_G, \boldsymbol{\gamma}_I$ denotes the additive genetic effects and GxE effects in study 1, and additive genetic effects in study 2.

Suppose that $(\boldsymbol{\beta}_G, \boldsymbol{\beta}_I, \boldsymbol{\gamma}_G, \boldsymbol{\gamma}_I)$ have mean zero and covariance matrix below:

$$\text{Var}\left(\begin{bmatrix} \boldsymbol{\beta}_G \\ \boldsymbol{\beta}_I \\ \boldsymbol{\gamma}_G \\ \boldsymbol{\gamma}_I \end{bmatrix}\right) = \frac{1}{M}\begin{bmatrix} \sigma_G^2 I_M & \rho_{GI} I_M & \rho_{G1,G2} I_M & \rho_{G1,I2} I_M \\ \rho_{GI} I_M & \sigma_I^2 I_M & \rho_{G2,I} I_M & \rho_{I,I2} I_M \\ \rho_{G1,G2} I_M & \rho_{G2,I} I_M & \sigma_{G2}^2 I_M & \rho_{G2,I2} I_M \\ \rho_{G1,I2} I_M & \rho_{I,I2} I_M & \rho_{G2,I2} I_M & \sigma_{I2}^2 I_M \end{bmatrix}$$

where $\sigma_G^2, \sigma_I^2, \sigma_{G2}^2$ and $\sigma_{I2}^2$ are the variance explained by additive genetic effects and GxE effects for trait 1 and trait 2, respectively.

For $N_S$ individuals who have trait measured in both studies, we assume $(\epsilon_{0i}, \epsilon_{1i}, \delta_{0i}, \delta_{1i})$ for the $i$-th individual have a mean zero and covariance matrix:

$$\text{Var}\left(\begin{bmatrix} \epsilon_{0i} \\ \epsilon_{1i} \\ \delta_{0i} \\ \delta_{1i} \end{bmatrix}\right) = \frac{1}{M}\begin{bmatrix} \sigma_{\epsilon_0}^2 & \rho_{\epsilon_0,\epsilon_1} & \rho_{\epsilon_0,\delta_0} & \rho_{\epsilon_0,\delta_1} \\ \rho_{\epsilon_0,\epsilon_1} & \sigma_{\epsilon_1}^2 & \sigma_{\epsilon_1,\delta_0} & \rho_{\epsilon_1,\delta_0} \\ \sigma_{\epsilon_0,\delta_0} & \sigma_{\epsilon_1,\delta_0} & \sigma_{\delta_0}^2 & \sigma_{\delta_0,\delta_1} \\ \sigma_{\epsilon_0,\delta_1} & \rho_{\epsilon_1,\delta_1} & \sigma_{\delta_0,\delta_1} & \sigma_{\delta_1}^2 \end{bmatrix}$$

#### 5.2 Multi-trait Oracle PGSxE

**Corollary 2** (Multi-trait oracle PGSxE and covariant GxE)**.** *Consider the oracle PGSxE regression*

$$Y_{1i} \sim \alpha_G^{(PGS2)} PGS_{2i} + \alpha_E^{(PGS2)} E_i + \alpha_I^{(PGS2)} PGS_{2i} E_i,$$

*where $Y_{1i}$ is the trait 1, $PGS_{2i} = \sum_j^M G_{ij}\gamma_j$ is the PGS for trait 2. Under the multi-trait PIGEON model (4) and G-E independence assumption in condition 1, the multi-trait oracle PGSxE interaction coefficient*

$$\alpha_I^{(PGS2)} = \frac{\rho_{G2,I}}{\sigma_{G2}^2},$$

*where $\rho_{GI}$ is the covariant GxE and $\sigma_G^2$ is the heritability for trait 2.*

*Proof.* This can be obtained by modifying the Proof of Theorem 1 in the Appendix into two traits case. □

#### 5.3 Multi-trait bivariate PIGEON-LDSC

**Proposition 7** (Bivariate multi-trait PIGEON-LDSC)**.** *Under the multi-trait PIGEON model (4) and G-E independence assumption in condition 1, we have*

$$\mathbb{E}\left[z_{Ij} z_{G2j} \mid \ell_j\right] = \frac{\sqrt{N_I N_{G2}}\rho_{G2,I}}{CM}\ell_j + \frac{N_{S2}}{C\sqrt{N_I N_{G2}}}(\rho_{G2,I} + \rho_{G,I2} + \rho_{\epsilon_0,\delta_1} + \rho_{\epsilon_1,\delta_0} + \beta_E\gamma_E\mu_E(3)),$$

*where $C = \sqrt{1 - \frac{\mathrm{z_E^2}}{\mathrm{z_E^2} + \mathrm{N}_G - 2}}$ is a correction factor to correct for the influence of the environment main effects on the Z-score approximation in the derivation.*

*Proof.* This can be obtained by modifying the Proof of Proposition 5 in the Appendix into two traits case. □

**Note:** The GWAS z-scores $z_{G2j}$ are obtained from standard marginal analysis. There is no need to add interaction terms when obtaining these additive effect z-scores.

The multi-trait oracle PGSxE can be estimated by normalizing the covariant GxE by SNP heritability described in Theorem 1.

##### 5.4 Note about GWAS summary statistics

#### 6 Some additional features of PIGEON

##### 6.1 Binary trait in PIGEON

Here, we demonstrate that the PIGEON framework can be applied to binary traits in case/control studies. We assume the liability threshold model, where the binary trait is determined by continuous liability $\zeta$, i.e. $\eta = 1[\zeta > \tau]$ and $\tau$ is the liability threshold. The liability threshold $\tau$ is determined by $\tau = \Phi^{-1}(1-K)$, where $\Phi$ is the standard normal CDF and K is the population prevalence. As an analogy of the PIGEON model (1) for the quantitative trait, we model continuous liability by

$$\underbrace{\boldsymbol{\zeta}}_{\text{Liability}} = \underbrace{\boldsymbol{G}\boldsymbol{\beta}_G^{(lia)}}_{\text{Additive effects}} + \underbrace{\boldsymbol{E}\boldsymbol{\beta}_E^{(lia)}}_{\text{E's effects}} + \underbrace{\boldsymbol{E}\boldsymbol{G}\boldsymbol{\beta}_I^{(lia)}}_{\text{GxE}} + \underbrace{\boldsymbol{\epsilon}_0}_{\text{Residual}} + \underbrace{\boldsymbol{E}\boldsymbol{\epsilon}_1}_{\text{RxE}} \tag{5}$$

We follow the PIGEON model for quantitative trait (1) and assumes that $(\boldsymbol{\beta}_G, \boldsymbol{\beta}_I)$ have mean zero and covariance matrix below:

$$\text{Var}\left(\begin{bmatrix} \boldsymbol{\beta}_G^{(lia)} \\ \boldsymbol{\beta}_I^{(lia)} \end{bmatrix}\right) = \frac{1}{M}\begin{bmatrix} {\sigma_G^{(lia)}}^2 I_M & \rho_{GI}^{(lia)} I_M \\ \rho_{GI}^{(lia)} I_M & {\sigma_I^{(lia)}}^2 I_M \end{bmatrix}$$

where ${\sigma_G^{(lia)}}^2$, ${\sigma_I^{(lia)}}^2$, and $\rho_{GI}^{(lia)}$ are the variance explained by genetic additive effects, GxE effects, and covariant GxE in the liability scale.

We first consider how to transform the liability covariant GxE and additive heritability from the observed scale results:

**Proposition 8** (Transformation between liability scale and observed scale)**.** *The liability scale results can be transformed from the observed scale results by*

$$\begin{aligned} {\sigma_G^{(obs)}}^2 &= {\sigma_G^{(lia)}}^2 \left(\frac{\Phi(\tau)^2 P(1-P)}{K^2(1-K)^2}\right) \\ {\sigma_I^{(obs)}}^2 &= {\sigma_I^{(lia)}}^2 \left(\frac{\Phi(\tau)^2 P(1-P)}{K^2(1-K)^2}\right) \\ \rho_{GI}^{(obs)} &= \rho_{GI}^{(lia)} \left(\frac{\Phi(\tau)^2 P(1-P)}{K^2(1-K)^2}\right) \end{aligned}$$

*where $\Phi$ is the standard normal density, $K$ is the population prevalence, $P$ is the sample prevalence, and $\tau$ is the liability threshold.*

*Proof.* This can be proven by following the original bivariate LDSC paper (Bulik-Sullivan et al., 2015a). □

Next, we consider the relationship between covariant GxE and the oracle PGSxE under the liability model.

**Proposition 9** (Oracle PGSxE in liability model)**.** *Consider the oracle PGSxE regression on the liability scale*

$$\zeta_i \sim \alpha_G^{(lia)} PGS_i + \alpha_E^{(lia)} E_i + \alpha_I^{(lia)} PGS_i E_i,$$

*where* $PGS_i = \sum_j^M G_{ij}\beta_j$*. Under the PIGEON liability model (5) and G-E independence assumption in condition 1, the oracle PGSxE interaction coefficient*

$$\alpha_I^{(lia)} = \frac{\rho_{GI}^{(lia)}}{\sigma_G^{(lia)2}},$$

*where* $\rho_{GI}^{(lia)}$ *is the covariant GxE and* $\sigma_G^{(lia)2}$ *is the additive heritability on the liability scale. Furthermore,*

$$\alpha_I^{(lia)} = \frac{\rho_{GI}^{(obs)}}{\sigma_G^{(obs)2}},$$

*where* $\rho_{GI}^{(obs)}$ *is the covariant GxE and* $\sigma_G^{(obs)2}$ *is the additive heritability on the observed scale.*

*Proof.* This can be shown by following the proof for quantitative traits and the transformation stated in proposition (8). □

**Note:** Directly applying PIGEON-LDSC on GWIS and GWAS summary statistics yields the estimates on the observed scale.

We then list several possible cases about how to transform the covariant GxE to oracle PGSxE when the binary trait is involved.

**Corollary 3** (Multi-trait oracle PGSxE under liability model)**.** *Consider these oracle PGSxE regression*

- *with liability* $\zeta_i$ *as outcome and PGS for a quantitative traits* $Y_{2i}$

$$\zeta_i \sim \alpha_G^{(PGS2,lia)} PGS_i + \alpha_E^{(PGS2,lia)} E_i + \alpha_I^{(PGS2,lia)} PGS_{2i} E_i,$$

*where* $PGS_i = \sum_j^M G_{ij}\gamma_j$*. Under the PIGEON liability model (5), quantitative model for* $Y_{2i}$ *and G-E independence assumption in condition 1, the oracle PGSxE interaction coefficient*

$$\alpha_I^{(PGS2,lia)} = \frac{\rho_{G2,I}\left(\frac{K(1-K)}{\Phi(\tau)\sqrt{P(1-P)}}\right)}{\sigma_{G2}^2}.$$

- *with quantitative trait* $Y_{1i}$ *as outcome and PGS for a liability traits* $\zeta_{2i}$

$$Y_{1i} \sim \alpha_G^{(PGS2,quan)} PGS_i + \alpha_E^{(PGS2,quan)} E_i + \alpha_I^{(PGS2,quan)} PGS_{2i} E_i,$$

*where* $PGS_{2i} = \sum_j^M G_{ij}\gamma_j^{(lia)}$. *Under the PIGEON liability model* (5)*, quantitative model for* $Y_{1i}$ *and G-E independence assumption in condition 1, the oracle PGSxE interaction coefficient*

$$\alpha_I^{(PGS2,quan)} = \frac{\rho_{G,I2}}{\sigma^2_{G2}\left(\frac{K(1-K)}{\Phi(\tau)\sqrt{P(1-P)}}\right)}.$$

- *liability* $\zeta_i$ *as outcome and PGS for a liability traits* $\zeta_{2i}$ *for*

$$Y_{1i} \sim \alpha_G^{(PGS2,quan)} PGS_i + \alpha_E^{(PGS2,quan)} E_i + \alpha_I^{(PGS2,quan)} PGS_{2i}E_i,$$

*where* $PGS_{2i} = \sum_j^M G_{ij}\gamma_j^{(lia)}$*, and*

$$\underbrace{\boldsymbol{\zeta}_2}_{\text{Liarbility}} = \underbrace{\boldsymbol{Z}\boldsymbol{\gamma}_G^{(lia)}}_{\text{Additive effects}} + \underbrace{\boldsymbol{X}\boldsymbol{\gamma}_E^{(lia)}}_{\text{Envir's effects}} + \underbrace{\boldsymbol{X}\boldsymbol{Z}\boldsymbol{\gamma}_I^{(lia)}}_{\text{GxE}} + \underbrace{\boldsymbol{\delta}_0}_{\text{Residual}} + \underbrace{\boldsymbol{X}\boldsymbol{\delta}_1}_{\text{RxE}}$$

*Under the PIGEON liability model* (5)*, quantitative model for* $Y_{1i}$ *and G-E independence assumption in condition 1, the oracle PGSxE interaction coefficient*

$$\alpha_I^{(PGS2,quan)} = \frac{\rho_{G,I2}\left(\frac{K_1(1-K_1)}{\Phi(\tau_1)\sqrt{P_1(1-P_1)}}\right)}{\sigma^2_{G2}\left(\frac{K_2(1-K_2)}{\Phi(\tau_2)\sqrt{P_2(1-P_2)}}\right)},$$

*where* $\Phi$ *is the standard normal density,* $K_1$ *and* $K_2$ *are the population prevalence,* $P_1$ *and* $P_2$ *are the sample prevalence, and* $\tau_1$ *and* $tau_2$ *are the liability threshold of binary trait 1 and trait 2.*

*Proof.* This can be shown by following the proof for quantitative traits and the transformation stated in proposition (8). □

#### 6.2 (Dichotomized PGS)-by-E interactions

Next, we demonstrate that the PIGEON framework can be applied to (Dichotomized PGS)-by-E interactions. Here, the (Dichotomized PGS)-by-E interactions mean the interaction between a binary variable depends on the percentile of PGS distribution and the environment.

**Proposition 10** (Dichotomized PGS-by-E interactions)**.** *Consider the oracle Dichotomized PGS-by-E regression*

$$Y_i \sim \alpha_G^{(dicho,q)} \mathbb{1}(PGS_i > q) + \alpha_E^{(dicho,q)} E_i + \alpha_I^{(dicho,q)} \mathbb{1}(PGS_i > w)E_i,$$

*where* $PGS_i = \sum_j^M G_{ij}\beta_j$ *and*

$$w = \inf\{x \in \mathbb{R} : 1 - q \leq F_{\boldsymbol{G}_i\boldsymbol{\beta}_G}(x)\} = \sqrt{\sigma^2_G}\sqrt{2}\,\mathrm{erf}^{-1}(1-2q),$$

*denotes the pre-specified threshold such that the individuals in the top* $q$ *of the PGS distribution are coded as 1 and others are coded as 0.* erf *is the error function defined as* $\mathrm{erf}(x) = \frac{2}{\sqrt{\pi}}\int_0^x e^{-t^2}dt$. *Under the PIGEON model* (1) *and G-E independence assumption in condition 1, the oracle Dichotomized PGS-by-E coefficient*

$$\alpha_I^{(dicho,q)} = \frac{\rho_{GI}\Phi\left(\sqrt{2}\,\mathrm{erf}^{-1}(1-2q)\right)}{q\sqrt{\sigma^2_G}}$$

*where $\rho_{GI}$ is the covariant GxE, $\sigma_G^2$ is the additive heritability, and $\Phi$ denotes the standard normal density function.*

*Proof.* See Proof of Proposition 9 in Appendix. □

#### 6.3 PIGEON conditional oracle PGSxE analysis

Next, we consider the PIGEON conditional oracle PGSxE analysis, representing the oracle PGSxE regression with multiple PGS and their interactions with the environment as predictors. We only prove the case with two PGS, but the results for more than two scores is easy to generalize and have been implemented in the software.

**Proposition 11** (Conditional PIGEON analysis)**.** *Consider the oracle PGSxE regression with two standardized PGS for two different traits*

$$\boldsymbol{Y} \sim \boldsymbol{E}\alpha_E^{(cond)} + \boldsymbol{PGS}^{(std)}\alpha_G^{(PGS,cond)} + \boldsymbol{PGS}_2^{(std)}\alpha_G^{(PGS2,cond)} + \boldsymbol{EPGS}^{(std)}\alpha_I^{(PGS,cond)} + \boldsymbol{EPGS}_2^{(std)}\alpha_I^{(PGS2,cond)}.$$

*Under the multi-trait PIGEON model* (4) *and G-E independence assumption in condition 1, the oracle PGSxE interaction coefficients are*

$$\begin{bmatrix} \alpha_I^{(PGS,cond)} \\ \alpha_I^{(PGS2,cond)} \end{bmatrix} = \begin{bmatrix} 1 & r_{G,G2} \\ r_{G,G2} & 1 \end{bmatrix}^{-1} \begin{bmatrix} \frac{\rho_{GI}}{\sqrt{\sigma_G^2}} \\ \frac{\rho_{G2,I}}{\sqrt{\sigma_{G2}^2}} \end{bmatrix},$$

*where $r_{G,G2}$ is the genetic correlation between these two traits.*

*Proof.* See Proof of Proposition 10 in Appendix. □

#### 6.4 Interpretation of the direction of the oracle PGSxE

Interpretation of the direction of the coefficient for GxE interaction depends on the coding of G and E. Here we mainly discuss how to interpret the direction for oracle PGSxE using genetic covariance and covariant GxE in PIGEON. Consider the oracle PGSxE regression of interest:

$$Y_i^{(std)} \sim \mu^{(std)} + \alpha_G^{(std)} PGS_i^{(std)} + \alpha_E^{(std)} E_i^{(raw)} + \alpha_I^{(std)} PGS_i^{(std)} E_i^{(raw)},$$

where $Y_i^{(std)}$ is the standardized trait, $PGS_i^{(std)}$ is the standardized oracle PGS and $E_i^{(raw)}$ is the raw scale environment. Then we have

$$\alpha_I^{(std)} = \frac{\rho_{GI}}{\sqrt{\text{Var}(E^{(raw)})\sigma_G^2}}$$
$$\alpha_G^{(std)} = \sqrt{\sigma_G^2} - \alpha_I^{(std)}\mathbb{E}[E_i^{(raw)}],$$

where $\mathbb{E}[E_i^{(raw)}]$ is the expectation of the raw environment

If we have multi-trait oracle PGSxE, we have

$$\alpha_I^{(std)} = \frac{\rho_{G2,I}}{\sqrt{\mathrm{Var}(E^{(raw)})\sigma_{G2}^2}}$$
$$\alpha_G^{(std)} = \frac{\rho_{G1,G2}}{\sqrt{\sigma_{G2}^2}} - \alpha_I^{(std)}\mathbb{E}[E_i^{(raw)}],$$

### 7 Comparing with several other estimation methods

#### 7.1 GxEsum for GxE variance components estimation

Shin and Lee (2021) proposed GxEsum for estimation GxE variance components (but not covariant GxE) with GWIS summary statistics. It is similar to PIGEON-LDSC but we account for the environment's main effects and G-E correlations that will be described in section 8. In terms of environmental main effect, GxEsum requires the GWIS to be performed on phenotypic residual that is pre-adjusted for covariates and the main effect of environment (i.e., marginal regression between SNP and phenotypic residual). We showed in simulation that directly applying their methods to the commonly applied multiple regression in GWIS would lead to biased results (**Supplementary Figure 2**). Applying our correction factor in the regression provides an unbiased estimate. We note that this phenomenon is related to the "marginal" and "conditional" variance for heritability estimation described in (Weissbrod et al., 2018) .We further proposed the correction factor to account for the impact of covariates (such as sex and age) in addition to the environment main effect in GWIS in section 9.2.

In terms for rGE, Shin and Lee (2021) demonstrate GxEsum is robust to rGE by simulation with the covariance between additive effect on the environment and SNPxE interaction effect on the trait outcome set to 0. In our main texts, we showed that this is exactly the sufficient condition to have unbiased results for GxE variance components. In our simulations, we showed that GxEsum provides biased estimates when this condition is violated, while PIGEON remains unbiased.

#### 7.2 Measurement error correction for PGSxE

Becker et al. (2021) proposed a measurement-error-corrected estimator for PGSxE regressions. We compared it with PIGEON in simulations described in section 11.1 and found that it has false positive results and a larger standard error than PIGEON estimates (**Supplementary Figure 3**). We suspect that the false positive is due to a numerical problem with the matrix inverse operation in the software when the PGS and environment have a correlation in the scale of $10^{-3}$. In addition to this limitation, this measurement error correction approach requires the individual-level phenotype data and genotype data to calculate the R-squared for PGS regression while PIGEON only needs summary statistics. Therefore, PIGEON is the superior approach in terms of statistical properties and data requirements.

### 8 Accounting for rGE in PIGEON

Next, we investigate the impact of rGE in polygenic GxE inference.

#### 8.1 Extending the PIGEON model to allow rGE

**Motivations.** The PIGEON model (1) assumes that

$$\underbrace{\boldsymbol{Y}}_{\text{Trait}} = \underbrace{\boldsymbol{G}\boldsymbol{\beta}_G}_{\text{Additive effects}} + \underbrace{\boldsymbol{E}\boldsymbol{\beta}_{\boldsymbol{E}}}_{\text{E's effects}} + \underbrace{\boldsymbol{E}\boldsymbol{G}\boldsymbol{\beta}_I}_{\text{GxE}} + \underbrace{\boldsymbol{\epsilon}_0}_{\text{Residual}} + \underbrace{\boldsymbol{E}\boldsymbol{\epsilon}_1}_{\text{RxE}}, \tag{6}$$

where $(\boldsymbol{\beta}_G, \boldsymbol{\beta}_I)$ have mean zero and covariance matrix below:

$$\text{Var}\left(\begin{bmatrix} \boldsymbol{\beta}_G \\ \boldsymbol{\beta}_I \end{bmatrix}\right) = \frac{1}{M}\begin{bmatrix} \sigma_G^2 I_M & \rho_{GI} I_M \\ \rho_{GI} I_M & \sigma_I^2 I_M \end{bmatrix}$$

The two main objectives in PIGEON are to estimate the GxE variance components $\sigma_I^2$ and the covariant GxE $\rho_{GI}$. Hypothetically, the two parameters should be defined with the exogenous environment (e.g. a randomized treatment, a policy change, a natural experiment) that is independent with genetics as well as all covariates for clear interpretation. Therefore, we have assumed the G-E independence in most derivations so far. However, environmental variables obtained from observational data can be "contaminated by" G-E correlation (rGE). Therefore, we aim to quantify such contamination in the data-generating model.

**PIGEON-rGE model.**

$$\begin{aligned} Y_i &= \sum_{j=1}^{M} G_{ij}\beta_{G_j} + V_i\beta_E + \sum_{j=1}^{M} G_{ij}V_i\beta_{I_j} + \epsilon_{i0} + V_i\epsilon_{i1} \\ V_i &= \sum_{j=1}^{M} G_{ij}\alpha_{G_j} + E_i\sqrt{(1-\sigma_E^2)}, \end{aligned}$$

where the $V_i$ is the standardized "contaminated" environment.

We rewrite into matrix form:

$$\begin{aligned} \boldsymbol{Y} &= \boldsymbol{\mu} + \boldsymbol{G}\boldsymbol{\beta}_G + \boldsymbol{V}\boldsymbol{\beta}_{\boldsymbol{E}} + \boldsymbol{V}\boldsymbol{G}\boldsymbol{\beta}_I + \boldsymbol{\epsilon}_0 + \boldsymbol{V}\boldsymbol{\epsilon}_1 \\ \boldsymbol{V} &= \text{diag}(\boldsymbol{v}) \\ \boldsymbol{v} &= \boldsymbol{G}\boldsymbol{\theta}_G + \sqrt{(1-\sigma_E^2)}\boldsymbol{e}, \end{aligned} \tag{7}$$

where $\boldsymbol{Y}$ is a $N$-dimensional standardized trait vector, $\boldsymbol{G}$ are $N \times M$ standardized genotype matrix, $M$ is the number of SNPs, $\boldsymbol{V}$ is the $N \times N$ matrix, $\boldsymbol{v}$ and $\boldsymbol{e}$ is the $N$-dimensional vector for the standardized "contaminated" and exogenous environment, $\boldsymbol{\epsilon}_0$ is the $N$-dimensional residual effects vector, $\boldsymbol{V}\boldsymbol{\epsilon}_1$ is the $N$-dimensional vector representing RxE effects. $\boldsymbol{\beta}_G$ and $\boldsymbol{\beta}_I$ denotes the genetic additive effects and GxE effects for the outcome trait $\boldsymbol{Y}$, $\boldsymbol{\beta}_E = [\beta_E, \cdots, \beta_E]^T$ is the $N$-dimensional vector representing environmental additive effects, $\boldsymbol{\theta}_G$ denotes the genetic additive effects for the standardized "contaminated" environment. We treat $\boldsymbol{\beta}_E$

as fixed and , $\boldsymbol{\epsilon}_0, \boldsymbol{\epsilon}_1, \boldsymbol{\beta}_G, \boldsymbol{\beta}_I, \boldsymbol{\theta}_G$ as random. We model all of these random variables as independent except for $(\boldsymbol{\beta}_G, \boldsymbol{\beta}_I, \boldsymbol{\theta}_G)$ and $(\boldsymbol{\epsilon}_0, \boldsymbol{\epsilon}_1)$.

Suppose that $(\boldsymbol{\beta}_G, \boldsymbol{\beta}_I, \boldsymbol{\theta}_G)$ and $(\boldsymbol{\epsilon}_0, \boldsymbol{\epsilon}_1)$ have mean zero and covariance matrix below:

$$\mathrm{Var}\begin{pmatrix}\boldsymbol{\beta}_G\\ \boldsymbol{\beta}_I\\ \boldsymbol{\theta}_G\end{pmatrix} = \frac{1}{M}\begin{pmatrix}\sigma^2_{GI}\boldsymbol{I}_M & \rho_{GI}\boldsymbol{I}_M & \rho_{GE}\boldsymbol{I}_M\\ \rho_{GI}\boldsymbol{I}_M & \sigma^2_I\boldsymbol{I}_M & \rho_{IE}\boldsymbol{I}_M\\ \rho_{GE}\boldsymbol{I}_M & \rho^2_{IE}\boldsymbol{I}_M & \sigma^2_E\boldsymbol{I}_M\end{pmatrix}, \mathrm{Var}\begin{pmatrix}\epsilon_{i0}\\ \epsilon_{i1}\end{pmatrix} = \begin{pmatrix}\sigma^2_{\epsilon_0} & \rho_{\epsilon_0,\epsilon_1}\\ \rho_{\epsilon_0,\epsilon_1} & \sigma^2_{\epsilon_1}\end{pmatrix}$$

where $\sigma^2_G$ and $\sigma^2_I$ are the variance explained by genetic additive effects and GxE effects for the trait $Y_i$, and $\sigma^2_E$ is the variance explained by genetic additive effects for the "contaminated" environment. We further assume that these parameters are constrained such that $\mathrm{Var}(Y_i) = 1$.

**Notations for two cohorts.**

Next, we describe the notations when there are two cohorts with two different traits for proofing multi-trait covariant GxE results. Denote the PIGEON model in the GWIS cohort for the first trait $\boldsymbol{Y}_1$ and GWAS model for the second trait $\boldsymbol{Y}_2$ as:

$$\begin{aligned}
&\boldsymbol{Y}_1 = \boldsymbol{\mu} + \boldsymbol{G}\boldsymbol{\beta}_G + \boldsymbol{V}\boldsymbol{\beta}_{\boldsymbol{E}} + \boldsymbol{V}\boldsymbol{G}\boldsymbol{\beta}_I + \boldsymbol{\epsilon}_0 + \boldsymbol{V}\boldsymbol{\epsilon}_1\\
&\boldsymbol{V} = \mathrm{diag}(\boldsymbol{v})\\
&\boldsymbol{v} = \boldsymbol{G}\boldsymbol{\theta}_G + \sqrt{(1-\sigma^2_E)}\boldsymbol{e}\\
&\boldsymbol{Y}_2 = \boldsymbol{Z}\boldsymbol{\gamma}_G + \boldsymbol{\delta},
\end{aligned} \tag{8}$$

where $\boldsymbol{Y}_1$ is a $N_I$-dimensional standardized trait vector, $\boldsymbol{Y}_2$ is a $N_G$-dimensional standardized trait vector. There are $N_S$ individuals who are included in both cohorts. The variance-covariance structure for $(\boldsymbol{\beta}_G, \boldsymbol{\beta}_I, \boldsymbol{\gamma}_G, \boldsymbol{\theta}_G)$ are

$$\mathrm{Var}\begin{pmatrix}\boldsymbol{\beta}_G\\ \boldsymbol{\beta}_I\\ \boldsymbol{\gamma}_G\\ \boldsymbol{\theta}_G\end{pmatrix} = \frac{1}{M}\begin{pmatrix}\sigma^2_{GI}\boldsymbol{I}_M & \rho_{GI}\boldsymbol{I}_M & \rho_{G,G2}\boldsymbol{I}_M & \rho_{GE}\boldsymbol{I}_M\\ \rho_{GI}\boldsymbol{I}_M & \sigma^2_I\boldsymbol{I}_M & \rho_{I,G2}\boldsymbol{I}_M & \rho_{IE}\boldsymbol{I}_M\\ \rho_{G,G2}\boldsymbol{I}_M & \rho_{I,G2}\boldsymbol{I}_M & \sigma^2_{G2}\boldsymbol{I}_M & \rho_{G2,E}\boldsymbol{I}_M\\ \rho_{GE}\boldsymbol{I}_M & \rho_{IE}\boldsymbol{I}_M & \rho_{G2,E}\boldsymbol{I}_M & \sigma^2_E\boldsymbol{I}_M\end{pmatrix},$$

where $\rho_{G2,E}$ is the genetic covariance between the second trait and environment, and $\rho_{IE}$ is the covariance between the genetic additive effects for the environment and GxE effects for the first trait.

#### 8.2 Estimating covariant GxE and oracle PGSxE under rGE

**Proposition 12** (Bivariate PIGEON-LDSC with rGE)**.** *Under the PIGEON-rGE model* (8)*, the slope in PIGEON-LDSC for estimating covariant GxE and oracle PGSxE is*

$$\frac{\sqrt{N_I N_G}(\rho_{G2,I} + \rho_{G2,E}\rho_{IE})}{CM}$$

$C = \sqrt{1 - \frac{\mathrm{z}^2_\mathrm{E}}{\mathrm{z}^2_\mathrm{E} + \mathrm{N}_G - 2}}$ *is a correction factor to correct for the influence of the environment main effects on the Z-score approximation in the derivation.*

*Proof.* See Proof of Proposition 11 in Appendix. □

This proposition has two implications:

1. The bias can be denoted as

$$\rho_{G2,I} + \rho_{G2,E}\rho_{IE} = r_{G2,I}\sqrt{\sigma_{G2}^2\sigma_I^2} + r_{G2,E}\sqrt{\sigma_{G2}^2\sigma_E^2}r_{IE}\sqrt{\sigma_I^2\sigma_E^2}$$
$$= \begin{cases} r_{GE}\sqrt{\sigma_{G2}^2\sigma_E^2}r_{IE}\sqrt{\sigma_I^2\sigma_E^2}, & r_{G2,I} = 0 \\ \rho_{G2,I}\left(1 + \frac{r_{G2,E}r_{IE}\sigma_E^2}{r_{G2,I}}\right), & r_{G2,I} \neq 0. \end{cases}$$

   Therefore, the non-zero covariance between genetic additive effects for the environment and GxE for the trait $\rho_{IE}$ and genetic covariance $\rho_{G2,E}$ between trait 2 and environment is necessary for the "naive" version PIGEON-LDSC (derived under G-E independence) to have biased and false positive ($r_{G2,I} = 0$ while the slope $\neq 0$) results.

2. We can obtain the covariant GxE $\rho_{GI}$ by subtracting $\rho_{G2,E}\rho_{IE}$ from the slope from of "naive" PIGEON-LDSC. The $\rho_{G2,E}$ and $\rho_{IE}$ can also be simply estimated from LDSC and PIGEON-LDSC using the GWAS summary statistics for the environment. We denote this new estimator for covariant GxE as the results for PIGEON-LDSC and implement it in our software.

##### 8.3 Impact on estimating GxE variance components

**Proposition 13** (PIGEON-LDSC for GxE variance components with rGE)**.** *Under the PIGEON-rGE model (8), the slope in PIGEON-LDSC for estimating GxE variance components with rGE is approximately* $\frac{(\sigma_I^2 + 2\rho_{IE}^2)N_I}{C^2 M}$*, where $C$ is the correction factor.*

*Proof.* See Proof of Proposition 12 in Appendix. □

Here, since $\sigma_I^2 + 2\rho_{IE}^2 = \sigma_I^2(1 + 2r_{EI}^2\sigma_E^2)$, we have the that "naive" PIGEON-LDSC do not have false positives results when testing the GxE variance components. Besides, we also propose a debiased estimator using the slope in PIGEON-LDSC minus by subtracting the $2\rho_{IE}^2$ from the PIGEON-LDSC slope. $\rho_{IE}$ can be estimated from PIGEON-LDSC using GWAS summary statistics for the environment.

#### 9 PGS portability and its impact on GxE inference

Next, we investigate the impact of PGS's "portability" problem in polygenic GxE inference. The motivation for this problem comes from a possible interpretation of our anorexia PGSxS for BMI: the original GWAS for Anorexia was ascertained toward women due to the higher prevalence of anorexia in women, although we argue later it is unlikely to bias our results. Also, we note that this is an issue generally neglected in all PGSxE analyses rather than a new problem induced by PIGEON. We are not aware of any solution under empirical

PGSxE design. Taking advantage of the PIGEON framework, we proposed several remedies to correct potential biases caused by these issues.

#### 9.1 Model and single-trait results

Recall the two cohort PIGEON model (2),

$$\begin{aligned}\boldsymbol{Y}^{(I)} &= \boldsymbol{G}^{(I)}\boldsymbol{\beta}_G + \boldsymbol{E}^{(I)}\boldsymbol{\beta}_{\boldsymbol{E}} + \boldsymbol{E}^{(I)}\boldsymbol{G}^{(I)}\boldsymbol{\beta}_I + \boldsymbol{\epsilon}_0^{(I)} + \boldsymbol{E}^{(I)}\boldsymbol{\epsilon}_1^{(I)} \\ \boldsymbol{Y}^{(G)} &= \boldsymbol{G}^{(G)}\boldsymbol{\beta}_G + \boldsymbol{E}^{(G)}\boldsymbol{\beta}_{\boldsymbol{E}} + \boldsymbol{E}^{(G)}\boldsymbol{G}^{(G)}\boldsymbol{\beta}_I + \boldsymbol{\epsilon}_0^{(G)} + \boldsymbol{E}\boldsymbol{\epsilon}_1^{(G)},\end{aligned} \tag{9}$$

where the superscript $(I)$ and $(G)$ denotes the PIGEON model (1) in GWIS cohort with $N_I$ individuals and GWAS cohort with $N_G$ individuals.

Suppose the raw-scale environment in GWIS and GWAS cohort are Bernoulli random variables with the probability of $P_{pop}$ for being 1 and $1 - P_{pop}$ for being 0. After standardization, we have the standardized environment in GWAS cohort is

$$\mathbb{P}(E_i^{(G)} = \sqrt{\frac{1-P_{pop}}{P_{pop}}}) = P_{pop} \text{ and } \mathbb{P}(E_i^{(G)} = -\sqrt{\frac{P_{pop}}{1-P_{pop}}}) = 1 - P_{pop}$$

We define the ascertained studies under GxE context as the GWAS samples with $P_{sample} \times N_G^{(sample)}$ individuals with raw-scale environment = 1 and $(1 - P_{sample}) \times N_G^{(sample)}$ individuals with raw-scale environment = 0, where $N_G^{(sample)}$ is the sample size. We have the conditional expectation for the phenotype in this ascertained study as

$$\begin{aligned}\mathbb{E}[Y_i|\text{ascertained studies}, \boldsymbol{\beta}_G, \boldsymbol{\beta}_E, \boldsymbol{\beta}_I] &= P_{sample}(\sum_{j=1}^{M} G_{ij}\beta_{G_j} + \sqrt{\frac{1-P_{pop}}{P_{pop}}}\beta_E + \sum_{j=1}^{M} G_{ij}\sqrt{\frac{1-P_{pop}}{P_{pop}}}\beta_{I_j}) + \\ &\quad (1-P_{sample})(\sum_{j=1}^{M} G_{ij}\beta_{G_j} - \sqrt{\frac{P_{pop}}{1-P_{pop}}}\beta_E - \sum_{j=1}^{M} G_{ij}\sqrt{\frac{P_{pop}}{1-P_{pop}}}\beta_{I_j}) \\ &= \sum_{j=1}^{M} G_{ij}(\beta_{Gj} + \frac{P_{sample} - P_{pop}}{\sqrt{(1-P_{pop})P_{pop}}}\beta_I) + \frac{P_{sample} - P_{pop}}{\sqrt{(1-P_{pop})P_{pop}}}\beta_E\end{aligned}$$

Therefore, in this ascertained study, we expect the PGS to be

$$PGS_i^{(asc)} = \sum_{j=1}^{M} G_{ij}(\beta_{Gj} + \frac{P_{sample} - P_{pop}}{\sqrt{(1-P_{pop})P_{pop}}}\beta_I),$$

rather than $\sum_{j=1}^{M} G_{ij}\beta_{Gj}$ expected in the population. With this setup, we have the next proportion to show what PGSxE estimates using ascertained GWAS.

**Proposition 14.** *Denote the* $\theta = \frac{P_{sample}-P_{pop}}{\sqrt{(1-P_{pop})P_{pop}}}$, *if we use the ascertained PGS* $PGS_i^{(asc)} = \sum_{j=1}^{M} G_{ij}(\beta_{Gj} + \theta\beta_I)$ *in the PGSxE regression*

$$Y_i \sim \alpha_G^{(asc)} PGS_i^{(asc)} + \alpha_E^{(asc)} E_i + \alpha_I^{(asc)} PGS_i^{(asc)} E_i,$$

*we have*

$$\alpha_I^{(asc)} = \frac{\rho_{GI} + \theta\sigma_I^2}{\sigma_G^2 + \theta^2\sigma_I^2 + 2\theta\rho_{GI}},$$

*where $\sigma_G^2$, $\sigma_I^2$, and $\rho_{GI}$ are the heritability, GxE variance components, and covariant GxE.*

*Proof.* See Proof of Proposition 13 in Appendix. □

Therefore, the hypothesis testing between oracle PGSxE and covariant GxE is not equivalent if using ascertained studies ($P_{sample} \neq P_{pop}$) as a GWAS sample.

#### 9.2 Multi-trait results

With this proposition, we can also have the results for oracle PGSxE (2) using PGS for other traits under the multi-trait model (4)

**Corollary 4.** *Denote the $\theta = \frac{P_{sample} - P_{pop}}{\sqrt{(1-P_{pop})P_{pop}}}$, if we use the ascertained PGS for $Y_{2i}$*

$$PGS_{2i}^{(asc)} = \sum_{j=1}^{M} G_{ij}(\gamma_{Gj} + \frac{P_{sample} - P_{pop}}{\sqrt{(1-P_{pop})P_{pop}}}\gamma_I)$$

*in the PGSxE regression*

$$Y_i \sim \alpha_G^{(PGS2,asc)} PGS_{2i}^{(asc)} + \alpha_E^{(PGS2,asc)} E_i + \alpha_I^{(PGS2,asc)} PGS_{2i}^{(asc)} E_i,$$

*we have*

$$\alpha_I^{(PGS2,asc)} = \frac{\rho_{G2,I} + \theta\rho_{I,I2}}{\sigma_{G2}^2 + \theta^2\sigma_{I2}^2 + 2\theta\rho_{G2,I2}},$$

*where $\sigma_{G2}^2$, $\sigma_{I2}^2$, and $\rho_{G2,I}$ is the heritability, GxE variance components, and covariant GxE for $Y_{2i}$, and $\rho_{I,I2}$ is the covariance between GxE effects for trait1 $Y_{1i}$ and trait2 $Y_{2i}$.*

*Proof.* This can be shown by extending Proof of Proposition 13 in the Appendix into two traits cases. □

Therefore, the hypothesis testing between multi-trait oracle PGSxE and covariant GxE is not equivalent if using ascertained studies ($P_{sample} \neq P_{pop}$) as GWAS sample.

In our findings for "Anorexia PGSxSex for BMI", the GWAS paper on anorexia did not observe differences in the polygenic structure of anorexia between males and females (*i.e.,* $\sigma_{I2}^2 = 0$). Therefore, the $\rho_{I,I2} = 0$ and it is unlikely that the "portability" issue leads to our interaction finding.

#### 9.3 Remedies

Here, we discuss three possible remedies:

1. **Ignore the bias.** The numerator for single-trait or multi-trait oracle PGSxE is

$$\rho_{GI} + \theta\sigma_I^2 = \rho_{GI}(1 + \theta\frac{\sqrt{\sigma_I^2}}{r_{GI}\sqrt{\sigma_G^2}}).$$

$$\rho_{G2,I} + \theta\rho_{I,I2} = \rho_{G2,I}(1 + \theta\frac{r_{I,I2}\sqrt{\sigma_I^2}}{r_{G,I2}\sqrt{\sigma_{G2}^2}}).$$

Note that the bias may be small in practice since the GxE variance components times the difference between a sample and population prevalence is much smaller than the heritability, i.e. $\theta\sqrt{\sigma_I^2} << r_{GI}\sqrt{\sigma_G^2}$ and $\theta_{I,I2}\sqrt{\sigma_I^2} << r_{G,I2}\sqrt{\sigma_{G2}^2}$. The risk for this approach is the violation of the statement above. Therefore, we consider the second remedy to correct for the bias

2. **Correct for the bias using GWIS summary statistics.** In the single trait case, we can directly correct for the bias since we have the GWIS summary statistics to estimate $\sigma_I^2$. In the multi-trait case, we can use the GWIS summary statistics for the second trait with the same environment to estimate $r_{I,I2}$ and correct for the bias. However, the GWIS summary statistics in the multi-trait case may not be easy to obtain (e.g. GxSex summary statistics for Anorexia).

3. **Conduct the GWAS in the GWIS cohort.** The final remedy directly tries to eliminate the bias by forcing the $\theta = 0$. To do so, the simplest way is to conduct the GWAS on the same cohort with the GWIS sample, and apply PIGEON to these GWIS and GWAS summary statistics. We note that this cannot be done using empirical PGSxE design due to the overfitted PGS. Since PIGEON estimates are robust to GWAS and GWIS sample overlap, this can be implemented. The limitation is a potential lack of power due to the limited sample size in the GWIS sample compared with the meta-analyzed GWAS summary statistics from the big consortium.

### 10 Some issues with input data

#### 10.1 Impact of covariates

Here, we add covariates in our PIGEON model (1):

$$\underbrace{\boldsymbol{Y}}_{\text{Trait}} = \underbrace{\boldsymbol{G\beta}_G}_{\text{Additive effects}} + \underbrace{\boldsymbol{E\beta_E}}_{\text{E's effects}} + \underbrace{\boldsymbol{EG\beta}_I}_{\text{GxE}} + \underbrace{\boldsymbol{\epsilon}_0}_{\text{Residual}} + \underbrace{\boldsymbol{E\epsilon}_1}_{\text{RxE}} + \underbrace{\boldsymbol{T\beta_T}}_{\text{covariates}}$$

where $\boldsymbol{T}$ is a $N_I \times K$ covariates matrix, $K$ is the number of covariates. For illustration, we assume the covariates (e.g. sex and age) are independent of the genetic components.

If we follow the original LDSC paper (Bulik-Sullivan et al., 2015a) to not consider the covariates' effect, we have $(\boldsymbol{EG})_j^T\boldsymbol{Y} = \sqrt{N_I}z_{Ij}$ in Proof of Proposition 5, where $z_{Ij}$ is the Z-score for SNPxE effects for j-th SNP. This relies on the assumption that the residual variance $\text{Var}(\epsilon_{0i}) \approx 1$, which is not the case in the presence of covariates and the environment's main effects.

Instead, we can use the formula below to adjust the covariates and environmental main effects:

$$(\boldsymbol{EG})_j^T \boldsymbol{Y} = \sqrt{N_I} z_{Ij} \times \sqrt{1 - R^2_{covar}},$$

where $R^2_{covar}$ is the R-squared obtained from the regression between the outcome traits and the covariates and environments. In our main texts, we only consider the environment's main effects, which leads to the correction factor

$$C = \sqrt{1 - R_E^2} = \sqrt{1 - \frac{Z_E^2}{Z_E^2 + N_I - 2}},$$

where $Z_E$ is the Z-score of environment effects.

Nevertheless, we note that the hypothesis testing is the same with or without this adjustment. The adjustment aims for the interpretation of the heritability and GxE variance components. Without adjustment, the heritability and GxE variance components is interpreted as the phenotypic residual variance (after adjusting for the covariates) explained by additive genetic and GxE effects (i.e. conditional variance). With the adjustment, the heritability and GxE variance components is interpreted as the raw phenotypic variance (before adjusting for the covariates) explained by additive genetic and GxE effects.

### 11 Additional simulations

#### 11.1 Heteroskedasticity variance in GxE variance components estimation

We used the simulation setting for GxE variance components described in main texts except changing the the residual variance $\sigma^2_{\epsilon_0} = 0.15 - \sigma^2_I$, heteroskedasticity variance $\sigma^2_{\epsilon_1} = 0.3$, and environmental main effect $\beta_E = \sqrt{0.05}$.

#### 11.2 Comparison with the measurement-error-corrected estimator in Becker et al.

We compared PIGEON with the measurement-error-corrected estimator in Becker et al in estimating PGSxE. We considered the simulation setting under G-E independence (described in Methods in the main texts) and used the code below:

```
python3 pgic.py \
--reg-data-file rep1.txt \
--outcome "Y" \
--pgi-var "PGS" \
--covariates "E" \
--pgi-interact-vars "E" \
--h2 0.5 \
--out rep1.txt
```

Here, instead of using the recommended GCTA (Yang et al., 2011) to estimate the h2, we directly plug in the true simulated h2 in the software to reduce the computational burden.

#### 11.3 GxE variance components simulation with rGE

We simulated the trait and environment for 20,000 samples in UKB by the PIGEON model with rGE (7):

$$\begin{aligned} \boldsymbol{Y} &= \boldsymbol{\mu} + \boldsymbol{G}\boldsymbol{\beta}_G + \boldsymbol{V}\boldsymbol{\beta}_{\boldsymbol{E}} + \boldsymbol{V}\boldsymbol{G}\boldsymbol{\beta}_I + \boldsymbol{\epsilon}_0 + \boldsymbol{V}\boldsymbol{\epsilon}_1 \\ \boldsymbol{V} &= \mathrm{diag}(\boldsymbol{v}) \\ \boldsymbol{v} &= \boldsymbol{G}\boldsymbol{\theta}_G + \sqrt{(1-\sigma_E^2)}\boldsymbol{e}, \end{aligned}$$

with the SNP effect size and residual simulated from a multivariate normal distribution:

$$\begin{pmatrix} \boldsymbol{\beta}_G \\ \boldsymbol{\beta}_I \\ \boldsymbol{\theta}_G \end{pmatrix} \sim \mathcal{N}\left[\begin{pmatrix} 0 \\ 0 \\ 0 \end{pmatrix}, \frac{1}{M}\begin{pmatrix} 0.5\boldsymbol{I}_M & \rho_{GI}\boldsymbol{I}_M & 0.07\boldsymbol{I}_M \\ \rho_{GI}\boldsymbol{I}_M & \sigma_I^2\boldsymbol{I}_M & \rho_{IE}\boldsymbol{I}_M \\ 0.07\boldsymbol{I}_M & \rho_{IE}^2\boldsymbol{I}_M & 0.25\boldsymbol{I}_M \end{pmatrix}\right], \begin{pmatrix} \epsilon_{i0} \\ \epsilon_{i1} \\ e_i \end{pmatrix} \sim \mathcal{N}\left[\begin{pmatrix} 0 \\ 0 \\ 0 \end{pmatrix}, \begin{pmatrix} \sigma_{\epsilon_0}^2 & 0 & 0 \\ 0 & 0.05 & 0 \\ 0 & 0 & 1 \end{pmatrix}\right]$$

where the GxE variance components $\sigma_I^2 \in \{0, 0.02, 0.04, 0.06, 0.08, 0.1\}$, $\rho_{IE} \in \{0, 0.8\sqrt{\sigma_I^2\sigma_E^2}\}$, $\rho_{GI} = 0.5\sqrt{\sigma_G^2\sigma_I^2}$, environment effect $\beta_v = \sqrt{0.1}$, the heteroskedasticity variance $\sigma_{\epsilon_1}^2 = 0.05$, $\sigma_E^2 = 0.25$, and the residual variance $\sigma_{\epsilon_0}^2$ is the value such that the phenotypic variance is 1. We then applied PIGEON-LDSC to GWIS summary statistics generated these 20,000 samples and evaluated the estimates for the GxE variance components component. We compare PIGEON with GxEsum and the naive estimation method without accounting for rGE.

#### 11.4 Covariant GxE & Oracle PGSxE simulations with rGE

We simulated the trait $\boldsymbol{Y}$ and environment for the first sub-cohort 20,000 samples and trait $\boldsymbol{Z}$ for the second sub-cohort 20,000 samples (without sample overlap with the first cohort) in UKB by the PIGEON model with rGE (8):

$$\begin{aligned} \boldsymbol{Y}_1 &= \boldsymbol{\mu} + \boldsymbol{G}\boldsymbol{\beta}_G + \boldsymbol{V}\boldsymbol{\beta}_{\boldsymbol{E}} + \boldsymbol{V}\boldsymbol{G}\boldsymbol{\beta}_I + \boldsymbol{\epsilon}_0 + \boldsymbol{V}\boldsymbol{\epsilon}_1 \\ \boldsymbol{V} &= \mathrm{diag}(\boldsymbol{v}) \\ \boldsymbol{v} &= \boldsymbol{G}\boldsymbol{\theta}_G + \sqrt{(1-\sigma_E^2)}\boldsymbol{e} \\ \boldsymbol{Y}_2 &= \boldsymbol{Z}\boldsymbol{\gamma}_G + \boldsymbol{\delta}, \end{aligned}$$

with the SNP effect size and residual simulated from a multivariate normal distribution:

$$\begin{pmatrix} \boldsymbol{\beta}_G \\ \boldsymbol{\beta}_I \\ \boldsymbol{\theta}_G \\ \boldsymbol{\gamma}_G \end{pmatrix} \sim \mathcal{N}\left[\begin{pmatrix} 0 \\ 0 \\ 0 \\ 0 \end{pmatrix}, \frac{1}{M}\begin{pmatrix} 0.5\boldsymbol{I}_M & 0.112\boldsymbol{I}_M & 0.07\boldsymbol{I}_M & 0.25\boldsymbol{I}_M \\ 0.112\boldsymbol{I}_M & 0.1\boldsymbol{I}_M & \rho_{IE}\boldsymbol{I}_M & \rho_{IZ}\boldsymbol{I}_M \\ 0.07\boldsymbol{I}_M & \rho_{IE}\boldsymbol{I}_M & 0.25\boldsymbol{I}_M & 0.2\boldsymbol{I}_M \\ 0.25\boldsymbol{I}_M & \rho_{IZ}\boldsymbol{I}_M & 0.2\boldsymbol{I}_M & 0.5\boldsymbol{I}_M \end{pmatrix}\right]$$

and

$$\begin{pmatrix} \epsilon_{i0} \\ \epsilon_{i1} \\ e_i \\ \delta_q \end{pmatrix} \sim \mathcal{N}\left[\begin{pmatrix} 0 \\ 0 \\ 0 \\ 0 \end{pmatrix}, \begin{pmatrix} \sigma^2_{\epsilon_0} & 0.05 & 0 & 0 \\ 0.05 & 0.05 & 0 & 0 \\ 0 & 0 & 1 & 0 \\ 0 & 0 & 0 & 0.5 \end{pmatrix}\right]$$

where the covariant GxE $\rho_{IZ} \in \{0, 0.01, 0.02, 0.03, 0.04, 0.05\}$, $\rho_{IE} \in \{0, 0.05\}$, $\sigma^2_E = 0.25$ too see if the bias and false positives are led by $\rho_{IE}$, and the residual variance $\sigma^2_{\epsilon_0}$ is the value such that the phenotypic variance for $\boldsymbol{Y}$ is 1. The $0.112 \approx 0.5\sqrt{0.5 \times 0.1}$ and $0.07 \approx 0.2\sqrt{0.5 \times 0.25}$ are generated using fixed correlation. We then applied PIGEON-LDSC to GWIS and GWAS summary statistics generating these first and second sub-cohort, respectively. We evaluated the estimates for PIGEON in estimating oracle PGSxE.

### 12 Appendix

#### 12.1 Proof of Proposition 1

*Proof.* Given (3), the heritability in environment $E_i = \sqrt{\frac{1-p}{p}}$ is

$$\begin{aligned} h^2|(E_i = \sqrt{\frac{1-p}{p}}) &= \frac{Var(\sum_{j=1}^{M} G_{ij}(\beta_{Gj} + \sqrt{\frac{1-p}{p}}\beta_{Ij}))}{Var(Y_{1i}|E_i = \sqrt{\frac{1-p}{p}})} \\ &= \frac{A_1}{A_1 + B_1} \end{aligned}$$

where $A_1 := \sigma^2_G + \frac{1-p}{p}\sigma^2_I + 2\sqrt{\frac{1-p}{p}}\rho_{GI}$, $B_1 := \sigma^2_{\epsilon 0} + \frac{1-p}{p}\sigma^2_{\epsilon 1} + 2\sqrt{\frac{1-p}{p}}\rho_{\epsilon_0,\epsilon_1}$

The heritability in environment $E_i = -\sqrt{\frac{p}{1-p}}$ is

$$\begin{aligned} h^2|(E_i = -\sqrt{\frac{p}{1-p}}) &= \frac{Var(\sum_{j=1}^{M} G_{ij}(\beta_{Gj} - \sqrt{\frac{p}{1-p}}\beta_{Ij}))}{Var(Y_{1i}|E_i = -\sqrt{\frac{p}{1-p}})} \\ &= \frac{A_0}{A_0 + B_0} \end{aligned}$$

where $A_0 := \sigma^2_G + \frac{p}{1-p}\sigma^2_I - 2\sqrt{\frac{p}{1-p}}\rho_{GI}$, $B_0 := \sigma^2_{\epsilon 0} + \frac{p}{1-p}\sigma^2_{\epsilon 1} - 2\sqrt{\frac{p}{1-p}}\rho_{\epsilon_0,\epsilon_1}$

Thus, the difference in heritability between the two environments is

$$h^2|(E_i = \sqrt{\frac{1-p}{p}}) - h^2|(E_i = -\sqrt{\frac{p}{1-p}})$$

$$= \frac{A_1B_0 - A_0B_1}{(A_1+B_1)(A_0+B_0)}$$

$$= \frac{\sigma_{\epsilon_0}^2[\frac{1-2p}{p(1-p)}\sigma_I^2 + \frac{2}{\sqrt{p(1-p)}}\rho_{GI}] + \sigma_{\epsilon_1}^2[\frac{2p-1}{p(1-p)}\sigma_G^2 + \frac{2}{\sqrt{p(1-p)}}\rho_{GI}] - \frac{2}{\sqrt{p(1-p)}}(\sigma_G^2+\sigma_I^2)\rho_{\epsilon_0,\epsilon_1}}{(A_1+B_1)(A_0+B_0)}$$

$$= \frac{\sigma_{\epsilon_0}^2\sqrt{\sigma_I^2}\left[(1-2p)\sqrt{\sigma_I^2} + 2r_{GI}\sqrt{p(1-p)\sigma_G^2}\right] + \sigma_{\epsilon_1}^2\sqrt{\sigma_G^2}\left[(2p-1)\sqrt{\sigma_G^2} + 2r_{GI}\sqrt{p(1-p)\sigma_I^2}\right] - 2\sqrt{p(1-p)}(\sigma_G^2+\sigma_I^2)\rho_{\epsilon_0,\epsilon_1}}{p(1-p)(A_1+B_1)(A_0+B_0)}$$

$$= \frac{\sqrt{\sigma_I^2\sigma_{\epsilon_0}^2}}{W}\left[(1-2p)\sqrt{\sigma_I^2\sigma_{\epsilon_0}^2} + 2\sqrt{p(1-p)}(r_{GI}\sqrt{\sigma_G^2\sigma_{\epsilon_0}^2} - r_{\epsilon_0,\epsilon_1}\sqrt{\sigma_I^2\sigma_{\epsilon_1}^2})\right] +$$
$$\frac{\sqrt{\sigma_G^2\sigma_{\epsilon_1}^2}}{W}\left[(2p-1)\sqrt{\sigma_G^2\sigma_{\epsilon_1}^2} + 2\sqrt{p(1-p)}(r_{GI}\sqrt{\sigma_I^2\sigma_{\epsilon_1}^2} - r_{\epsilon_0,\epsilon_1}\sqrt{\sigma_G^2\sigma_{\epsilon_0}^2})\right],$$

where $r_{GI} = \frac{\rho_{GI}}{\sqrt{\sigma_G^2\sigma_I^2}}$ and $W = p(1-p) \times \mathrm{Var}\left(Y_i \mid E_i = \sqrt{\frac{1-p}{p}}\right) \times \mathrm{Var}\left(Y_i \mid E_i = \sqrt{\frac{1-p}{p}}\right)$ is a scaling factor related to the phenotypic variance. □

#### 12.2 Proof of Proposition 2

*Proof.* Given (3) and Proof of Proposition 1, the difference of genetic variance between two environments is

$$A_1 - A_0 = \sqrt{\sigma_I^2}\left[(1-2p)\sqrt{\sigma_I^2} + 2r_{GI}\sqrt{p(1-p)\sigma_G^2}\right]$$

□

#### 12.3 Proof of Proposition 3

*Proof.* Given (3) and Proof of Proposition 1, the genetic correlation between traits across these two environments is

$$\frac{\mathrm{Cov}(\frac{\sum_{j=1}^M G_{ij}(\beta_{Gj}+\sqrt{\frac{1-p}{p}}\beta_{Ij})}{\sqrt{A_1+B_1}}, \frac{\sum_{j=1}^M G_{ij}(\beta_{Gj}-\sqrt{\frac{p}{1-p}}\beta_{Ij})}{\sqrt{A_0+B_0}})}{\sqrt{\frac{A_1A_0}{(A_1+B_1)(A_0+B_0)}}} = \frac{M\,\mathrm{Cov}(\beta_{Gj}+\sqrt{\frac{1-p}{p}}\beta_{Ij}, \beta_{Gj}-\sqrt{\frac{p}{1-p}}\beta_{Ij})}{\sqrt{A_1A_0}}$$

The genetic correlation $= 1$ between traits across these two environments implies

$$\frac{\mathrm{Cov}(\frac{\sum_{j=1}^M G_{ij}(\beta_{Gj}+\sqrt{\frac{1-p}{p}}\beta_{Ij})}{\sqrt{A_1+B_1}}, \frac{\sum_{j=1}^M G_{ij}(\beta_{Gj}-\sqrt{\frac{p}{1-p}}\beta_{Ij})}{\sqrt{A_0+B_0}})}{\sqrt{\frac{A_1A_0}{(A_1+B_1)(A_0+B_0)}}} = 1$$

$$\Leftrightarrow M\,\mathrm{Cov}(\beta_{Gj}+\sqrt{\frac{1-p}{p}}\beta_{Ij}, \beta_{Gj}-\sqrt{\frac{p}{1-p}}\beta_{Ij}) = \sqrt{A_1A_0}$$

$$\Leftrightarrow \sigma_G^2\sigma_I^2 = \rho_{GI}^2$$

$$\Leftrightarrow \sigma_I^2\sigma_G^2(r_{GI}^2 - 1) = 0$$

□

#### 12.4 Proof of Theorem 1

*Proof.* We denote the oracle PGS for individual $i$ as $PGS_i = \sum_j^M G_{ij}\beta_j$. We define

$$\alpha_I = \frac{\mathrm{Cov}\left(Y_i, PGS_i E_i\right)}{\mathrm{Var}\left(PGS_i E_i\right)}.$$

Since $PGS_i$ and $E_i$ are centered and uncorrelated, $\alpha_I$ is the oracle PGSxE interaction regression coefficient for ordinary linear regression $Y_i \sim \alpha_G PGS_i + \alpha_E E_i + \alpha_I PGS_i E_i$. Then we have

$$\begin{aligned}
\alpha_I &= \frac{\mathrm{Cov}(Y_{i1}, PGS_i E_i)}{\mathrm{Var}(PGS_i E_i)} \\
&= \frac{\mathrm{Cov}(\boldsymbol{G}_i \boldsymbol{\beta}_G + E_i \beta_E + \boldsymbol{G}_i * E_i \boldsymbol{\beta}_I + \epsilon_{0i} + \epsilon_{1i} E_i, \boldsymbol{G}_i \boldsymbol{\beta}_G E_i)}{\mathbb{E}[E_i]^2 \,\mathrm{Var}(PGS_i) + \mathbb{E}[PGS_i]^2 \,\mathrm{Var}(E_i) + \mathrm{Var}(PGS_i)\,\mathrm{Var}(E_i)} \\
&= \frac{\mathbb{E}[\boldsymbol{G}_i * E_i \boldsymbol{\beta}_I]\mathbb{E}[\boldsymbol{G}_i \boldsymbol{\beta}_G E_i]}{\sigma_G^2} \\
&= \frac{M\mathbb{E}[\boldsymbol{\beta}_I \boldsymbol{\beta}_G]}{\sigma_G^2} \\
&= \frac{\rho_{GI}}{\sigma_G^2}
\end{aligned}$$

□

#### 12.5 Proof of Lemma 1

*Proof.* Given the arbitrary location and scale transformation of the trait, oracle PGS and environment, we have

$$\begin{aligned}
& Y_i^{(arb)} = \mu^{(arb)} + \alpha_G^{(arb)} PGS_i^{(arb)} + \alpha_E^{(arb)} E_i^{(arb)} + \alpha_I^{(arb)} PGS_i^{(arb)} E_i^{(arb)} + q_i^{(arb)} \\
\Rightarrow\quad & \sqrt{\mathrm{Var}(Y_i^{(arb)})} Y_i + \mathbb{E}[Y_i^{(arb)}] = \\
& (\sqrt{\mathrm{Var}(PGS_i^{(arb)})} PGS_i + \mathbb{E}[PGS_i^{(arb)}]) \frac{\alpha_G^{(arb)}}{\sqrt{\sigma_G^2}} + \\
& (\sqrt{\mathrm{Var}(E_i^{(arb)})} E_i + \mathbb{E}[E_i^{(arb)}]) \alpha_E^{(arb)} + \\
& (\sqrt{\mathrm{Var}(PGS_i^{(arb)})} PGS_i + \mathbb{E}[PGS_i^{(arb)}])(\sqrt{\mathrm{Var}(E_i^{(arb)})} E_i + \mathbb{E}[E_i^{(arb)}]) \frac{\alpha_I^{(arb)}}{\sqrt{\sigma_G^2}} + q_i^{(arb)}
\end{aligned}$$

Matching the coefficient between the above equation and

$$Y_i = \alpha_G PGS_i + \alpha_E E_i + \alpha_I PGS_i E_i + q,$$

we have

$$\alpha_G = \frac{\sqrt{\mathrm{Var}(PGS_i^{(arb)})}(\alpha_G^{(arb)} + \alpha_I^{(arb)}\mathbb{E}[E_i^{(arb)}])}{\sqrt{\sigma_G^2}}$$

$$\alpha_I = \frac{\sqrt{\mathrm{Var}(PGS_i^{(arb)})\,\mathrm{Var}(E_i^{(arb)})}}{\sqrt{\mathrm{Var}(Y_i^{(arb)})\sigma_G^2}}\alpha_I^{(arb)}$$

Thus,

$$\alpha_G^{(arb)} = \frac{\alpha_G\sqrt{\sigma_G^2} - \alpha_I^{(arb)}\mathbb{E}[E_i^{(arb)}]}{\sqrt{\mathrm{Var}(PGS_i^{(arb)})}}$$

$$\alpha_I^{(arb)} = \frac{\sqrt{\mathrm{Var}(Y_i^{(arb)})\sigma_G^2}}{\sqrt{\mathrm{Var}(PGS_i^{(arb)})\,\mathrm{Var}(E_i^{(arb)})}}\alpha_I$$

□

#### 12.6 Proof of Theorem 2

*Proof.* The empirical PGSxE coefficient for regression $Y_i \sim \alpha_G^{(Emp)}\widehat{PGS}_i + \alpha_E^{(Emp)}E_i + \alpha_I^{(Emp)}\widehat{PGS}_iE_i$ is

$$\begin{aligned}
\alpha_I^{(Emp)} &= \frac{\mathrm{Cov}\left(Y_i, \widehat{PGS}_iE_i\right)}{\mathrm{Var}\left(\widehat{PGS}_iE_i\right)} \\
&= \frac{\mathrm{Cov}\left(y_{1i}, PGS_iE_i + s_iE_i\right)}{\sqrt{\mathrm{Var}\left(PGS_iE_i\right) + \mathrm{Var}\left(s_iE_i\right)}} \\
&= \frac{\mathrm{Cov}\left(y_{1i}, PGS_iE_i\right)}{\sqrt{\mathrm{Var}\left(PGS_iE_i\right) + \mathrm{Var}\left(s_iE_i\right)}} \\
&= \frac{\rho_{GI}}{\sqrt{\sigma_G^2 + \mathrm{Var}(s_i)}}
\end{aligned}$$

By theorem 1, the oracle PGSxE coefficient is

$$\alpha_I = \frac{\rho_{GI}}{\sigma_G^2}$$

Organizing the equation by plug in $\rho_{GI} = \alpha_I\sigma_G^2$, we have

$$\begin{aligned}
\alpha_I^{(Emp)} &= \frac{\alpha_I\sigma_G^2}{\sqrt{\sigma_G^2 + \mathrm{Var}(s_i)}} \\
&= \frac{\rho_{GI}}{\sigma_G^2} + \frac{(\sqrt{\sigma_G^2 + \mathrm{Var}(s_i)} - \sqrt{\sigma_G^2})}{\sqrt{(\sigma_G^2 + \mathrm{Var}(s_i))\sigma_G^2}}\rho_{GI} \\
&= \alpha_I + \frac{\sqrt{\sigma_G^2}(\sqrt{\sigma_G^2} - \sqrt{\sigma_G^2 + \mathrm{Var}(s_i)})}{\sqrt{(\sigma_G^2 + \mathrm{Var}(s_i))}}\alpha_I
\end{aligned}$$

□

##### 12.7 Proof of Theorem 3

*Proof.* Consider the GWAS with $N_G$ individuals,

$$\boldsymbol{Y}^{(G)} = \boldsymbol{G}^{(G)}\boldsymbol{\beta}_G + \boldsymbol{E}^{(G)}\beta_E + \boldsymbol{G}^{(G)} * \boldsymbol{E}^{(G)}\boldsymbol{\beta}_I + \boldsymbol{\epsilon}_0^{(G)} + \boldsymbol{\epsilon}_1^{(G)}\boldsymbol{E}^{(G)}$$

and the target sample with $N_I$ individuals,

$$\boldsymbol{Y}^{(I)} = \boldsymbol{G}^{(I)}\boldsymbol{\beta}_G + \boldsymbol{E}^{(I)}\beta_E + \boldsymbol{G}^{(I)} * \boldsymbol{E}^{(I)}\boldsymbol{\beta}_I + \boldsymbol{\epsilon}_0^{(I)} + \boldsymbol{\epsilon}_1^{(I)}\boldsymbol{E}^{(I)}$$

For this proof, we assume there are $N_S$ overlapped individuals in two samples. We further assumed that the columns of $\boldsymbol{G}^{(G)}$ and $\boldsymbol{G}^{(I)}$ are assumed to be independent after linkage disequilibrium (LD)-based pruning. We use simulations to verify the conclusion of Theorem 3 when the genotype matrix is not pruned.

Using the estimated additive effect for $j$-th SNP in discover GWAS $\hat{\beta}_{Gj} = \dfrac{\sum_i^{N_G} X_{ij}^{(G)} Y_i}{N_G}$ as SNP weights, the Emp, Ovpirical PGS for $q$-th individual is

then

$$\widehat{PGS}_q^{(I)} = \sum_j^M X_{qj}\hat{\beta}_{Gj}$$

The least squares estimators for Emp, Ovpirical PGSxE using the target sample is then

$$\begin{aligned}
\widehat{\alpha}_I^{(Emp,Ovp)} &= \frac{\sum\limits_q^{N_I} \widehat{PGS}_q^{(I)} E_q^{(I)} Y_q^{(I)}}{\sum\limits_q^{N_I} \widehat{PGS}_q^{(t)^2}} \\
&= \frac{\sum\limits_q^{N_I}\sum\limits_j^M X_{qj}^{(I)} \hat{\beta}_{Gj} E_q^{(I)} Y_q^{(I)}}{\sum\limits_q^{N_I} \widehat{PGS}_q^{(t)^2}} \\
&= \frac{\sum\limits_q^{N_I}\sum\limits_j^M X_{qj}^{(I)} \sum\limits_i^{N_G} X_{ij}^{(G)} Y_i^{(G)} E_q^{(I)} Y_q^{(I)}}{\sum\limits_q^{N_I} \widehat{PGS}_q^{(t)^2} N_G} \\
&= \frac{\sum\limits_q^{N_I}\sum\limits_i^{N_G}\sum\limits_j^M X_{qj}^{(I)} E_q^{(I)} Y_q^{(I)} X_{ij}^{(G)} Y_i^{(G)}}{\sum\limits_q^{N_I} \widehat{PGS}_q^{(t)^2} N_G}
\end{aligned}$$

Since $\widehat{\alpha}_I^{(Emp,Ovp)}$ is the least squares estimators, we have $\mathbb{E}[\widehat{\alpha}_I^{(Emp,Ovp)}] = \alpha_I^{(Emp,Ovp)}$.

Denote $\mathcal{O}_s$ as the set of samples overlapping discovery GWAS and target sample. We use $i \in \mathcal{O}_s$ and $i \notin \mathcal{O}_s$" to indicate that individual $i$ belongs or does not belong to $\mathcal{O}_s$, respectively. Then we have

$$\mathbb{E}\left[\sum_q^{N_I}\sum_i^{N_G}\sum_j^{M} X_{qj}^{(I)}E_q^{(I)}Y_q^{(I)}X_{ij}^{(G)}Y_i^{(G)}\right] = \sum_j^{M}\mathbb{E}\left[\sum_{(q,i)\in\mathcal{O}_s} X_{qj}^{(I)}E_q^{(I)}Y_q^{(I)}X_{ij}^{(G)}Y_i^{(G)}\right] + \sum_j^{M}\mathbb{E}\left[\sum_{(q,i)\notin\mathcal{O}_s} X_{qj}^{(I)}E_q^{(I)}Y_q^{(I)}X_{ij}^{(G)}Y_i^{(G)}\right]$$
$$= M[\frac{\rho_{GI}}{M}(N_GN_I + 2N_S) + \left(2\rho_{\boldsymbol{\epsilon_0},\boldsymbol{\epsilon_1}} + \beta_E^2\mu_E(3)\right)N_S]$$

An expectation of the denominator is

$$\mathbb{E}\left[\sum_q^{N_I}\widehat{PGS}_q^{(t)^2}N_G\right] = N_IN_G(\mathrm{Var}(s_q) + \sigma_G^2),$$

where $\mathrm{Var}(s_q)$ is the estimation error of Empirical PGS defined in.

Then, by the delta theorem, we have the following.

$$\begin{aligned}
\mathbb{E}\left[\widehat{\alpha}_I^{(Emp,Ovp)}\right] &\approx \frac{\mathbb{E}\left[\sum_q^{N_I}\sum_i^{N_G}\sum_j^{M} X_{qj}^{(I)}E_q^{(I)}Y_q^{(I)}X_{ij}^{(G)}Y_i^{(G)}\right]}{\mathbb{E}\left[\sum_q^{N_I}\widehat{PGS}_q^{(t)^2}N_G\right]} \\
&= \frac{[\frac{\rho_{GI}}{M}[N_GN_I + 2N_S] + [2\rho_{\boldsymbol{\epsilon_0},\boldsymbol{\epsilon_1}} + \beta_E^2\mu_E(3)]N_S]M}{N_IN_G(\mathrm{Var}(s_q) + \sigma_G^2)} \\
&\approx \frac{MN_S(2\rho_{GI} + 2\rho_{\boldsymbol{\epsilon_0},\boldsymbol{\epsilon_1}} + \beta_E^2\mu_E(3)) + N_GN_I\rho_{GI}}{N_IN_G(\mathrm{Var}(s_q) + \sigma_G^2)} \\
&= \frac{\rho_{GI}}{\mathrm{Var}(s_q) + \sigma_G^2} + \frac{MN_S(2\rho_{GI} + 2\rho_{\boldsymbol{\epsilon_0},\boldsymbol{\epsilon_1}} + \beta_E^2\mu_E(3))}{N_IN_G(\mathrm{Var}(s_q) + \sigma_G^2)}
\end{aligned}$$

Thus,

$$\begin{aligned}
\alpha_I^{(Emp,Ovp)} &= \mathbb{E}\left[\widehat{\alpha}_I^{(Emp,Ovp)}\right] \\
&= \frac{\rho_{GI}}{\mathrm{Var}(s_q) + \sigma_G^2} + \frac{MN_S(2\rho_{GI} + 2\rho_{\epsilon_0,\epsilon_1} + \beta_E^2\mu_E(3))}{N_IN_G(\mathrm{Var}(s_q) + \sigma_G^2)} \\
&= \alpha_I\left[1 + \frac{\sqrt{\sigma_G^2}(\sqrt{\sigma_G^2} - \sqrt{\sigma_G^2 + \frac{M}{N_G}})}{\sqrt{(\sigma_G^2 + \frac{M}{N_G})}}\right](1 + \frac{2MN_S}{N_IN_G}) + \frac{MN_S(2\rho_{\boldsymbol{\epsilon_0},\boldsymbol{\epsilon_1}} + \beta_E^2\mu_E(3))}{N_IN_G(\frac{M}{N_G} + \sigma_G^2)}
\end{aligned}$$

□

#### 12.8 Proof of Proposition 4

*Proof.* Consider the polygenic GxE model in :

$$\boldsymbol{Y} = \boldsymbol{G}\boldsymbol{\beta}_G + \boldsymbol{E}\boldsymbol{\beta}_{\boldsymbol{E}} + \boldsymbol{E}\boldsymbol{G}\boldsymbol{\beta}_I + \boldsymbol{\epsilon}_0 + \boldsymbol{E}\boldsymbol{\epsilon}_1$$

We have the Z-score for SNPxE interaction have the moment condition:

$$\mathbb{E}[\boldsymbol{z}_I|\boldsymbol{\beta}_G, \boldsymbol{\beta}_{\boldsymbol{E}}, \boldsymbol{\beta}_I] = \frac{\boldsymbol{G}^T\boldsymbol{E}\boldsymbol{G}\boldsymbol{\beta}_G}{\sqrt{N_I}} + \frac{\boldsymbol{G}^T\boldsymbol{E}\boldsymbol{E}\boldsymbol{\beta}_E}{\sqrt{N_I}} + \frac{\boldsymbol{G}^T\boldsymbol{E}\boldsymbol{E}\boldsymbol{G}\boldsymbol{\beta}_I}{\sqrt{N_I}}$$

and

$$\text{Var}[\boldsymbol{z}_I|\boldsymbol{\beta}_G, \boldsymbol{\beta}_{\boldsymbol{E}}, \boldsymbol{\beta}_I] = \frac{\sigma^2_{\epsilon_0}\boldsymbol{G}^T\boldsymbol{E}\boldsymbol{E}\boldsymbol{G}}{N_I} + \frac{\sigma^2_{\epsilon_1}\boldsymbol{G}^T\boldsymbol{E}\boldsymbol{E}\boldsymbol{E}\boldsymbol{E}\boldsymbol{G}}{N_I}$$

Therefore, by the law of total expectation, we have the expectation of the squared SNPxE Z-score for j-th SNPs is

$$\begin{aligned}
\mathbb{E}[z^2_{Ij}] &= \mathbb{E}[\mathbb{E}[z^2_{Ij}|\boldsymbol{\beta}_G, \boldsymbol{\beta}_{\boldsymbol{E}}, \boldsymbol{\beta}_I]] = \mathbb{E}[\mathbb{E}[z_{Ij}|\boldsymbol{\beta}_G, \boldsymbol{\beta}_{\boldsymbol{E}}, \boldsymbol{\beta}_I]^2 + \text{Var}[z_{Ij}|\boldsymbol{\beta}_G, \boldsymbol{\beta}_{\boldsymbol{E}}, \boldsymbol{\beta}_I]] \\
&= \mathbb{E}[(\frac{\boldsymbol{G}^T_{j\cdot}\boldsymbol{E}\boldsymbol{G}\boldsymbol{\beta}_G}{\sqrt{N_I}} + \frac{\boldsymbol{G}^T_{j\cdot}\boldsymbol{E}\boldsymbol{E}\boldsymbol{\beta}_E}{\sqrt{N_I}} + \frac{\boldsymbol{G}^T_{j\cdot}\boldsymbol{E}\boldsymbol{E}\boldsymbol{G}\boldsymbol{\beta}_I}{\sqrt{N_I}})^2 + \frac{\sigma^2_{\epsilon_0}\boldsymbol{G}^T_{j\cdot}\boldsymbol{E}\boldsymbol{E}\boldsymbol{G}_{j\cdot}}{N_I} + \frac{\sigma^2_{\epsilon_1}\boldsymbol{G}^T_{j\cdot}\boldsymbol{E}\boldsymbol{E}\boldsymbol{E}\boldsymbol{E}\boldsymbol{G}_{j\cdot}}{N_I}] \\
&= \mathbb{E}[(\frac{\boldsymbol{G}^T_{j\cdot}\boldsymbol{E}\boldsymbol{G}\boldsymbol{\beta}_G}{\sqrt{N_I}} + \frac{\boldsymbol{G}^T_{j\cdot}\boldsymbol{E}\boldsymbol{E}\boldsymbol{\beta}_E}{\sqrt{N_I}} + \frac{\boldsymbol{G}^T_{j\cdot}\boldsymbol{E}\boldsymbol{E}\boldsymbol{G}\boldsymbol{\beta}_I}{\sqrt{N_I}})^2 + \frac{\sigma^2_{\epsilon_0}\boldsymbol{G}^T_{j\cdot}\boldsymbol{E}\boldsymbol{E}\boldsymbol{G}_{j\cdot}}{N_I} + \frac{\sigma^2_{\epsilon_1}\boldsymbol{G}^T_{j\cdot}\boldsymbol{E}\boldsymbol{E}\boldsymbol{E}\boldsymbol{E}\boldsymbol{G}_{j\cdot}}{N_I}] \\
&= \mathbb{E}[\frac{\boldsymbol{G}^T_{j\cdot}\boldsymbol{E}\boldsymbol{G}\boldsymbol{\beta}_G(\boldsymbol{G}^T_{j\cdot}\boldsymbol{E}\boldsymbol{G}\boldsymbol{\beta}_G)^T}{N_I} + \frac{\boldsymbol{G}^T_{j\cdot}\boldsymbol{E}\boldsymbol{E}\boldsymbol{\beta}_E(\boldsymbol{G}^T_{j\cdot}\boldsymbol{E}\boldsymbol{E}\boldsymbol{\beta}_E)^T}{N_I} + \frac{\boldsymbol{G}^T_{j\cdot}\boldsymbol{E}\boldsymbol{E}\boldsymbol{G}\boldsymbol{\beta}_I(\boldsymbol{G}^T_{j\cdot}\boldsymbol{E}\boldsymbol{E}\boldsymbol{G}\boldsymbol{\beta}_I)^T}{N_I} + \\
&\quad \frac{\sigma^2_{\epsilon_0}\boldsymbol{G}^T_{j\cdot}\boldsymbol{E}\boldsymbol{E}\boldsymbol{G}_{j\cdot}}{N_I} + \frac{\sigma^2_{\epsilon_1}\boldsymbol{G}^T_{j\cdot}\boldsymbol{E}\boldsymbol{E}\boldsymbol{E}\boldsymbol{E}\boldsymbol{G}_{j\cdot}}{N_I}] \\
&= \frac{\sigma^2_g M}{M} + \beta^2_E\mu_E(4) + \frac{\sigma^2_I}{M}[\frac{(N^2_I - N_I)}{N_I}\sum^M_k r^2_{jk} + \frac{N_I(\sum^M_k 2r^2_{jk} + M)\mu_E(4)}{N_I}] + \sigma^2_{\epsilon_0} + \mu_E(4)\sigma^2_{\epsilon_1} \\
&= \frac{\sigma^2_I N_I}{M}\ell_j + 1 + (\beta^2_E + \sigma^2_{\epsilon_1} + \sigma^2_I)(\mu_E(4) - 1)
\end{aligned}$$

where $\ell_j = \sum^M_k r^2_{jk}$ is the LD score for j-th SNP □

#### 12.9 Proof of Proposition 5

*Proof.* Denote subscript "$(S)$" the vector/matrix for the overlapped samples. Then we can have

$$\begin{aligned}
&\mathbb{E}[z_{Ij}z_{Gj}] \\
&= \frac{1}{C\sqrt{N_G N_I}}\mathbb{E}\left[(\boldsymbol{G}^{(I)} * \boldsymbol{E}^{(I)})^\top_j \boldsymbol{Y}^{(I)}\boldsymbol{Y}^{(G)^\top}\boldsymbol{G}^{(G)}_j\right] \\
&= \frac{\rho_{GI}}{CM\sqrt{N_G N_I}}\mathbb{E}\left[(\boldsymbol{G}^{(I)} * \boldsymbol{E}^{(I)})^\top_j(\boldsymbol{G}^{(I)} * \boldsymbol{E}^{(I)})\mathbf{I}_M\boldsymbol{G}^{(G)^\top}\boldsymbol{G}^{(G)}_j\right] + \frac{\rho_{\boldsymbol{\epsilon_0},\boldsymbol{\epsilon_1}}}{C\sqrt{N_G N_I}}\mathbb{E}\left[(\boldsymbol{G}^{(S)} * \boldsymbol{E}^{(S)})^\top_j\mathbf{1}_{N_S}\boldsymbol{E}^{(S)^\top}\boldsymbol{G}^{(S)}_j\right] \\
&\quad + \frac{\rho_{\boldsymbol{\epsilon_0},\boldsymbol{\epsilon_1}}}{C\sqrt{N_G N_I}}\mathbb{E}\left[(\boldsymbol{G}^{(S)} * \boldsymbol{E}^{(S)})^\top_j\boldsymbol{E}^{(S)}\mathbf{1}^\top_{N_S}\boldsymbol{G}^{(S)}_j\right] + \frac{\beta^2_E}{C\sqrt{N_G N_I}}\mathbb{E}\left[(\boldsymbol{G}^{(S)} * \boldsymbol{E}^{(S)})^\top_j\boldsymbol{E}^{(S)^\top}\boldsymbol{E}^{(S)^\top}\boldsymbol{G}^{(S)}_j\right] \\
&\quad + \frac{\rho_{GI}}{CM\sqrt{N_G N_I}}\mathbb{E}\left[(\boldsymbol{G}^{(S)} * \boldsymbol{E}^{(S)})^\top_j\boldsymbol{G}^{(S)}\mathbf{I}_M(\boldsymbol{G}^{(I)} * \boldsymbol{E}^{(I)})\boldsymbol{G}^{(S)}_j\right]
\end{aligned}$$

We further have

$$\mathbb{E}\left[(\boldsymbol{G}^{(I)} * \boldsymbol{E}^{(I)})^\top_j(\boldsymbol{G}^{(I)} * \boldsymbol{E}^{(I)})\mathbf{I}_M\boldsymbol{G}^{(G)^\top}\boldsymbol{G}^{(G)}_j\right] = \sum^M_{k=1}\sum^{N_G}_{i=1}\sum^{N_I}_{q=1}\mathbb{E}\left[E^{(I)^2}_i\right]\mathbb{E}\left[G^{(I)}_{qk}G^{(I)}_{qj}G^{(G)}_{ik}G^{(G)}_{ij}\right],$$

Following the MQS paper (Zhou, 2017), by Isserlis' theorem, we have

$$\mathbb{E}\left[G^{(I)}_{qk}G^{(I)}_{qj}G^{(G)}_{ik}G^{(G)}_{ij}\right] = \mathbb{E}\left[G^{(I)}_{qk}G^{(I)}_{qj}\right]\mathbb{E}\left[G^{(G)}_{ik}G^{(G)}_{ij}\right] + \mathbb{E}\left[G^{(I)}_{qk}G^{(G)}_{ik}\right]\mathbb{E}\left[G^{(I)}_{qj}G^{(G)}_{ij}\right] + \mathbb{E}\left[G^{(I)}_{qk}G^{(G)}_{ij}\right]\mathbb{E}\left[G^{(I)}_{qj}G^{(G)}_{ik}\right].$$

We further have

$$\mathbb{E}\left[G_{qk}^{(I)^2}\right] = \mathbb{E}\left[G_{ij}^{(G)^2}\right] = 1$$
$$\mathbb{E}\left[G_{ik}^{(G)}G_{ij}^{(G)}\right] = \mathbb{E}\left[G_{qk}^{(I)}G_{qj}^{(I)}\right] = r_{jk}$$
$$\mathbb{E}\left[G_{ik}^{(G)}G_{qk}^{(I)}\right] = \mathbb{E}\left[G_{ik}^{(G)}G_{qj}^{(I)}\right] = 0 \text{ for } i \neq q$$

$$\begin{aligned}\mathbb{E}\left[G_{ik}^{(G)^2}G_{ij}^{(G)^2}\right] &= \mathbb{E}\left[G_{ik}^{(G)}G_{ij}^{(G)}\right]^2 + \mathrm{Var}\left[G_{ik}^{(G)}G_{ij}^{(G)}\right] \\ &= \mathbb{E}\left[G_{ik}^{(G)}G_{ij}^{(G)}\right]^2 + \mathbb{E}\left[\mathrm{Var}\left[G_{ik}^{(G)}G_{ij}^{(G)} \mid G_{ik}^{(G)}\right]\right] + \mathrm{Var}\left[\mathbb{E}\left[G_{ik}^{(G)}G_{ij}^{(G)} \mid G_{ik}^{(G)}\right]\right] \\ &= r_{jk}^2 + 1 - r_{jk}^2 + 2r_{jk}^2 \\ &= 1 + 2r_{jk}^2\end{aligned}$$

Take these back, we have

$$\begin{aligned}\mathbb{E}\left[(\boldsymbol{G}^{(I)} * \boldsymbol{E}^{(I)})_j^\top (\boldsymbol{G}^{(I)} * \boldsymbol{E}^{(I)})\mathbf{I}_M \boldsymbol{G}^{(G)^\top}\boldsymbol{G}_j^{(G)}\right] &= \sum_{k=1}^{M}\sum_{i=1}^{N_G}\sum_{q=1}^{N_I}\mathbb{E}\left[G_{qk}^{(I)}G_{qj}^{(I)}G_{ik}^{(G)}G_{ij}^{(G)}\right] \\ &= \sum_{k=1}^{M}\left[N_S\left(1+2r_{jk}^2\right) + \left(N_S^2 - N_S\right)r_{jk}^2 + \left(N_I N_G - N_S^2\right)r_{jk}^2\right] \\ &\approx N_G N_I \sum_{k=1}^{M} r_{jk}^2 + N_S M \\ &= N_G N_I \ell_j + N_S M\end{aligned}$$

where the approximation comes from $\frac{N_S}{N_G N_I} \ll 1$. Further, we have

$$\mathbb{E}\left[(\boldsymbol{G}^{(S)} * \boldsymbol{E}^{(S)})_j^\top \mathbf{1}_{N_S}\boldsymbol{E}^{(S)^\top}\boldsymbol{G}_j^{(S)}\right] = N_S$$
$$\mathbb{E}\left[(\boldsymbol{G}^{(S)} * \boldsymbol{E}^{(S)})_j^\top \boldsymbol{E}^{(S)}\mathbf{1}_{N_S}^\top\boldsymbol{G}_j^{(S)}\right] = N_S$$
$$\mathbb{E}\left[(\boldsymbol{G}^{(S)} * \boldsymbol{E}^{(S)})_j^\top \boldsymbol{E}^{(S)^\top}\boldsymbol{E}^{(S)^\top}\boldsymbol{G}_j^{(S)}\right] = N_S\mu_E(3)$$
$$\mathbb{E}\left[(\boldsymbol{G}^{(S)} * \boldsymbol{E}^{(S)})_j^\top \boldsymbol{G}^{(S)}\mathbf{I}_M(\boldsymbol{G}^{(S)} * \boldsymbol{E}^{(S)})\boldsymbol{G}_j^{(S)}\right] = N_S\sum_{k=1}^{M}\left(1+2r_{jk}^2\right) = N_S(M+2\ell_j)$$

Taken together, we have

$$\begin{aligned}\mathbb{E}[z_{Ij}z_{Gj}] &= \frac{1}{C\sqrt{N_G N_I}}\left(N_G N_I \ell_j \frac{\rho_{GI}}{M} + N_s\rho_{GI} + \frac{\rho_{GI}}{M}N_S\left(M+2\ell_j\right) + N_S\rho_{\boldsymbol{\epsilon_0},\boldsymbol{\epsilon}_1} + N_S\rho_{\boldsymbol{\epsilon_0},\boldsymbol{\epsilon}_1} + N_S\beta_E^2\mu_E(3)\right) \\ &\approx \frac{\sqrt{N_I N_G}\rho_{GI}}{CM}\ell_j + \frac{N_S}{C\sqrt{N_I N_G}}\left(2\rho_{GI} + 2\rho_{\epsilon_0,\boldsymbol{\epsilon}_1} + \mu_E(3)\right),\end{aligned}$$

where the approximation comes from $\frac{N_S}{N_G N_I} \ll 1$. □

#### 12.10 Proof of Proposition 9

*Proof.* Consider the oracle Dichotomized PGS-by-E regression We have

$$\begin{aligned}
\alpha_I^{(dicho,q)} &= \frac{\text{Cov}(Y_{i1}, \mathbb{1}(PGS_i > w)E_i)}{\text{Var}(\mathbb{1}(PGS_i > w)E_i)} \\
&= \frac{\text{Cov}(\boldsymbol{G}_i\boldsymbol{\beta}_G + E_i\beta_E + \boldsymbol{G}_i\boldsymbol{\beta}_I E_i + \delta_{0i} + \delta_{1i}E_i, \mathbb{1}(\boldsymbol{G}_i\boldsymbol{\beta}_G > w)E_i)}{\mathbb{E}[(\mathbb{1}(PGS_i > w))^2]} \\
&= \frac{\mathbb{E}[E_i^2]\mathbb{E}[\mathbb{1}(\boldsymbol{G}_i\boldsymbol{\beta}_G > w)\boldsymbol{G}_i\boldsymbol{\beta}_I]}{\mathbb{E}[\mathbb{1}(PGS_i > w)]} \\
&= \mathbb{E}[\boldsymbol{G}_i\boldsymbol{\beta}_I | \boldsymbol{G}_i\boldsymbol{\beta}_G > w]
\end{aligned}$$

Here, $\boldsymbol{G}_i\boldsymbol{\beta}_I$ and $\boldsymbol{G}_i\boldsymbol{\beta}_G$ are bivariate normal distributed as:

$$\begin{pmatrix} \boldsymbol{G}_i\boldsymbol{\beta}_I \\ \boldsymbol{G}_i\boldsymbol{\beta}_G \end{pmatrix} \sim \mathcal{N}\left( \begin{pmatrix} 0 \\ 0 \end{pmatrix}, \begin{pmatrix} \sigma_I^2 & \rho_{G2,I} \\ \rho_{G2,I} & \sigma_{G2}^2 \end{pmatrix} \right).$$

Thus, according to the theory of conditional expectation for bivariate normal distribution. We have

$$\mathbb{E}[\boldsymbol{G}_i\boldsymbol{\beta}_I | \boldsymbol{G}_i\boldsymbol{\beta}_G > w] = \frac{r_{G2,I}\sigma_I \Phi\left(\frac{w}{\sigma_{G2}}\right)}{1 - \Phi\left(\frac{w}{\sigma_{G2}}\right)},$$

where $\Phi$ denotes the standard normal density function, and $\Phi$ is the standard normal cumulative distribution function.

Typically, people choose $w$ based on the top percentile of the PGS distribution. Suppose that one recodes the PGS into a 0/1 variable such that the individuals in the top $q$ are coded as 1 and the others are coded as 0. Then we have

$$w = \inf\{x \in \mathbb{R} : 1 - q \leq F_{\boldsymbol{G}_i\boldsymbol{\gamma}_G}(x)\} = \sigma_{G2}\sqrt{2}\,\text{erf}^{-1}(1 - 2q),$$

where $F$ is the cumulative distribution function for $\boldsymbol{G}_i\boldsymbol{\gamma}_G$, erf is the error function defined as $\text{erf}(x) = \frac{2}{\sqrt{\pi}}\int_0^x e^{-t^2}dt$.

Thus,

$$\mathbb{E}[\boldsymbol{G}_i\boldsymbol{\beta}_I | \boldsymbol{G}_i\boldsymbol{\gamma}_G > w] = \frac{r_{G2,I}\sigma_I \Phi\left(\sqrt{2}\,\text{erf}^{-1}(1-2q)\right)}{1 - \Phi\left(\sqrt{2}\,\text{erf}^{-1}(1-2q)\right)} = \frac{r_{G2,I}\sigma_I \Phi\left(\sqrt{2}\,\text{erf}^{-1}(1-2q)\right)}{q},$$

Taking together,

$$\begin{aligned}
\alpha_I^{(w)} &= \frac{r_{G2,I}\sqrt{\sigma_I^2}\Phi\left(\sqrt{2}\,\text{erf}^{-1}(1-2q)\right)}{q} \\
&= \frac{\rho_{G2,I}\Phi\left(\sqrt{2}\,\text{erf}^{-1}(1-2q)\right)}{q\sqrt{\sigma_G^2}}
\end{aligned}$$

□

#### 12.11 Proof of Proposition 10

*Proof.* The design matrix for this regression is

$$D = \left( E \quad PGS^{(std)} \quad PGS_2^{(std)} \quad EPGS^{(std)} \quad EPGS_2^{(std)} \right)$$

Due to the assumption of G-E independence, $D^T D$ is a block matrix, and the part related to the estimation of the interaction coefficient is

$$\begin{bmatrix} \alpha_I^{(PGS,cond)} \\ \alpha_I^{(PGS2,cond)} \end{bmatrix} = \begin{bmatrix} \text{Var}(PGS_i^{(std)} E_i) & \text{Cov}(PGS_{2i}^{(std)} E_i, PGS_{2i}^{(std)} E_i) \\ \text{Cov}(PGS_i^{(std)} E_i, PGS_{3i} E_i) & \text{Var}(PGS_{2i}^{(std)} E_i) \end{bmatrix}^{-1} \begin{bmatrix} \text{Cov}(Y_i, PGS_i^{(std)} E_i) \\ \text{Cov}(Y, PGS_{2i}^{(std)} E_i) \end{bmatrix}$$
$$= \begin{bmatrix} 1 & r_{G,G2} \\ r_{G,G2} & 1 \end{bmatrix}^{-1} \begin{bmatrix} \frac{\rho_{GI}}{\sqrt{\sigma_G^2}} \\ \frac{\rho_{G2,I}}{\sqrt{\sigma_{G2}^2}} \end{bmatrix}$$

□

#### 12.12 Proof of Proposition 11

*Proof.* The PIGEON-rGE model (8) is

$$\begin{aligned} Y_1 &= \mu + G\beta_G + V\beta_E + VG\beta_I + \epsilon_0 + V\epsilon_1 \\ V &= \text{diag}(v) \\ v &= G\theta_G + \sqrt{(1-\sigma_E^2)}e \\ Y_2 &= Z\gamma_G + \delta_0. \end{aligned} \tag{10}$$

We can rewrite $V$ as $V = \text{diag}(G\theta_G) + \sqrt{(1-\sigma_E^2)}\text{diag}(e)$.

We obtain the linear regression Z-score for interaction effects in GWIS for $Y_1$ by performing the linear regression $Y_1 \sim G_j + V + VG_j$. We note that centering outcomes or predictors in linear regression will not change the Z-score for interaction effects. By centering the outcome $Y_1$, the intercept term in the regression is negligible. Thus, we can directly obtain the Z-score without doing the inverse of the Gram matrix of the design matrix. Then, the Z-score for interaction effects for j-th SNP,

$$z_{I,j} \approx \frac{(VG)_j^\top (Y_1 - \mathbb{E}[Y_1])}{C\sqrt{N_I}},$$

where $C = \sqrt{1 - \frac{\mathrm{z}_\mathrm{E}^2}{\mathrm{z}_\mathrm{E}^2 + \mathrm{N}_G - 2}}$ is a correction factor.

Also, we have the linear regression Z-score of j-th SNP's additive genetic effects for $Y_2$ is $z_{G2,j} = \frac{Z_j^\top Y_2}{\sqrt{N_G}}$

Then we have

$$
\begin{aligned}
&\mathbb{E}\left[z_{I,j}z_{G2,j} \mid \boldsymbol{G},\boldsymbol{V},\boldsymbol{Z}\right] \\
&= \frac{1}{C\sqrt{N_I N_G}}\mathbb{E}\left[(\boldsymbol{V}\boldsymbol{G})_j^\top(\boldsymbol{Y}_1-\mathbb{E}[\boldsymbol{Y}_1])\boldsymbol{Y}_2^\top\boldsymbol{Z}_j\right] \\
&= \frac{1}{C\sqrt{N_I N_G}}\mathbb{E}\left[(\boldsymbol{V}\boldsymbol{G})_j^\top(\boldsymbol{G}\boldsymbol{\beta}_G+(\boldsymbol{G}\boldsymbol{\theta}_G+\sqrt{(1-\sigma_E^2)}\boldsymbol{e})\beta_E+\text{diag}(\boldsymbol{G}\boldsymbol{\theta}_G+\sqrt{(1-\sigma_E^2)}\boldsymbol{e})\boldsymbol{G}\boldsymbol{\beta}_I+\boldsymbol{\epsilon}_0+\boldsymbol{V}\boldsymbol{\epsilon}_1)(\boldsymbol{Z}\boldsymbol{\gamma}_G+\boldsymbol{\delta}_0)^\top\boldsymbol{Z}_j\right] \\
&= \frac{1}{C\sqrt{N_I N_G}}(T_1+T_2-T_3)
\end{aligned}
$$

where

$$
\begin{aligned}
T_1 &= (\boldsymbol{V}\boldsymbol{G})_j^\top(\frac{\rho_{G2,I}}{M}\boldsymbol{V}\boldsymbol{G}\boldsymbol{I}_M\boldsymbol{Z}^\top)\boldsymbol{Z}_j, \\
T_2 &= (\text{diag}(\boldsymbol{G}\boldsymbol{\theta}_G)\boldsymbol{G})_j^\top(\text{diag}(\boldsymbol{G}\boldsymbol{\theta}_G)\boldsymbol{G}\boldsymbol{\beta}_I\boldsymbol{\gamma}_G^\top\boldsymbol{Z}^\top, \\
T_3 &= (\text{diag}(\boldsymbol{G}\boldsymbol{\theta}_G)\boldsymbol{G})_j^\top\mathbb{E}[\text{diag}(\boldsymbol{G}\boldsymbol{\theta}_G)\boldsymbol{G}\boldsymbol{\beta}_I]\boldsymbol{\gamma}_G^\top\boldsymbol{Z}^\top\boldsymbol{Z}_j
\end{aligned}
$$

Then, we remove the conditioning on $\boldsymbol{G},\boldsymbol{V}$ and $Z$.

$$
\mathbb{E}\left[z_{I,j}z_{G2,j}\right] = \mathbb{E}[\mathbb{E}\left[z_{I,j}z_{G2,j} \mid \boldsymbol{G},\boldsymbol{V},\boldsymbol{Z}\right]]
$$

For the first term, with the derivation under the G-E independence case, we have

$$
\mathbb{E}[T_1] = \mathbb{E}[(\boldsymbol{V}\boldsymbol{G})_j^\top(\frac{\rho_{G2,I}}{M}\boldsymbol{V}\boldsymbol{G}\boldsymbol{I}_M\boldsymbol{Z}^\top)\boldsymbol{Z}_j] = (1-\sigma_E^2)(N_I N_G \ell_j\frac{\rho_{G2,I}}{M}+N_s\rho_{G2,I})
$$

For the second term, we have

$$
\begin{aligned}
\mathbb{E}[T_2] &= \mathbb{E}[\text{diag}(\boldsymbol{G}\boldsymbol{\theta}_G)(\boldsymbol{G})_j^\top(\text{diag}(\boldsymbol{G}\boldsymbol{\theta}_G)\boldsymbol{G}\boldsymbol{\beta}_I\boldsymbol{\gamma}_G^\top\boldsymbol{Z}^\top)\boldsymbol{Z}_j] \\
&= \sum_{i=1}^{N_I}\sum_{q=1}^{N_G}\sum_{a=1}^{M}\sum_{b=1}^{M}\sum_{c=1}^{M}\sum_{d=1}^{M}\mathbb{E}[G_{ij}G_{ia}\theta_a G_{ib}\beta_{Ib}G_{ic}\theta_c Z_{qj}Z_{qd}\gamma_{Gd}] \\
&= \sum_{i=1}^{N_I}\sum_{q=1}^{N_G}\sum_{a=1}^{M}\sum_{b=1}^{M}\sum_{c=1}^{M}\sum_{d=1}^{M}\mathbb{E}[G_{ij}G_{ia}G_{ib}G_{ic}Z_{qj}Z_{qd}]\mathbb{E}[\theta_a\beta_{Ib}\theta_c\gamma_{Gd}]
\end{aligned}
$$

Then,

$$
\mathbb{E}[G_{ij}G_{ia}G_{ib}G_{ic}Z_{qj}Z_{qd}] = \begin{cases}\mathbb{E}[G_{ij}G_{ia}G_{ib}G_{ic}]\mathbb{E}[Z_{qj}Z_{qd}] & \text{if } i\in[N_I]\backslash[N_s] \text{ or } q\in[N_G]\backslash[N_s], \\ \mathbb{E}[G_{ij}^2G_{ia}G_{ib}G_{ic}G_{id}] & \text{if } i\in[N_s] \text{ and } q\in[N_s]\end{cases}
$$

By Isserlis' theorem, we have

$$
\begin{aligned}
\mathbb{E}[G_{ij}G_{ia}G_{ib}G_{ic}]\mathbb{E}[Z_{qj}Z_{qd}] &= (\mathbb{E}[G_{ij}G_{ia}]\mathbb{E}[G_{ib}G_{ic}]+\mathbb{E}[G_{ij}G_{ib}]\mathbb{E}[G_{ia}G_{ic}]+\mathbb{E}[G_{ia}G_{ib}]\mathbb{E}[G_{ij}G_{ic}])\mathbb{E}[Z_{qj}Z_{qd}] \\
&= (r_{ja}r_{bc}+r_{jb}r_{ac}+r_{ab}r_{jc}])r_{jd}
\end{aligned}
$$

and

$$
\mathbb{E}[\theta_a\beta_{Ib}\theta_c\gamma_{Gd}] = \mathbb{E}[\theta_a\beta_{Ib}]\mathbb{E}[\theta_c\gamma_{Gd}]+\mathbb{E}[\theta_a\theta_c]\mathbb{E}[\beta_{Ib}\gamma_{Gd}]+\mathbb{E}[\theta_a\gamma_{Gd}]\mathbb{E}[\beta_{Ib}\theta_c]
$$

Then,

$$\mathbb{E}[\theta_a\beta_{Ib}\theta_c\gamma_{Gd}] = \begin{cases} \mathbb{E}[\theta_a\beta_{Ia}]\mathbb{E}[\theta_c\gamma_{Gc}] & \text{if } a = b \neq c = d, \\ \mathbb{E}[\theta_a\theta_a]\mathbb{E}[\beta_{Ib}\gamma_{Gb}] & \text{if } a = c \neq b = d, \\ \mathbb{E}[\theta_a\gamma_{Ga}]\mathbb{E}[\beta_{Ib}\theta_b] & \text{if } a = d \neq b = c \\ 2\mathbb{E}[\theta_a\beta_{Ia}]\mathbb{E}[\theta_a\gamma_{Ga}] + \mathbb{E}[\theta_a\theta_a]\mathbb{E}[\beta_{Ia}\gamma_{Ga}] & \text{if } a = b = c = d \end{cases}$$

If $i \in [N_I] \setminus [N_s]$ or $q \in [N_G] \setminus [N_s]$:

$$\begin{aligned} &\sum_{a=1}^{M}\sum_{b=1}^{M}\sum_{c=1}^{M}\sum_{d=1}^{M}\mathbb{E}[G_{ij}G_{ia}G_{ib}G_{ic}Z_{qj}Z_{qd}]\mathbb{E}[\theta_a\beta_{Ib}\theta_c\gamma_{Gd}] \\ &\approx \sum_{a=1}^{M}\sum_{c=1}^{M}(2r_{ja}r_{ac} + r_{jc})r_{jc}\mathbb{E}[\theta_a\beta_{Ia}]\mathbb{E}[\theta_c\gamma_{Gc}] + \sum_{a=1}^{M}\sum_{b=1}^{M}(r_{ja} + 2r_{jb}r_{ab})r_{ja}\mathbb{E}[\theta_a\gamma_{Ga}]\mathbb{E}[\beta_{Ib}\theta_b] \\ &\quad + \sum_{a=1}^{M}\sum_{b=1}^{M}(r_{jb} + 2r_{ja}r_{ab})r_{jb}\mathbb{E}[\theta_a\theta_a]\mathbb{E}[\beta_{Ib}\gamma_{Gb}] \\ &= (\sigma_E^2\rho_{G2,I} + 2M\mathbb{E}[\theta_a\beta_{Ia}]\mathbb{E}[\theta_a\gamma_{Ga}])\ell_j \end{aligned}$$

If $i \in [N_s]$ and $q \in [N_s]$:

$$\begin{aligned} &\sum_{a=1}^{M}\sum_{b=1}^{M}\sum_{c=1}^{M}\sum_{d=1}^{M}\mathbb{E}[G_{ij}G_{ia}G_{ib}G_{ic}Z_{qj}Z_{qd}]\mathbb{E}[\theta_a\beta_{Ib}\theta_c\gamma_{Gd}] \\ &\approx (\sigma_E^2\rho_{G2,I} + 2M\mathbb{E}[\theta_a\beta_{Ia}]\mathbb{E}[\theta_a\gamma_{Ga}])\ell_j + \sigma_E^2\rho_{G2,I} + 2M\mathbb{E}[\theta_a\beta_{Ia}]\mathbb{E}[\theta_a\gamma_{Ga}] \end{aligned}$$

Thus,

$$\mathbb{E}[T_2] = N_I N_G(\sigma_E^2\rho_{G2,I} + 2M\mathbb{E}[\theta_a\beta_{Ia}]\mathbb{E}[\theta_a\gamma_{Ga}])\ell_j + N_s(\sigma_E^2\rho_{G2,I} + 2M\mathbb{E}[\theta_a\beta_{Ia}]\mathbb{E}[\theta_a\gamma_{Ga}]) \tag{11}$$

For the third term, with a similar calculation for the second term above, we have

$$\begin{aligned} \mathbb{E}[T_3] &= \mathbb{E}[\text{diag}(\boldsymbol{G}\boldsymbol{\theta}_G)(\boldsymbol{G})_j^\top\mathbb{E}[\text{diag}(\boldsymbol{G}\boldsymbol{\theta}_G)\boldsymbol{G}\boldsymbol{\beta}_I]\boldsymbol{\gamma}_G^\top\boldsymbol{Z}^\top\boldsymbol{Z}_j] \\ &= M\mathbb{E}[\theta_a\beta_{Ia}]\mathbb{E}[(\text{diag}(\boldsymbol{G}\boldsymbol{\theta}_G)\boldsymbol{G})_j^\top\boldsymbol{\gamma}_G^\top\boldsymbol{Z}^\top\boldsymbol{Z}_j] \\ &= N_I N_G M\mathbb{E}[\theta_a\beta_{Ia}]\mathbb{E}[\theta_a\gamma_{Ga}]\ell_j + N_s\mathbb{E}[\theta_a\beta_{Ia}]\mathbb{E}[\theta_a\gamma_{Ga}] \end{aligned}$$

Plug in these three terms, adding the correction factor back, and with a similar derivation for the error terms in the G-E independence case, we have the PIGEON-LDSC slope is

$$\frac{\sqrt{N_I N_G}(\rho_{G2,I} + \rho_{G2,E}\rho_{IE})}{CM}\ell_j$$

□

##### 12.13 Proof of Proposition 12

*Proof.* The PIGEON-rGE model (8) is

$$\begin{aligned}
\boldsymbol{Y}_1 &= \boldsymbol{\mu} + \boldsymbol{G}\boldsymbol{\beta}_G + \boldsymbol{V}\boldsymbol{\beta}_{\boldsymbol{E}} + \boldsymbol{V}\boldsymbol{G}\boldsymbol{\beta}_I + \boldsymbol{\epsilon}_0 + \boldsymbol{V}\boldsymbol{\epsilon}_1 \\
\boldsymbol{V} &= \text{diag}(\boldsymbol{v}) \\
\boldsymbol{v} &= \boldsymbol{G}\boldsymbol{\theta}_G + \sqrt{(1-\sigma_E^2)}\boldsymbol{e}
\end{aligned} \tag{12}$$

We can rewrite $\boldsymbol{V}$ as $\boldsymbol{V} = \text{diag}(\boldsymbol{G}\boldsymbol{\theta}_G) + \sqrt{(1-\sigma_E^2)}\text{diag}(\boldsymbol{e})$. We have the Z-score for SNPxE interaction have the moment condition:

$$\begin{aligned}
\mathbb{E}[\boldsymbol{z}_I|\boldsymbol{\beta}_G, \boldsymbol{\beta}_{\boldsymbol{E}}, \boldsymbol{\beta}_I] &= \frac{1}{C^2}[\frac{\boldsymbol{G}^T\boldsymbol{V}\boldsymbol{G}\boldsymbol{\beta}_G}{\sqrt{N_I}} + \frac{\boldsymbol{G}^T\boldsymbol{V}\boldsymbol{V}\boldsymbol{\beta}_E}{\sqrt{N_I}} + \frac{\boldsymbol{G}^T\boldsymbol{V}\boldsymbol{V}\boldsymbol{G}\boldsymbol{\beta}_I}{\sqrt{N_I}}] \\
&= \frac{1}{C^2}[\frac{\boldsymbol{G}^T\sqrt{(1-\sigma_E^2)}\boldsymbol{V}\boldsymbol{G}\boldsymbol{\beta}_G}{\sqrt{N_I}} + \frac{\boldsymbol{G}^T\sqrt{(1-\sigma_E^2)}\boldsymbol{V}\sqrt{(1-\sigma_E^2)}\boldsymbol{V}\boldsymbol{\beta}_E}{\sqrt{N_I}} + \frac{\boldsymbol{G}^T\sqrt{(1-\sigma_E^2)}\boldsymbol{V}\sqrt{(1-\sigma_E^2)}\boldsymbol{V}\boldsymbol{G}\boldsymbol{\beta}_I}{\sqrt{N_I}} + \\
&\quad \frac{\boldsymbol{G}^T\text{diag}(\boldsymbol{G}\boldsymbol{\theta}_G)\boldsymbol{G}\boldsymbol{\beta}_G}{\sqrt{N_I}} + \frac{\boldsymbol{G}^T\text{diag}(\boldsymbol{G}\boldsymbol{\theta}_G)\text{diag}(\boldsymbol{G}\boldsymbol{\theta}_G)\boldsymbol{\beta}_E}{\sqrt{N_I}} + \frac{\boldsymbol{G}^T\text{diag}(\boldsymbol{G}\boldsymbol{\theta}_G)\text{diag}(\boldsymbol{G}\boldsymbol{\theta}_G)\boldsymbol{G}\boldsymbol{\beta}_I}{\sqrt{N_I}}] \\
&= \frac{1}{C^2}[A+B],
\end{aligned}$$

where $A = \frac{\boldsymbol{G}^T\sqrt{(1-\sigma_E^2)}\boldsymbol{V}\boldsymbol{G}\boldsymbol{\beta}_G}{\sqrt{N_I}} + \frac{\boldsymbol{G}^T\sqrt{(1-\sigma_E^2)}\boldsymbol{V}\sqrt{(1-\sigma_E^2)}\boldsymbol{V}\boldsymbol{\beta}_E}{\sqrt{N_I}} + \frac{\boldsymbol{G}^T\sqrt{(1-\sigma_E^2)}\boldsymbol{V}\sqrt{(1-\sigma_E^2)}\boldsymbol{V}\boldsymbol{G}\boldsymbol{\beta}_I}{\sqrt{N_I}}$
and $B = \frac{\boldsymbol{G}^T\text{diag}(\boldsymbol{G}\boldsymbol{\theta}_G)\boldsymbol{G}\boldsymbol{\beta}_G}{\sqrt{N_I}} + \frac{\boldsymbol{G}^T\text{diag}(\boldsymbol{G}\boldsymbol{\theta}_G)\text{diag}(\boldsymbol{G}\boldsymbol{\theta}_G)\boldsymbol{\beta}_E}{\sqrt{N_I}} + \frac{\boldsymbol{G}^T\text{diag}(\boldsymbol{G}\boldsymbol{\theta}_G)\text{diag}(\boldsymbol{G}\boldsymbol{\theta}_G)\boldsymbol{G}\boldsymbol{\beta}_I}{\sqrt{N_I}}$.
Also, we have

$$\text{Var}[\boldsymbol{z}_I|\boldsymbol{\beta}_G, \boldsymbol{\beta}_{\boldsymbol{E}}, \boldsymbol{\beta}_I] = \frac{\sigma_{\epsilon_0}^2\boldsymbol{G}^T\boldsymbol{V}\boldsymbol{V}\boldsymbol{G}}{N_I} + \frac{\sigma_{\epsilon_1}^2\boldsymbol{G}^T\boldsymbol{V}\boldsymbol{V}\boldsymbol{V}\boldsymbol{V}\boldsymbol{G}}{N_I}$$

We denote

$$\begin{aligned}
A_j &= \frac{\boldsymbol{G}_{j\cdot}^T\sqrt{(1-\sigma_E^2)}\boldsymbol{V}\boldsymbol{G}\boldsymbol{\beta}_G}{\sqrt{N_I}} + \frac{\boldsymbol{G}_{j\cdot}^T\sqrt{(1-\sigma_E^2)}\boldsymbol{V}\sqrt{(1-\sigma_E^2)}\boldsymbol{V}\boldsymbol{\beta}_E}{\sqrt{N_I}} + \frac{\boldsymbol{G}_{j\cdot}^T\sqrt{(1-\sigma_E^2)}\boldsymbol{V}\sqrt{(1-\sigma_E^2)}\boldsymbol{V}\boldsymbol{G}\boldsymbol{\beta}_I}{\sqrt{N_I}} \\
B_j &= \frac{\boldsymbol{G}_{j\cdot}^T\text{diag}(\boldsymbol{G}\boldsymbol{\theta}_G)\boldsymbol{G}\boldsymbol{\beta}_G}{\sqrt{N_I}} + \frac{\boldsymbol{G}_{j\cdot}^T\text{diag}(\boldsymbol{G}\boldsymbol{\theta}_G)\text{diag}(\boldsymbol{G}\boldsymbol{\theta}_G)\boldsymbol{\beta}_E}{\sqrt{N_I}} + \frac{\boldsymbol{G}_{j\cdot}^T\text{diag}(\boldsymbol{G}\boldsymbol{\theta}_G)\text{diag}(\boldsymbol{G}\boldsymbol{\theta}_G)\boldsymbol{G}\boldsymbol{\beta}_I}{\sqrt{N_I}}.
\end{aligned}$$

Therefore, by the law of total expectation, we have the expectation of the squared SNPxE Z-score for j-th SNPs is

$$\begin{aligned}
\mathbb{E}[z_{Ij}^2] &= \mathbb{E}[\mathbb{E}[z_{Ij}^2|\boldsymbol{\beta}_G, \boldsymbol{\beta}_{\boldsymbol{E}}, \boldsymbol{\beta}_I]] = \mathbb{E}[\mathbb{E}[z_{Ij}|\boldsymbol{\beta}_G, \boldsymbol{\beta}_{\boldsymbol{E}}, \boldsymbol{\beta}_I]^2 + \text{Var}[z_{Ij}|\boldsymbol{\beta}_G, \boldsymbol{\beta}_{\boldsymbol{E}}, \boldsymbol{\beta}_I]] \\
&= \mathbb{E}[(A_j+B_j)^2 + \frac{\sigma_{\epsilon_0}^2\boldsymbol{G}_{j\cdot}^T\boldsymbol{V}\boldsymbol{V}\boldsymbol{G}_{j\cdot}}{N_I} + \frac{\sigma_{\epsilon_1}^2\boldsymbol{G}_{j\cdot}^T\boldsymbol{V}\boldsymbol{V}\boldsymbol{V}\boldsymbol{V}\boldsymbol{G}_{j\cdot}}{N_I}] \\
&= \mathbb{E}[A_j^2 + B_j^2 + 2A_jB_j + \frac{\sigma_{\epsilon_0}^2\boldsymbol{G}_{j\cdot}^T\boldsymbol{V}\boldsymbol{V}\boldsymbol{G}_{j\cdot}}{N_I} + \frac{\sigma_{\epsilon_1}^2\boldsymbol{G}_{j\cdot}^T\boldsymbol{V}\boldsymbol{V}\boldsymbol{V}\boldsymbol{V}\boldsymbol{G}_{j\cdot}}{N_I}] \\
&= \mathbb{E}[A_j^2 + \frac{\sigma_{\epsilon_0}^2\boldsymbol{G}_{j\cdot}^T\boldsymbol{V}\boldsymbol{V}\boldsymbol{G}_{j\cdot}}{N_I} + \frac{\sigma_{\epsilon_1}^2\boldsymbol{G}_{j\cdot}^T\boldsymbol{V}\boldsymbol{V}\boldsymbol{V}\boldsymbol{V}\boldsymbol{G}_{j\cdot}}{N_I}] + \mathbb{E}[B_j^2] + \mathbb{E}[2A_jB_j]
\end{aligned}$$

With Proof of Proposition 4, we have the first term

$$\mathbb{E}[A_j^2 + \frac{\sigma_{\epsilon_0}^2 \boldsymbol{G}_{j\cdot}^T \boldsymbol{V}\boldsymbol{V}\boldsymbol{G}_{j\cdot}}{N_I} + \frac{\sigma_{\epsilon_1}^2 \boldsymbol{G}_{j\cdot}^T \boldsymbol{V}\boldsymbol{V}\boldsymbol{V}\boldsymbol{V}\boldsymbol{G}_{j\cdot}}{N_I}]$$
$$= (1-\sigma_E^2)(\frac{\sigma_g^2 M}{M} + \beta_E^2 \mu_E(4) + \frac{\sigma_I^2}{M}[\frac{(N_I^2 - N_I)}{N_I}\sum_k^M r_{jk}^2 + \frac{N_I(\sum_k^M 2r_{jk}^2 + M)\mu_E(4)}{N_I}]) + \sigma_{\epsilon_0}^2 + \mu_E(4)\sigma_{\epsilon_1}^2$$

where $\ell_j = \sum_k^M r_{jk}^2$ is the LD score for j-th SNP. Follow by the above proof for oracle PGSxE (11), the summation of the second and the third term is approximately (by ignoring higher order terms) to

$$\mathbb{E}[B_j^2] + \mathbb{E}[2A_jB_j] \approx \sigma_E^2\sigma_g^2 + \sigma_E^2\beta_E^2\mu_E(4) + \sigma_E^2\sigma_I^2 + \sigma_E^2 2\rho_{IE}^2\frac{\sum_k^M r_{jk}^2 N_I}{M} + (\sigma_E^2)^2\sigma_I^2\frac{\sum_k^M r_{jk}^2 N_I}{M} +$$
$$2(1-\sigma_E^2)[\sigma_E^2\sigma_I^2 + \rho_{IE}^2]\frac{\sum_k^M r_{jk}^2 N_I}{M}$$

Merging these terms, we see that the slope in the LDSC is $\frac{(\sigma_I^2 + 2\rho_{IE}^2)N_I}{C^2 M}$. □

#### 12.14 Proof of Proposition 13

*Proof.* We consider the PGSxE regression

$$Y_i \sim \alpha_G^{(asc)} PGS_i^{(asc)} + \alpha_E^{(asc)} E_i + \alpha_I^{(asc)} PGS_i^{(asc)} E_i.$$

Seince $PGS_i^{(asc)}$ and $E_i$ are centered and uncorrelated, we have

$$\begin{aligned}
\alpha_I^{(asc)} &= \frac{\mathrm{Cov}(Y_{i1}, PGS_i^{(asc)} E_i)}{\mathrm{Var}(PGS_i^{(asc)} E_i)} \\
&= \frac{\mathrm{Cov}(\boldsymbol{G}_i\boldsymbol{\beta}_G + E_i\beta_E + \boldsymbol{G}_i * E_i\boldsymbol{\beta}_I + \epsilon_{0i} + \epsilon_{1i}E_i, \boldsymbol{G}_i(\boldsymbol{\beta}_G + \theta\boldsymbol{\beta}_I)E_i))}{\mathbb{E}[E_i]^2 \mathrm{Var}(PGS_i^{(asc)})] + \mathbb{E}[PGS_i^{(asc)})]^2 \mathrm{Var}(E_i) + \mathrm{Var}(PGS_i^{(asc)})] \mathrm{Var}(E_i)} \\
&= \frac{\mathbb{E}[\boldsymbol{G}_i * E_i\boldsymbol{\beta}_I]\mathbb{E}[\boldsymbol{G}_i\boldsymbol{\beta}_G E_i + \theta\boldsymbol{G}_i\boldsymbol{\beta}_I E_i]}{\mathrm{Var}(PGS^{(asc)})} \\
&= \frac{M(\mathbb{E}[\boldsymbol{\beta}_I\boldsymbol{\beta}_G] + \theta\mathbb{E}[\boldsymbol{\beta}_I\boldsymbol{\beta}_I]}{\mathrm{Var}(PGS^{(asc)})} \\
&= \frac{\rho_{GI} + \theta\sigma_I^2}{\sigma_G^2 + \theta^2\sigma_I^2 + 2\theta\rho_{GI}}
\end{aligned}$$

□
